# Synchronous L1 retrotransposition events promote chromosomal crossover early in human tumorigenesis

**DOI:** 10.1101/2024.08.27.596794

**Authors:** Sonia Zumalave, Martin Santamarina, Nuria P. Espasandín, Daniel Garcia-Souto, Javier Temes, Toby M. Baker, Ana Pequeño-Valtierra, Iago Otero, Jorge Rodríguez-Castro, Ana Oitabén, Eva G. Álvarez, Paula Otero, Iria Díaz-Arias, Mónica Martínez-Fernández, Peter Van Loo, Gael Cristofari, Bernardo Rodriguez-Martin, Jose M. C. Tubio

**Affiliations:** Mobile Genomes, Centre for Research in Molecular Medicine and Chronic Diseases (CIMUS), Universidade de Santiago de Compostela – IDIS, Santiago de Compostela, Spain; Instituto de Investigaciones Sanitarias de Santiago de Compostela (IDIS), Santiago de Compostela, Spain; Department of Zoology, Genetics and Physical Anthropology, Universidade de Santiago de Compostela, Santiago de Compostela, Spain; The Francis Crick Institute, London, UK; Translational Oncology Research Group, Galicia Sur Health Research Institute (IIS Galicia Sur), SERGAS-UVIGO, Vigo, Spain; Department of Genetics, The University of Texas MD Anderson Cancer Center, Houston, Texas, USA; Department of Genomic Medicine, The University of Texas MD Anderson Cancer Center, Houston, Texas, USA; University Cote d’Azur, INSERM, CNRS, Institute for Research on Cancer and Aging of Nice (IRCAN), Nice, France; European Molecular Biology Laboratory (EMBL), Genome Biology Unit, Heidelberg, Germany; Centre for Genomic Regulation (CRG), The Barcelona Institute of Science and Technology, Barcelona, Spain; Universitat Pompeu Fabra (UPF), Barcelona, Spain

## Abstract

L1 retrotransposition is a significant source of genomic variation in human epithelial tumours, which can contribute to tumorigenesis. However, fundamental questions about the causes and consequences of L1 activity in cancer genomes remain unresolved, primarily due to the limitations of short-read sequencing technologies. Here, we employ multiplatform sequencing, with an emphasis on long reads, to analyse a fine selection of 10 tumours exhibiting high rates of somatic retrotransposition, encompassing over 6000 events. The analysis of L1 locus-specific single-nucleotide variants reveals a novel panorama of L1 loci activity. Furthermore, examination of the internal structure of somatic L1s uncovers the mechanisms behind their inactivation. A hidden landscape of chromosomal aberrations emerges in the light of long reads, where reciprocal translocations mediated by L1 insertion represent frequent events. Resolution of L1 bridges’ configuration elucidates the mechanisms of their formation, where typically two independent, but synchronous, somatic L1 insertions drive the reciprocal exchange between non-homologous chromosomes. Timing analyses indicate that L1 retrotransposition is an early driver of chromosomal instability, active before the first whole-genome doubling event. Overall, these findings highlight L1 activity as a more significant contributor to tumour genome plasticity than previously recognized, extending its impact beyond simple insertional mutagenesis.

---

L1 retrotransposons are widespread repetitive elements in the human genome, representing ∼17% of the total nuclear DNA content ^1^. These ∼6 kb-long elements behave as intragenomic parasites that copy and paste themselves elsewhere in the genome by a mechanism known as retrotransposition. This process requires the transcription of a source element into an intermediary messenger RNA, which is later reverse transcribed into a DNA molecule during integration of the derived copy ^2,3^. Most of the approximately half-a-million copies of L1 in the human reference genome represent 5’-truncated, inactive elements incapable of retrotransposition. However, a small subset of 150-200 L1 loci exhibits evidence of ongoing activity within the human genome, acting as source elements for further retrotransposition. A handful of these source elements, termed “hot L1s”, represent highly active loci with the ability to generate a significant fraction of derived copies ^4–8^. Insertional mutagenesis by L1 retrotransposons in the human genome is a recognized cause of disease including cancer ^9–12^.

L1 retrotranspositions can drive tumorigenesis ^13–16^. Somatic L1 insertions are detected in about 35% of all human tumours ^10^, being a common feature in oesophageal adenocarcinoma (ESAD), head-and-neck squamous carcinoma (HNSC), lung squamous carcinoma (LUSC), and colorectal adenocarcinoma (COAD). Notably, these four tumour types alone account for 70% of all somatic retrotransposition events detected in human tumours ^10^. Major restructuring of cancer genomes, characterized by large chromosomal deletions, can sometimes arise from aberrant L1 insertion events in tumours with high retrotransposition rates. Occasionally, L1-mediated deletions can promote the loss of megabase-scale regions of a chromosome and drive oncogenesis ^10^. It is likely that genomic structural variants induced by somatic retrotransposition in human cancer are more frequent than could unambiguously be characterized in previous cancer retrotransposition studies, given the constraints on the fragment sizes of paired-end sequencing libraries. Long read sequencing technologies should be able to provide a more comprehensive picture of how frequent such events are, and their role in human tumorigenesis. Thus, here, we employ multiplatform sequencing with an emphasis on long reads, to analyse retrotransposition in an exquisite selection of 10 tumours with high retrotransposition rates, encompassing over 6000 somatic insertions. By introducing novel algorithms, we characterize this mutagenic process to an unprecedented resolution, providing a comprehensive view of retrotransposition dynamics in cancer with a focus on the patterns and mechanisms of L1-mediated rearrangements in the cancer genome.

## The landscape of cancer retrotransposition in the light of long reads

We performed shallow whole-genome sequencing with Illumina paired-ends to identify tumours with high rates of somatic retrotransposition in 137 cancer patients: 37 with HNSC, 50 with LUSC and 50 with COAD. This analysis identified a set of 10 tumours with more than 100 somatic retrotransposition events, including five HNSC, four LUSC and one COAD, which were selected for further sequencing with long reads using Oxford Nanopore technologies (ONT), together with their corresponding matched-non-tumoral adjacent tissues (**Fig. 1a; Supplementary Table 1**). This approach resulted in a median coverage of 33.7x and N50 read size of 19.9 kb for the tumours, and of 31.1x and 18.7 kb for the adjacent tissues (**Supplementary Note**; **Supplementary Fig. 1a-d**).

**Figure 1.**
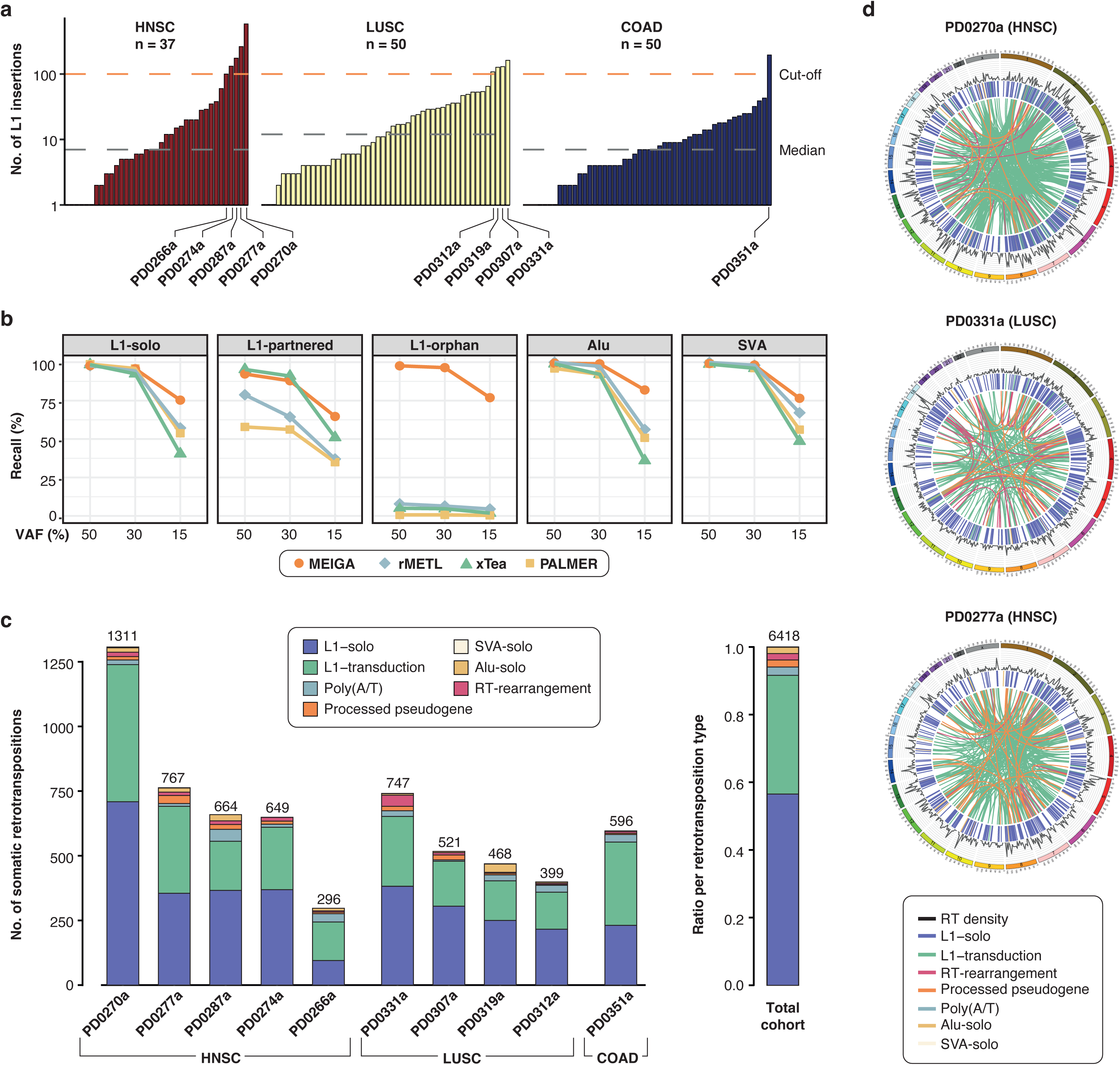
Landscape of somatic retrotransposition in the light of long reads. **(a)** Screening for somatic retrotransposition events in 137 tumours using shallow sequencing with Illumina paired-ends, and selection of 10 tumours showing >100 retrotranspositions insertions for subsequent analysis with long reads. **(b)** MEIGA benchmarking demonstrates high precision and recall for all types of somatic retrotransposition events (see **Supplementary Fig 2c-d**). Here, recall of retrotransposition events at three different variant allele fraction (VAF) levels shows MEIGA performance relative to other relevant long read pipelines, including rMETL, xTea, and PALMER. **(c)** Distribution of the number and type of somatic retrotransposition insertions observed in 10 tumours with high and ultrahigh retrotransposition rates further sequenced with ONT long reads. **(c)** Circos plots showing the landscape of somatic retrotransposition insertions in three relevant tumours; PD0270a shows the highest number of retrotranspositions in the cohort (n=1311); PD0331a bears the highest number of retrotransposon-mediated rearrangements (n=49); and PD0277a shows the highest number of processed pseudogenes (n=31). Note that the coloured links inside the Circos represent mobilizations that connect source elements with their progeny ^7^.

To find and characterize somatic retrotransposition in the long reads’ dataset, we developed a bioinformatics algorithm coined MEIGA (Mobile Element Integration Genome Analyzer, **Supplementary Note; Supplementary Fig. 2a-b**). The method relies on the identification of read clusters indicative of structural variation breakpoints, followed by a reconstruction of such variants through local assembly and the identification of hallmarks of somatic retrotransposition. MEIGA can detect seven main types of somatic retrotranspositions according to the type of sequence retrotransposed. These types are solo retrotranspositions events, which include full or partial L1, Alu or SVA sequences (L1-solo, Alu-solo and SVA-solo, respectively); L1-partnered transductions, in which an L1 sequence along with a piece of sequence downstream are retrotransposed; L1-orphan transductions, in which only the unique sequence downstream of an active L1 is retrotransposed without the cognate L1; processed pseudogenes, where mRNA from nuclear genes are retrotransposed; and solitary polyadenylate [poly(A)] tracts resulting from severely 5’ truncated retrotransposition events. Notably, MEIGA not only detects canonical insertions but also retrotransposition-mediated rearrangements resulting from aberrant integration events.

To evaluate our algorithm, we generated a mock cancer genome into which we seeded retrotransposon insertions of different types and structural features. We then simulated long reads from this mock genome to the standard levels of coverage achieved in out cohort, generating different allele frequency (VAF) levels (0.5, 0.3 and 0.15) of the relevant insertions. Next, we ran MEIGA on this dataset together with three additional long-reads pipelines to detect retrotranspositions. The results confirmed a high performance of MEIGA for all insertion types and tumour clonalities tested (**Fig. 1b**; **Supplementary Fig. 2c-d**; **Supplementary Note**). Additionally, our algorithm successfully reconstructed the size and sequence of the simulated insertions (**Supplementary Fig. 2e**; **Supplementary Note**).

We ran MEIGA on the long-read sequencing data from the 10 relevant tumours and their adjacent tissues. The analysis retrieved a total of 6,418 retrotransposition events acquired somatically and confirmed that all tumours exhibit high rates of somatic retrotransposition (median = 622.5, range [296-1,311]) (**Fig. 1c**; **Supplementary Table 2**). In one remarkable HNSC tumour, PD0270a, we found 1,311 somatic retrotranspositions, accounting for 20% of the total events in the cohort, where all types of insertion are observed (**Fig. 1c-d**). Overall, L1-solos represented the most frequent type of insertions (56%, n = 3,611), followed by L1 transductions (35%, n = 2,240; of which 1,535 are partnered and 705 are orphan), poly(A/T) tracts (2%, n = 157), pseudogenes (2%, n = 133), Alu-solos (2%, n = 119) and SVA-solos (<0.1%, n = 6) (**Fig. 1c**). Additionally, our analysis identified 152 instances of retrotransposon insertion, typically L1s, bridging genomic rearrangements caused by aberrant integration (**Fig. 1c**; **Supplementary Table 3; Supplementary Note**). In a notable LUSC tumour, PD0331a, we identified 49 retrotransposon bridges associated with different types of rearrangements, accounting for one-third (49/152) of the total events (**Fig. 1d**). These non-canonical retrotransposition-mediated structural variants, which represent 2% of the total number of detected somatic retrotranspositions in the cohort (152/6,418), are addressed in a later section.

To validate our results, we performed further sequencing on PD0270a – the tumour with the highest number of retrotranspositions in our cohort (n = 1,290) – and its matched-normal tissue, employing long reads with PacBio Hi-Fidelity and short reads with Illumina paired-ends. Then, we run MEIGA together with two additional retrotransposition pipelines for long reads and short reads (**Supplementary Note**). The comparison of the results from the three independent pipelines indicated that a majority (92.5%, 1,193/1,290) of canonical retrotransposon events called by MEIGA, were consistently identified by at least one additional long or short reads algorithm and/or one additional technology, supporting that they represent genuine events (**Supplementary Fig. 3a**). Additionally, we carried out visual inspection with Integrative Genomic Viewer (IGV) of 150 retrotransposition insertions that represented MEIGA private calls, which supported the somatic acquisition of 142 (true positive rate 94.7%) (**Supplementary Fig. 3b**).

## Patterns and mechanisms of L1 inactivating rearrangements

Previous analyses of germline L1 insertions and de novo retrotransposition events, derived from engineered L1 elements in cultured cells, revealed a variety of L1-inactivating internal rearrangements and their associated mechanisms ^2,3,16–21^. In primary tumours, retrotransposition studies using short reads confirmed that a majority of L1 insertions in cancer represent 5’-truncated elements exhibiting diverse internal structures ^7,14,22–26^. However, the detailed characterization of these internal rearrangements in cancer using long reads has been confined to a limited number of somatic insertions and tumours ^27,28^. Thus, we employed MEIGA to reconstruct and dissect the internal structure of solo-L1 insertions in our long reads’ cohort, aiming to elucidate the mechanisms underlying their inactivation upon integration (note that transductions and pseudogenes are addressed in **Supplementary Note**).

Our analysis shows that the median insertion size of somatic L1-solo insertions is 878 bp, ranging from 55 to 6,022 bp (**Fig. 2a; Supplementary Table 2**). Only a minority fraction (0.3%, 10/3,611) of the solo-L1s are full-length elements with a canonical size and structure (**Fig. 2b; Supplementary Table 2)**, which represent insertions that are likely competent for retrotransposition. Most somatic L1-solos (n=3,598) are non-functional copies bearing a 5’-deletion that truncate the element. Specifically, half of these insertions (49%, 1,762/3,598) show a 5’-deletion-only with no additional internal rearrangements, while the other half (51%, 1,836/3,598) display co-occurrence of 5’-deletion with 5’-inversion. Both groups of truncated insertions arise from distinct mechanisms. The former group results from canonical target primed reverse transcription (TPRT) ^29–35^, which typically generates an L1 insertion with a deletion at the 5’ end ^2^. As for the second group of truncated insertions, the presence of a 5’-inversion represents a hallmark of the twin-priming process, in which two different DNA ends are used as primers (namely, poly(dT) primer and internal primer) on the same L1 RNA template to initiate cDNA synthesis ^3^ (see description in **Fig. 2c, central panel**).

**Figure 2.**
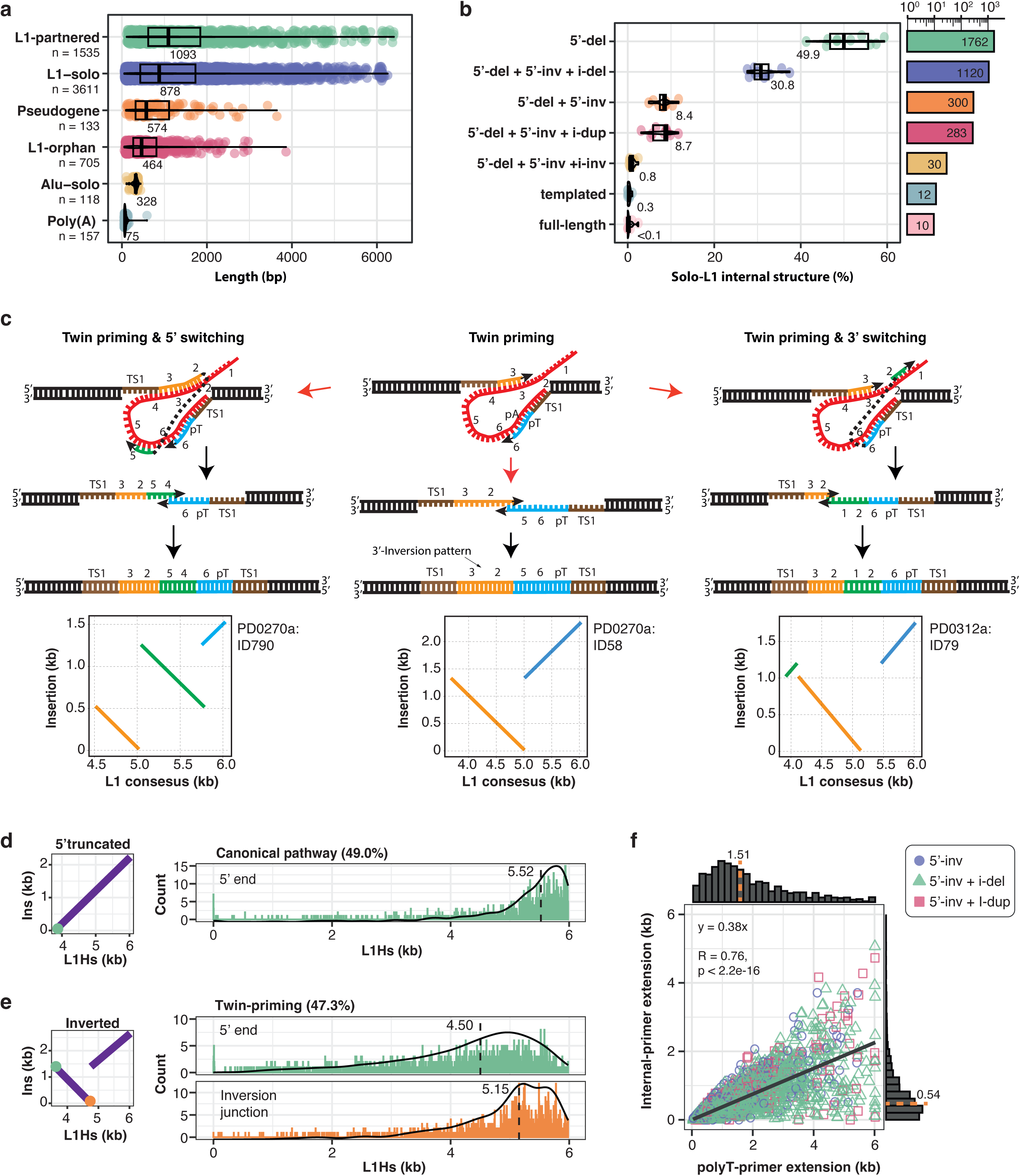
Long reads reveal the internal structure of somatic retrotransposition insertions to unprecedented resolution. **(a)** Box plot showing the insertion length distribution of somatic retrotransposition events across six categories: Alu-solo, L1-solo, L1-partnered transductions, L1-orphan transductions, poly(A/T) insertions and processed pseudogenes. The number of insertions (n) is indicated below each category. The median size for each category is indicated at the corresponding box plot. Given the small number of SVA-solo insertions (n = 6), these were excluded from the analysis. **(b)** Left: Boxplot of the percentage of L1-solo insertions per tumour corresponding to each L1 internal structure category, including: full-length L1 insertions produced by canonical TPRT; 5’-deleted (5’-del) by TPRT; co-occurrence of 5’-deletion and 5’-inversion (5’-del + 5’ inv), co-occurrence of 5’-deletion and 5’-inversion with internal deletion (5’-del + 5’ inv + i-del), co-occurrence of 5’-deletion and 5’-inversion with internal duplication (5’-del + 5’ inv + i-dup), all of them resulting from twin priming; co-occurrence of 5’-deletion and 5’inversion with internal inversion (5’ del + 5’ inv + i-inv) produced by twin priming & switching; and templated insertions. Right: Bars show the total number of insertions detected for each internal rearrangement category in the 10 tumours. Of note, there are 81 out 3,598 insertion that were not classified by our method and were removed from this analysis. **(c)** Mechanisms and associated patterns resulting from twin priming and its variants, namely “twin priming & 5’ switching” and “twin priming and 3’ switching”. In twin priming (centre), both DNA nicks generated at the target insertion site initiate reverse transcription using the same L1 RNA template. While the T-rich DNA extremity anneals at the 3’ poly(A) tail of the L1 RNA, the second DNA end base pairs internally within the L1 RNA, through spurious complementarity, initiating the reverse transcription of a second cDNA molecule. Subsequently, microcomplementarity between the two extending cDNA ends allows second strand DNA synthesis. This process generates a 5’ inversion of the L1 body (orange line in the dot plot), which is typically accompanied by a 5’ deletion (blue line). In the twin priming and 5’ switching variation (left), the extending cDNA primed internally is transferred to an upstream region of the L1 RNA that has not yet undergone reverse transcription and continues extension from that point. In the twin priming and 3’ switching variation (right), it is the cDNA strand extending from the poly(dT) end that is transferred to a region of the L1 RNA downstream of the internally primed cDNA, enabling further extension from that position. Both variations generate an additional internal inversion of the L1 body (green line in the dot plot) with different configuration, in addition to the 5’ inversion typical of twin priming (orange line in the dot plot). Other figure details: L1-RNA in red; cDNA extension from de poly-T primer in blue; cDNA extension from internal primer in orange; cDNA extension upon template switching in green. The dot-plots represent the comparison of the sequence from the L1 bridge reconstructed with ONT (Y-axis) and an L1 canonical insertion (X-axis); colours (blue, orange, and green) in the dot plots match the L1 DNA sequences matching the configuration shown in the models above. **(d)** Distribution of the 5’-breakpoint position along the canonical L1 sequence for 5’-truncated L1-solo insertions, showing a median at position 5.52 kb (dashed line). **(e)** Distribution of the internal breakpoints for L1-solo with a 5’ inversion along L1 canonical sequence. Top: 5’ truncation position (dashed line, median at position 4.50 kb). Bottom: Inversion junction position (dashed line, median at position 5.15 kb). **(f)** Correlation between the lengths of the forward sequence (synthesized from the poly(T) primer) and the inverted sequence (synthesized from the internal primer) in different types of L1 insertions carrying a 5’ inversion generated by twin priming within our cohort.

In our dataset, the breakpoints of 5’-deletions resulting from TPRT show a skewed distribution with median at position of 5.5 kb along the canonical L1 sequence (**Fig. 2d**), while the breakpoints of 5’-deletions and 5’-inversions from insertions following twin priming show distributions with median at positions 4.5 kb and 5.15 kb, respectively (**Fig. 2e**). These results confirm previous findings obtained with long reads in the germline ^20,21^ and in one cancer cell line ^28^ that reported these breakpoints are highly clustered towards the 3’ end of the L1 element. Of note, we find that solo-L1s with 5’-inversions are, overall, three times longer than solo-L1s without inversions (median lengths are 1480 bp and 503 bp, respectively; Wilcoxon rank-sum test, p<0.05; **Supplementary Fig. 4a**), indicating that insertion size is a significantly influenced by the insertion mechanism.

Another distinctive feature of twin priming is the frequent occurrence of internal deletions or duplications at the breakpoint of the 5’-inversion ^3^. An internal deletion arises when the extension from the poly(dT) DNA end fails to reach the internal primer, while an internal duplication results from the progression of the cDNA strand initiated at the poly(dT) end beyond the limits of the internal primer binding site ^3^ (**Supplementary Fig. 4b**). In our cohort, 66% (1,120/1,703) and 17% (283/1,703) of the insertions with a 5’-inversion bear an internal deletion or duplication, respectively (**Fig. 2b**). These events are typically short, with median lengths of 14.5 bp for internal deletions and 22 bp for internal duplications (**Supplementary Fig. 4c**). Overall, we find that the extension of the retrotranscription from the poly(dT) primer is on average 2.6-fold larger than from the internal primer (with median sizes of 1.51 kb and 0.54, respectively; **Fig. 2f**), suggesting sequential rather than simultaneous priming events. Furthermore, we find a strong positive correlation between the lengths of the extension products initiated at each primer (R = 0.76, p-value = 0; **Fig. 2f**), which suggests that twin priming is a tightly regulated process.

Notably, we find that some L1 insertions in our dataset follow distinctive patterns of internal rearrangements that do not completely fit with the canonical insertion models described above, shedding light on alternative insertion mechanisms. First, we observe 20 instances in which an internal inversion is found at the breakpoint of a twin priming-derived 5’-inversion (**Fig. 2c, left panel**). Here, we propose a “twin priming and 5’ switching” mechanism model, in which, always in the context of twin priming, the extending cDNA primed internally is transferred to an upstream region of the L1 RNA that has not yet undergone reverse transcription, and continues extension from that point (**Fig. 2c, left**). Second, we identify three instances where the breakpoint of a twin priming-derived 5’-inversion shows a non-inverted L1 template copied from a region of the L1 RNA that is located far away from the canonical poly(dT) primer (**Fig. 2c, right panel**). This uncommon pattern may be explained by “twin priming & 3’ switching”, in which it is the cDNA strand extending from the poly(dT) end that is transferred to a region of the L1 RNA downstream of the internally primed cDNA, enabling further extension from that position (**Fig. 2c, right**).

Our analysis also identified 22 instances, namely templated insertions, in which non-L1 templates were copied and incorporated to the L1 sequence during the integration process, resulting in chimeric L1s that duplicate a genomic segment of DNA ^2^. These events typically involve templates smaller than 250 bp that are located within 15 bp of the insertion site (**Supplementary Fig. 4d**). Previous work proposed that during L1 reverse transcription, the growing cDNA may disassociate from the L1 RNA and pair with nearby homologous DNA regions in the genome, a process akin to synthesis-dependent strand annealing ^2^. Since our pipelines do not specifically search for such events, we believe they may be more frequent than we show here.

## An extended panorama of source elements activity

During transcription of an L1 source element, a fraction of L1 transcripts extend beyond the L1 canonical polyadenylation signal located at the 3’-end of the element ^36–38^, terminating within the downstream genomic sequence when encountering an alternative polyadenylation site. These extended L1 transcripts, when serving as templates for reverse transcription, result in new insertions carrying a segment of DNA originating from the source locus. This phenomenon, known as 3’ transduction, enables the unambiguous identification of the source L1 element from which they derived ^39,40^. Utilizing this property, previous studies based on paired-end sequencing data have identified 124 active L1 source elements in human cancer ^7,10,41^. Similarly, we employed a transduction-based approach to assess L1 activity in our long-reads cohort, identifying 151 active L1 loci across the 10 tumours (**Supplementary Table 4**).

These analyses, however, assume that the proportion of readthrough transcripts is consistent across all loci, suggesting that the number of 3’ transduction originating from a specific L1 locus reflects its activity, which is not likely the case, leading to an underestimation of the number of active L1 loci. Thus, to mitigate this potential bias and obtain a more comprehensive envision of the activity of L1 retrotransposons in cancer, we leveraged long read data to adopt an alternative approach, which can also use information from L1-solo somatic insertions. Our approach, called Source Inference method, relies on identifying unique combinations of diagnostic nucleotide substitutions specific to each L1 locus, enabling us to trace L1-solos back to the source element whence they derive ^22^ (**Fig. 3a**; see **Supplementary Note** and **Supplementary Fig. 5** for a description of the method and its evaluation).

**Figure 3.**
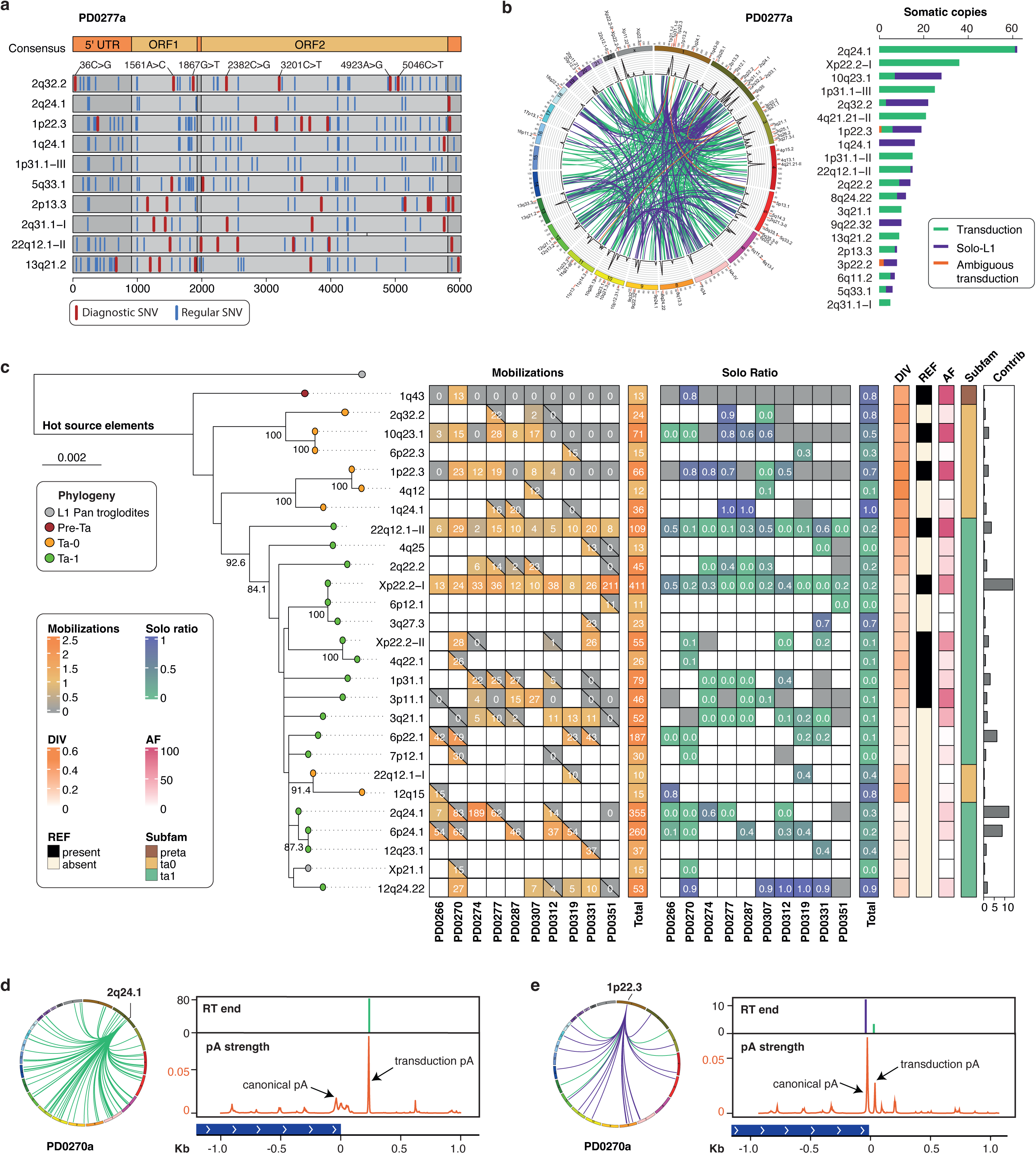
Long reads illuminate a novel panorama of L1 source elements’ activity. **(a)** The diagnostic nucleotide strategy identifies, in each patient, nucleotide substitutions specific to each potential source L1 locus. Then, diagnostic nucleotides are called in the set of somatic solo-L1 events from the tumour, allowing us to distinguish between different potential source elements. For illustrative purposes, we show diagnostic nucleotides (in red) identified in 10 L1 source loci from patient PD0277. **(b)** Activity of L1 source elements detected in tumour PD0277a using diagnostic nucleotides. Left: Circos plot illustrating the source loci of somatic L1 insertions. Green links represent copies traced using a transduction-based strategy, blue links show copies traced using a diagnostic nucleotide-based strategy, and orange links represent ambiguous transductions (i.e., typically short L1-partnered transduction whose element-of-origin could only be identified by diagnostic nucleotides present in the companion L1 sequence); Right: Histogram showing the number of somatic L1 insertions (green for transductions, blue for solo insertions) originating from the twenty most active L1 source elements in tumour PD0277a. **(c)** Phylogeny and properties of hot-L1 source elements in the cohort. We identified 27 hot-L1 sources across the 10 tumours, considering both solo insertions and transduction events. Left: Phylogeny of these hot L1 loci. Right: Heat maps depicting for each source element and each tumour the number of derived copies (mobilizations) and the solo ratio (calculated by dividing the number of solo insertions by the total number of insertions, including transductions). The presence of a box with a diagonal line means that locus is heterozygous for presence/absence of a L1 source element. Other features are shown, including the divergence relative to the consensus L1 sequence (DIV), the genotype (presence or absence) in the genome reference (REF), the allele frequency (AF), the L1 subfamily (pre-Ta, Ta-0, Ta-1), and the overall relative contribution (i.e., number of derived copies from a given element divided by the total number of derived copies for all source elements). **(d-e)** Polyadenylation signal strength determined the activity patterns of source L1 elements active in tumour PD0270a. **(d)** Left: Circos plot showing the segregation of transductions (green links; n = 83) from a source element located at chromosome 2q24.1. Right: This source exclusively propagates through somatic transductions, all ending precisely 234 bp downstream of its 3’ end. Analysis of polyadenylation (pA) signals reveals an absence of the canonical pA signal typically found at the end of L1 elements. However, an alternative pA site was identified 234 bp downstream, exactly where the derived transductions terminate. **(e)** Left: In contrast, the L1 source element at chromosome 1p22.3 primarily generates solo insertions (blue links; n=18) compared to transductions (green links; n=5).. Right: Analysis using the APARENT tool shows a stronger canonical pA signal at the end of this source element compared to an alternative pA site further downstream. Interestingly, this alternative pA site predicted by APARENT coincides with the endpoint of the transductions.

Notably, our Source Inference algorithm successfully attributed 899 somatic L1-solo insertions to specific donor elements, which represents a significant fraction of L1 retrotranspositions that is inaccessible to transduction-based approaches. This had a substantial impact on the patterns of source element activity. In addition to the 151 source elements identified in the cohort by their derived transductions, another set of 46 elements emerged as donors of somatic solo-L1s using internal SNVs (**Supplementary Table 4**; **Supplementary Fig. 6a-b**), bringing the total number of active source elements in the cohort to 197. This includes 117 previously unreported sources in the Pan-Cancer Analysis of Whole Genomes (PCAWG) project ^42^. For instance, in tumour PD0277a, the diagnostic nucleotides strategy identifies up to 12 active sources that were missing in the transduction-based analysis, with three of them (1q24.1, 9q22.32, 3p22.2) ranking within the top 20 most active elements in this tumour (**Fig. 3b**). Furthermore, some source elements, already represented in the transduction analysis, considerably increased their activity when derived solos were included. For instance, in PD0277a, the source at 2q32.2 increased its activity from three transductions to 22 derived insertions, after adding 19 derived solos, elevating this source from the twentieth to the fifth position in the activity ranking (**Fig. 3b**). Overall, in 37% (73/197) of the source elements from our cohort, the number of assigned solos exceeds the number of assigned transductions (**Supplementary Table 4**; **Supplementary Fig. 6a-b**).

In view of these results, we aimed to identify and characterize the mobilization of the most active L1 loci in our cohort, known as “hot-L1s” ^4,5,7^ (details in **Supplementary Note**; **Supplementary Fig. 6c**). Our analysis identified 27 hot-L1 sources (**Fig. 3c**), including 18 elements from the youngest Ta-1 subfamily, eight from the Ta-0 subfamily, and one from the ancient pre-Ta subfamily. Remarkably, two hot source elements, located at Xp22.2-I and 2q24.1, exhibited exceptional activity, giving rise to 411 and 355 somatic insertions, respectively (**Fig. 3c**; **Supplementary Table 4**), accounting for 12% (766/6266) of the total somatic canonical retrotranspositions events in the cohort. Notably, derived solos represent 19% (79/411) and 30% (105/355) of the insertions attributed to Xp22.2 and 2q24.1 (**Fig. 3c**). Overall, the inclusion of solos increased the activity of hot-L1s by an average of 1.37-fold compared to a strategy based solely on transductions.

Notably, among hot-L1s, eight elements exhibit a higher number of assigned solos relative to transductions (**Fig. 3c**; **Supplementary Fig. 6a-b**). In one remarkable example, a hot element at 1q24.1 promoted up to 36 solo-L1s, including 16 in tumour PD0277a and 20 in PD0287a, but no transductions at all. Other hot-L1 sources, although supported by transductions, show a substantial increase in activity when considering derived solos. For instance, in the hot source at 12q24.22, solo-L1s represent 92% (49/53) of the total derived copies, ranking this element among the 10 most active progenitor copies in the cohort (**Fig. 3c**). Similarly, 66% (43/65) of the copies derived from a hot element at 1p22.3 are solo-L1s. On the contrary, in the HNSC tumour PD0270a, the L1 source element at 2q24.1 gave rise to 83 derived copies, all carrying transductions and no solo-L1s (**Fig. 3c**; **Supplementary Table 4**). Of note is the hot source at 1q43, an ancient pre-Ta element that likely integrated into the human genome more than 1 million years ago, representing one of the oldest hot-L1s in the cohort. This source is active in tumour P0270a, yielding to 13 derived copies, of which 11 are solo-L1s (**Fig. 3c**).

## Polyadenylation signals determine somatic solo-L1 insertion rates

To elucidate why some L1 loci primarily propagate through insertions carrying transductions, while others preferentially produce solo-L1s, we compared the downstream sequences of two L1 sources with contrasting patterns. Using the neural network APARENT ^43^, we analysed the strength of canonical and alternative polyadenylation sites in these loci. In the source L1 at 2q24.1 from tumour PD0270a, which showed 83 derived transductions and no solos (mentioned above), APARENT predicted a very weak polyadenylation signal at the end of the element. Instead, the first well-supported alternative polyadenylation site was identified 234 bp downstream of the L1 3’ end, coinciding with the termination point of most transductions (**Fig. 3d**). In contrast, the source element at 1p22.3 from the same tumour predominantly produced solo-L1s (n=18) relative to transductions (n=5), with transductions terminating 64 bp downstream of the L1 element. Here, a robust canonical polyadenylation signal was predicted at the end of the L1 source element, followed by a weaker, but still well-supported, polyadenylation site located exactly 64 bp downstream of the element, corresponding to the end of transductions end (**Fig. 3e**). These findings underscore the critical role of canonical and alternative polyadenylation signals in determining the proportion of derived solo-L1s versus transductions.

## DNA methylation profiles of L1 source elements and their progeny

To explore potential factors influencing the activation of source elements, we conducted whole-genome methylation analysis on samples from our cohort. Using nanopore sequencing data, we estimated the average methylation level for each CpG site in both tumoral genomes and their adjacent tissues (**Supplementary Note**). Our analysis shows that tumoral genomes are on average 17.9% less methylated than their non-tumoral counterparts (Mann–Whitney-Wilcoxon test: p = 1.817×10^-4^; **Supplementary Fig. 7a**). Furthermore, the analysis of 269 full-length L1 loci shared among the 10 patients in our cohort reveals that L1 promoters in tumours consistently display lower methylation compared to adjacent tissues (methylation average of 0.413 and 0.792, respectively, for tumours and normal tissues; Mann-Whitney-Wilcoxon test: p < 0.001; **Supplementary Fig. 7b-c**). Notably, analysis of 71 somatically acquired L1 copies that retained either full or partial promoters revealed a similar hypomethylation pattern (methylation average = 0.315; **Supplementary Fig. 7d**), indicating that full-length somatic L1s are poised for mobilization in tumours, a competence that was indeed demonstrated in previous work ^7^. These observations underscore the association between DNA hypomethylation and L1 activity in tumours, including for new somatic copies, though additional layers of epigenetic regulation may also play a role ^7,22,27,44^.

## A hidden landscape of balanced translocations mediated by L1

We previously reported that aberrant somatic L1 retrotransposition can mediate genomic rearrangements of different types and complexity, where genomic deletions were the most common events with only a minority proportion of other rearrangement types ^10^. However, giving the constraints of short-read sequencing technologies, we investigated whether such events could be more frequent than previously detected. Hence, we used MEIGA to explore the patterns and mechanisms of cryptic chromosomal instability mediated by retrotransposons in our long reads’ dataset (**Supplementary Note**). Here, a retrotransposon-mediated rearrangement (RT-RG) is defined as a junction between two distant breakpoints in the genome bridged by a retrotransposition event acquired somatically.

Our algorithm identified a total of 152 instances following such configuration in our cohort (**Supplementary Table 3**). Notably, although deletion junctions still are the most prevalent category (43%, 66/152), we find that interchromosomal bridges represent the second category with one-third (30%, 45/152) of the events, followed by inversions (18%, 27/152) and duplications (9%, 14/152) (**Fig. 4a**). These rearrangement types showed different size distributions (**Supplementary Fig. 8a**). We identify RT-RGs in each one of the 10 tumour samples from our cohort, where they show a considerable inter-patient variability in both number (median = 13.5, range 3 to 49) and type of rearrangement (**Fig. 4b**). Of note is the LUSC tumour PD0331a, bearing 32% (49 out of 152) of all RT-RGs from the cohort and encompassing all rearrangement types (**Fig. 4b**; **Fig. 1d**). Despite solo-L1s represent the most frequent type of bridge (66%, n=101), we observe that other types of retrotransposons can also be involved (**Fig. 4c; Supplementary Table 3**). For instance, our analysis identifies two rearrangements mediated by processed pseudogenes, and one rearrangement mediated by an Alu element. Overall, these results demonstrate that all RT-RG types but deletions were underestimated in previous work and reveal a more complex landscape of L1-mediated rearrangements in cancer than previously observed ^10^.

**Figure 4.**
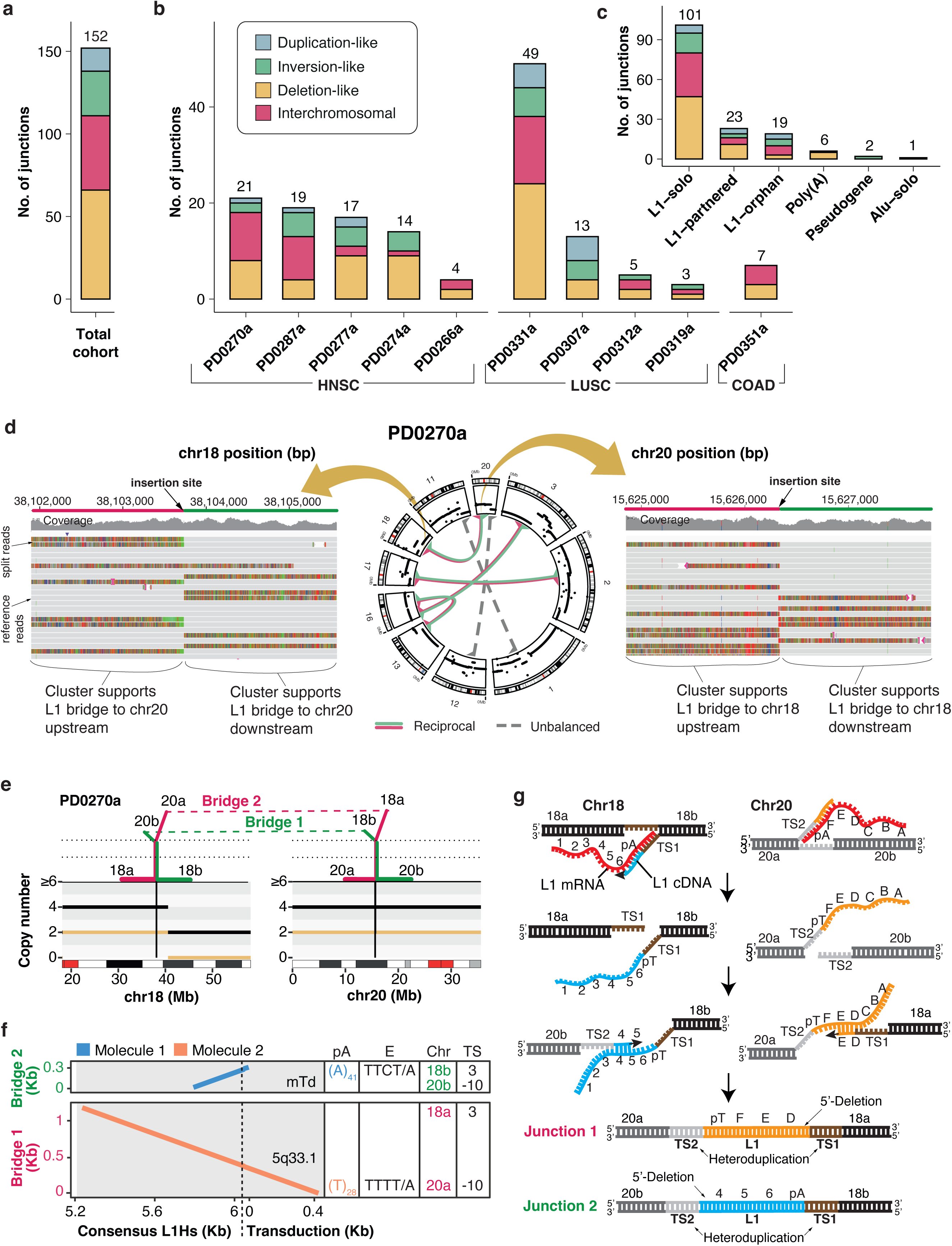
Long reads reveal the landscape of cryptic retrotransposon-mediated rearrangements. **(a)** Proportion of the different types of retrotransposon-mediated rearrangements (n=152) acquired somatically in the 10-tumour cohort sequenced with long reads. Retrotransposition bridges are classified into four categories (see **Supplementary Note**): deletion-like, duplication-like, inversion-like, and interchromosomal rearrangements. **(b)** Histogram showing the distribution of the number and types of somatic retrotransposon-mediated rearrangements per sample observed in the 10-tumour cohort. **(c)** Number and category of retrotransposed sequences at the rearrangement bridges. **(d)** Circos plot (centre) depicting retrotransposon-mediated translocations in tumour PD0270a identified using long reads. Red and green links within the circle represent the L1 bridges with the segment on chromosomes indicating the orientation of the reciprocal translocation junctions. Non-reciprocal (unbalanced) translocations are shown in grey. Integrative Genomics Viewer (IGV) snapshots (left and right) of the long read sequencing data illustrate the breakpoints of the reciprocal translocation between chromosomes 18 and 20 highlight the types of read clusters employed by MEIGA to identify this translocation. **(e)** ReConPlot ^57^ representing a retrotransposon-mediated reciprocal translocation between chromosomes 18q and 20p in tumour PD0270a, which involves two L1 bridges. Chromosome ideograms with positions in megabases (Mb) and cytogenetic bands are shown below (centromere in red). Copy number plots above the ideograms show the minor copy number (yellow line) and the total copy number (black line). On the top, the red and green lines indicate the configuration of the two derivative chromosomes and the bridges (discontinued lines). Here, bridge 2 (red) joins chr18a with chr20a, and bridge 1 (green) joins chr18b with chr20b. **(f)** Structure of the L1 bridges for the translocation shown in (e). Dot plots show the pairwise nucleotide sequence similarity between the L1 bridges (Y-axis) and the consensus L1 sequence (X-axis). For partnered transductions, the bridge sequence is aligned against the transduced sequence in the reference genome. Alignments are read bottom (0 kb) to top, with sequences oriented 5’ to 3’. Colours (blue and orange) indicate the molecular origin of the L1 sequences. Poly(A) tails (pA) and endonuclease motifs (E) are shown at the corresponding breakpoint (bottom for 5’ end; or top for 3’ end). The dot plots revealed the participation of two distinct L1 insertions in the promotion of this translocation. Bridge 2 is exhibits an estimated 41 bp pA tail and TTCT/A endonuclease motif at its 3’ end that is preceded by an L1, and bridge 1 shows a poly(T) tail and TTTT/A endonuclease motif at its 5’ end followed by a distinct L1 insertion. The chromosomal region (Chr) matching the ReConPlot from above is indicated at the corresponding breakpoint (bottom, when present at the 5’ end of the bridge; or top, when present at the 3’ end of the bridge). Here, bridge 1 is demarcated by chr20a and chr18a at the 5’ and 3’ end, respectively; while bridge 2 is demarcated by chr20b and chr18b at the 5’ and 3’ end, respectively. The size of the target site (TS) is indicated in bp at the corresponding breakpoint (bottom, when present at the 5’ end of the bridge; or top, when present at the 3’ end of the bridge). Here, bridge 1 and bridge 2 are demarcated by a heteroduplication of 3 bp and -10 bp. (of note, the last is a target site deletion, which is a fairly frequent event in L1 integration ^2^). **(g)** Mechanism for the reciprocal translocation between chromosomes 18q and 20p, explaining the configuration of the bridges and the genomic rearrangement described in (e-f), involving two insertions following TPRT. Refer to the main text for a detailed description of the mechanism.

Notably, 44% (20/45) of the retrotransposon bridges mediating interchromosomal junctions are involved in mediating 13 reciprocal translocations, a type of chromosomal rearrangement characterized by a balanced exchange of genetic material between two non-homologous chromosomes, which were largely undetected in previous work ^10^ but can now be resolved (**Fig. 4d**). Analysis of the sequencing data revealed that different structural configurations of the junctions are possible: (i) first, a bridge can be conformed of chimeric sequences arising from the recombination of two independent retrotransposition events; (ii) second, a bridge can be conformed of an independent (not recombined) retrotransposition event; (iii) third, in a minority of cases, the bridges of the two derived chromosomes are made of fragments from a single retrotransposition event that was sawed in half. The patterns and mechanisms underlying these outcomes are explored below.

For example, in the HNSC tumour PD0270a we identified a reciprocal translocation between chromosomes 18q and 20p in which the junctions of the rearranged chromosomes exhibit canonical hallmarks of two independent retrotransposition events (**Fig. 4e-f**). Each junction from the two derivative chromosomes shows distinct L1-partnered transductions, followed by its own poly(A) tail and its own L1 endonuclease motif (TTCT/A and TTTT/A). Moreover, the analysis of the target site duplications (TSDs) resulting from these insertion events revealed that each individual L1 insertion is flanked by an heteroduplication, where the two copies of each TSD are located on different non-homologous chromosomes, rather than being on the same chromosome. These hallmarks suggest a retrotransposition-mediated chromosomal exchange requiring spatiotemporal coincidence of two independent L1 insertions (**Fig. 4g**). In this scenario, second strand synthesis is initiated in trans on the second nick of a distinct insertion located on another chromosome, rather than in cis. Alternatively, the observed rearrangement could potentially result from post-integration homologous recombination between two somatic L1 insertions. However, we do not favour a post-integration process as the intermediate state with the two canonical L1 insertions was not supported by the sequencing data.

In another example, the LUSC tumour PD0331a exhibits a reciprocal translocation between chromosomes 22p and 12q, where L1 insertions once again serve as bridges at the breakpoint junctions (**Fig. 5a**). However, here we observe a more complex pattern than the one described above, in which the two L1 bridges are chimeric sequences made of pieces from the two original retrotransposition insertions involved (**Fig. 5b**). Furthermore, the breakpoints at which both retrotranspositions are exchanged match a change in the orientation of the two retrotransposon sequences involved, reminiscent of the 5’-inversion generated by twin priming. Additionally, the bridge in one of the derivative chromosomes shows no poly(A) tail and no endonuclease motif; however, the bridge in the other derivative chromosome presents two poly(A) tails and two endonuclease motifs, which include the poly(A) and the endonuclease motif that were missing in the first bridge. Each L1 bridge is flanked by a target site heteroduplication as described above. Note that the surgical precision of the L1 sutures described here rules out the possibility of any post-integration homologous recombination mechanism and can only be explained by the aberrant resolution of two concomitant twin priming events, as shown in **Fig. 5c**. Again, our findings suggest that synchronicity of two L1 retrotransposon insertions in required to mediate this type of reciprocal translocations. We speculate that such events possibly occur within active biomolecular condensates, related to DNA replication, transcription, or retrotransposition itself ^45^ where DNA molecules from different chromosomes are brought into close proximity.

**Figure 5.**
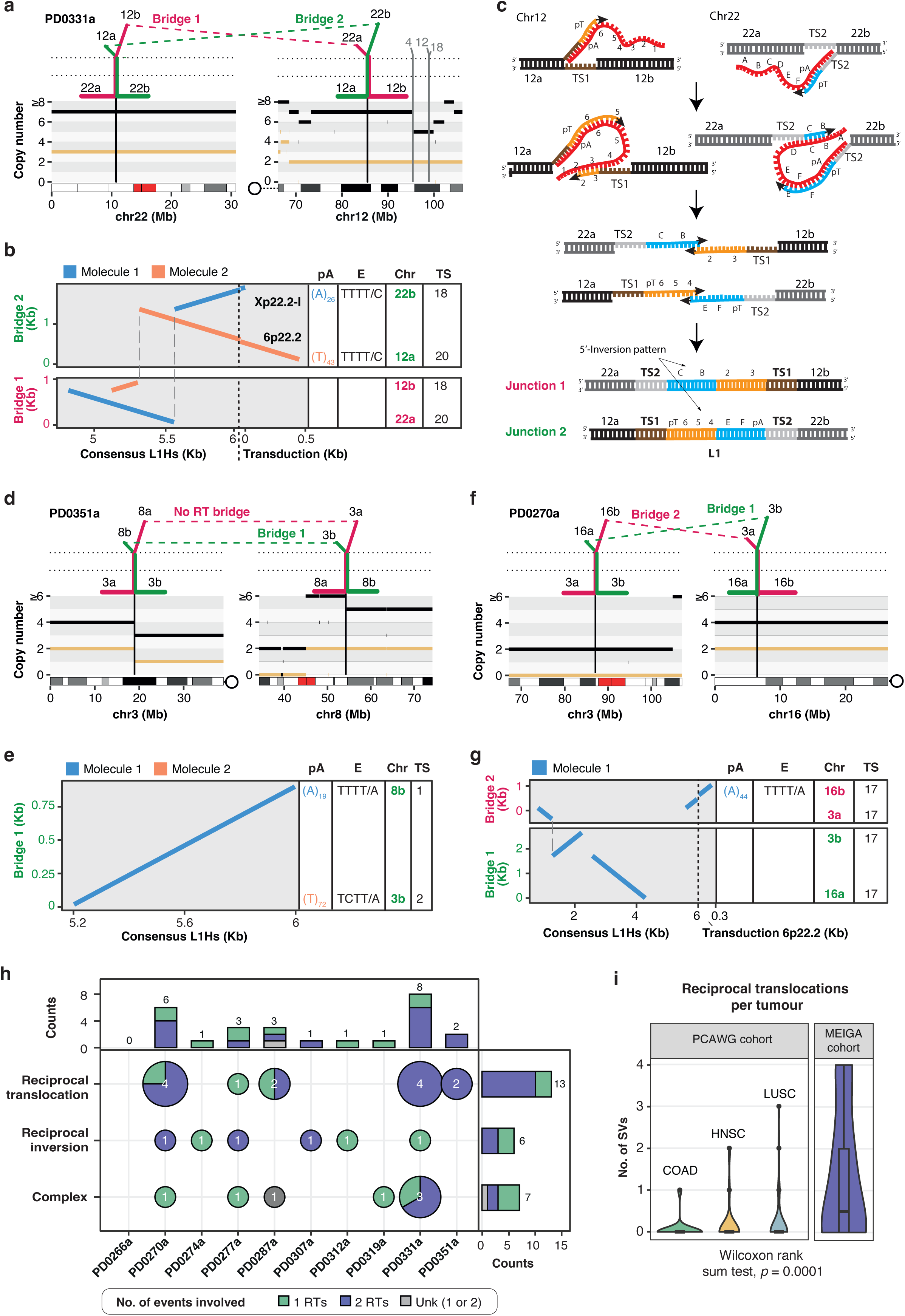
Structural variability of L1 bridges mediating reciprocal translocations. We observe different patterns of L1 bridges suggesting alternative mechanisms for the promotion of L1-mediated reciprocal translocations. **(a)** ReConPlot depicting a reciprocal translocation between chromosomes 22p and 12q in tumour PD0331a, which is mediated by two L1 bridges. Here, bridge 1 joins chr22a with chr12b, while bridge 2 joins chr22b with chr12a. A circle with a dashed line next to the chromosome ideogram marks the relative position of the centromere. **(b)** Structure of the bridges for the translocation shown in (a). Dot plots reveal the presence of two distinct L1 molecules. Bridge 2 exhibits a L1 sequence (made up of two different L1 partnered transductions) that is demarcated by a poly(A) tail and a TTTT/C endonuclease motif at the 3’ end, and a poly(T) tail and a TTTT/C endonuclease motif at its the 5’ end. However, bridge 1 shows a L1 sequence made up of remnants from the two mentioned L1s, with no poly(A/T) tail nor endonuclease motif. Of note, the L1 sequences (blue and orange) involved share two inversion-like breakpoints between bridges 1 and 2 (dashed lines across bridge boxes) that are indicative of twin priming. Furthermore, these bridges are flanked by a heteroduplication of 18 bp and 20 bp. **(c)** Mechanism for the reciprocal translocation between chromosomes 22p and 12q, explaining the configuration of the bridges and the genomic rearrangement described in (a-b), which requires two distinct L1 partnered transductions following twin priming and subsequent recombination between the growing cDNAs. Refer to the main text for a detailed description of the mechanism. **(d)** A reciprocal translocation between chromosomes 3p and 8q in tumour PD0351a. Only bridge 1 has an L1, which joins chr3b with chr8b, the other breakpoint of the translocation (red dashed line) joins chr3a with chr8a. **(e)** Structure of the L1 bridge for the translocation shown in (d). The dot plot reveals a single retrotransposition bridge resulting from two L1 retrotransposition events. The 5’ end of the bridge exhibits a poly(T) tail and TCTT/A endonuclease motif, while its 3’ end has a poly(A) tail and TTTT/A endonuclease motif. Associated mechanisms shown in **Supplementary Fig. 8b**. **(f)** A reciprocal translocation between chromosomes 3p and 16p in tumour PD0270a with two L1 bridges. Bridge 1 joins chr3b with chr16a, while bridge 2 joins chr3a with chr16b. **(g)** Dot plots reveal two bridges generated by a single L1 retrotransposition event. Here, a poly(A) tail together with a TTTT/A endonuclease motif are found at the 3’ of the second L1 bridge, while a single L1 fragment is found at bridge 1. A shared inversion breakpoint (dashed line) indicates for L1 fragments are related and likely arose from one single L1 event. **(h)** Number of structural variants (reciprocal translocations, reciprocal inversions, complex) classified by the number of independent retrotransposition events involved in their formation (1, 2, or unknown). **(i)** Retrotransposon-mediated reciprocal translocations are significantly more frequent than classical reciprocal translocations (i.e., those not involving a somatic retrotransposition) in PCAWG for the same tumour types (HNSC, LUSC, COAD; Wilcoxon rank sum test, p=0.0001121).

In addition to the patterns described above, where two L1 bridges are involved, we also identified reciprocal translocations in which a retrotransposon bridge is exclusively present in one of the two derivative chromosomes. For example, the COAD tumour PD0351a exhibits a reciprocal translocation between chromosomes 3p and 8q where the sequencing data shows a single L1 element inserted in one of the derivative chromosomes, which is flanked by two poly(dA) tails and two endonuclease motifs (**Fig. 5d-e**). However, the translocation junctions at the other derivative chromosome shows no L1 and no retrotransposition hallmarks. These findings suggest two incipient L1 insertion events in 3p and 8q, whose cDNA strands recombined in such a way that left one single L1 body flanked by two poly(dA) tails and two endonuclease motifs in one of the derivative chromosomes only (details in **Supplementary Fig. 8b**).

Finally, our analysis identified reciprocal translocations in which one-single retrotransposition event is involved in mediating both bridges of the derivative chromosomes. For instance, in tumour PD02170a, a reciprocal translocation between chromosomes 3 and 8 shows two L1 bridges but only one endonuclease motif and one poly(dA) tail. Moreover, the sequences of the two L1s share one breakpoint (**Fig. 5f-g**). These hallmarks suggest that both bridges are conformed by one-single retrotransposition event that was sawed in half to mediate the balanced rearrangement.

To validate the occurrence of retrotransposon-mediated reciprocal translocations with orthogonal methods, we selected the lung adenocarcinoma cell-line NCI-H2009, previously identified for its high rates of somatic retrotransposition ^7^. We first sequenced the cell line with ONT, generating long-reads with an average coverage of 25x and N50=16 kb, and subsequently run MEIGA to detect somatic retrotransposition events and associated rearrangements. We identified 23 RT-RGs, including a reciprocal translocation involving chromosomes 3q and 6q with two L1 bridges (**Supplementary Fig. 8c-d**). We then conducted fluorescence in-situ hybridization (FISH) using whole-chromosome probes specific to chromosomes 3 and 6. This approach confirmed, at the cytogenetic level, a reciprocal translocation with the expected configuration between the 3q and the 6q arms (**Supplementary Fig. 8e**). Similarly, chromatin conformation capture by Hi-C in this cell line indicated interchromosomal interactions in the 3D genome between chromosomes 3q and 6q at the expected breakpoints, consistent with a reciprocal translocation (**Supplementary Fig. 8f**).

Overall, our results show that reciprocal translocations can arise from either one or two independent retrotransposition events, although the later seems to be the more frequent outcome (**Fig. 5h**). In 10 out of 13 instances, reciprocal translocations resulted from the aberrant integration of two distinct and synchronous insertions. However, our dataset also contains three reciprocal translocations where a single retrotransposition event was involved. We find that retrotransposon-mediated reciprocal translocations are more frequent in our high-retrotransposition rate tumours than canonical reciprocal translocations are in PCAWG ^46^ (Wilcoxon rank sum test, p<0.01; **Fig. 5i**), revealing somatic retrotransposition as a principal cause of this type of variation in cancer.

## Synchronous L1 insertions can generate reciprocal inversions and complex rearrangements

We also find instances in which one or two independent retrotransposition events, by employing mechanisms analogous to those described above, can be mediating one or two bridges from a same reciprocal inversion (**Fig. 5h**). For instance, in tumour PD0307a, we detected a paracentric inversion spanning 2.67 Mb on chromosome 5p. Analysis of the breakpoint junctions revealed the involvement of two independent retrotransposition events (**Fig. 6a-b)**. One bridge comprises two distinct 5’-truncated L1 elements in a relative inverted orientation and joined internally at their 5’ breakpoints. Each truncated L1 fragment displayed a poly(dA) tail at its 3’ end, with a target site heteroduplication marking the boundaries of the bridge. The second bridge, located megabases away, exhibits a heteroduplication encompassing an interstitial L1 fragment. Noteworthy, the 3’ breakpoint of this fragment matches the 5’ breakpoint of one of the two L1s in the first bridge, indicating they represent two halves of the same L1 element. The configuration indicates a mechanism of retrotransposon-mediated rearrangement where second-strand synthesis occurs in trans between two distant L1 retrotransposition reactions, including one twin priming event, in a way analogous to that described above for reciprocal translocations. However, in this case, the two L1s involved are located on the same chromosomal arm rather than on distinct chromosomes (**Fig. 6c**). Another example of a megabase-scale paracentric inversion is described in **Supplementary Fig. 9a-c**. Although L1-mediated reciprocal inversions are not very frequent in our cohort as reciprocal translocations are, they represent a type of balanced structural variation that remained largely undetected in previous work using short reads only ^10^.

**Figure 6.**
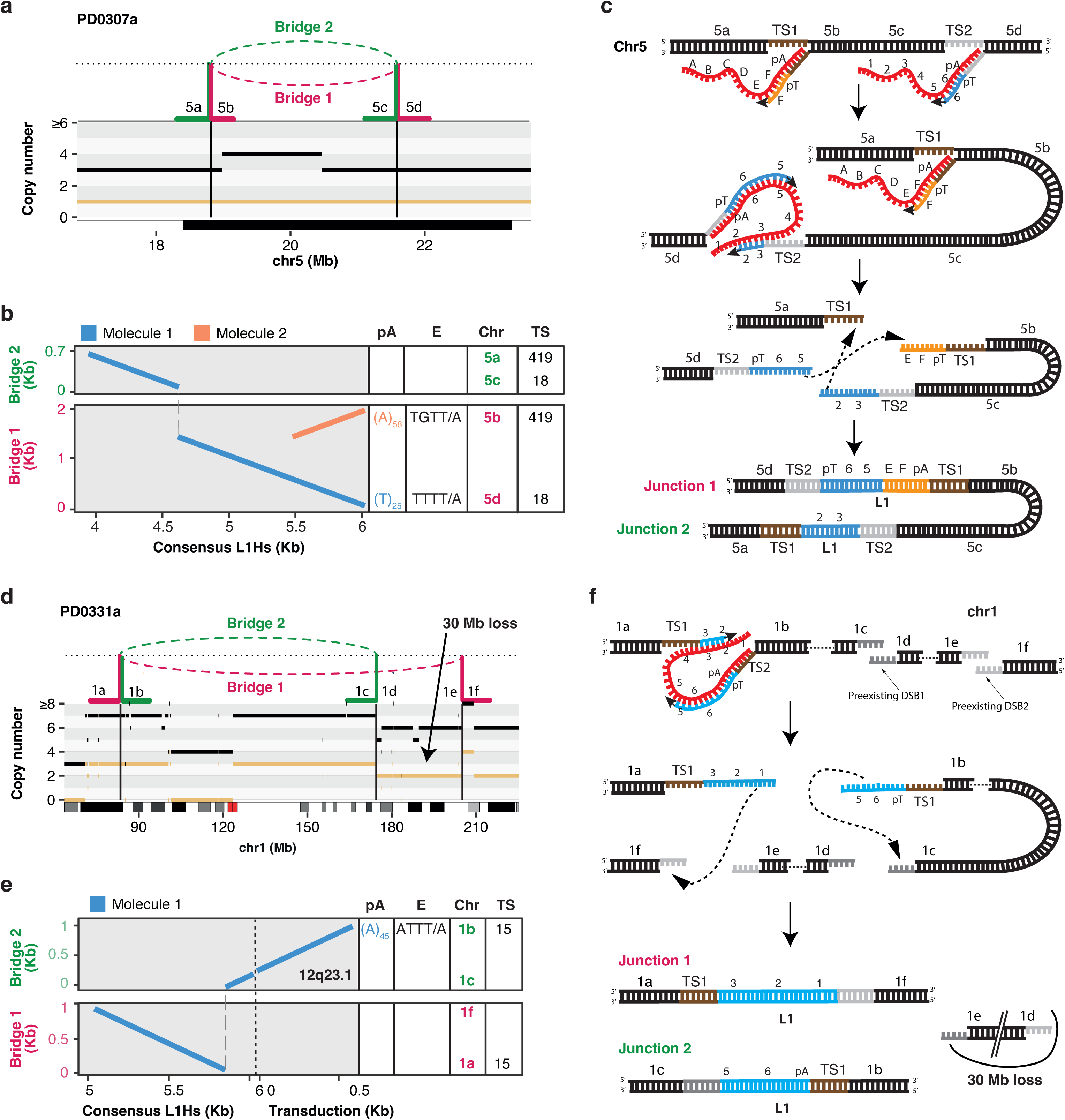
Somatic retrotransposition promotes reciprocal inversions and complex structural variation. **(a)** A paracentric inversion in chromosome 5 from tumour PD0307a, with two L1 bridges, where bridge 1 joins chr5b with chr5d and bridge 2 joins chr5a with chr5c. **(b)** Structure of the bridges for the inversion shown in (a). Here, Bridge 1 exhibits a poly(A) and TGTT/A endonuclease motif at its 3’ end, while bridge 2 shows a poly(T) tail and TTTT/A endonuclease motif at its 5’ end. Bridge 1 and bridge 2 are flanked by a heteroduplication of 18 bp and 419 bp. **(c)** Mechanism explaining the configuration of the bridges and genomic rearrangement described in (a-b), requiring two L1 insertions, one following canonical TPRT and the other following twin priming. A genomic inversion with two L1 bridges results after recombination. **(d)** A complex L1-mediated structural variant at chromosome 1 in tumour PD0331a involving two L1 bridges. Here, bridge 1 joins chr1a with chr1f, while bridge 2 joins chr1b with chr1c. As a result, they are generated a virtual acentric chromosome, by joining chr1a with chr1f, and a virtual circular chromosome by joining chr1b with chr1c, together with a 30 Mb deletion between chr1d and chr1e. **(e)** Dot plots reveal the structure of the L1 bridges depicted in (d), revealing the involvement of one single L1 DNA molecule (blue). Here, bridge 2 exhibits a poly(A) tail and an ATTT/A endonuclease motif at the 3’ end. Of note, the two L1 bridges share the same inversion breakpoint, suggesting twin priming. **(f)** Mechanism for the complex rearrangement at chromosome 1 in tumour PD0331a, explaining the configuration of the bridges and the genomic rearrangements observed in (d-e). The mechanism requires one insertion following twin priming, whose cDNA growing molecules are used to repair the breakpoints from two different preexisting doubled-strand breaks. These anomalous repairs generate a virtual acentric chromosome, a virtual circular chromosome by joining chr1b with chr1c, and a 30 Mb-long fragment of DNA lacking a centromere that is lost.

Finally, the same mechanism can promote more complex types of structural variation where one or more synchronous retrotransposition events can be involved in mediating two or more bridges from a series of concatenated rearrangements (**Fig. 5h**). The rearrangements involved here can be very heterogeneous and include inversion, translocations, deletions or duplications. For example, in tumour PD0331a, the two cDNA ends from a single L1 insertion event following twin priming on chromosome 1p, engage with two distant pre-existing DNA breaks for second-strand synthesis. This process results in two retrotransposon bridges that mediate a virtual acentric chromosome and a virtual circular chromosome, with a 30 Mb loss between the two original breaks (**Fig. 6d-f**). Similarly, we identified in the same tumour another complex event involving two independent retrotransposition events, one of which is a processed pseudogene from the *STK3* gene and the other a L1, that initiate insertion at chromosomes 12p and 18q, respectively (**Supplementary Fig. 9d-f**). The cDNA ends from these insertions engage to each one of the two ends from a pre-existing double-strand DNA break that occurred at chromosomal arm 12q. After second-strand DNA synthesis, this resulted in two retrotransposition bridges, one involving the *STK3* pseudogene and the other the L1 event, that repair the mentioned DNA break ends but concomitantly split chromosome 12, generating an inversion-like rearrangement on chromosome 12 and an unbalanced translocation between chromosomes 12q and 18q (**Supplementary Fig. 9d-f**).

Overall, our results illuminate distinct patterns and mechanisms by which aberrant integration of one or more somatic retrotransposition events can mediate different types of somatic structural variation in cancer. The mechanisms suggest that distinct somatic retrotranspositions act in a coordinate and synchronous manner. Most of these rearrangements remained unexplored in previous work due to the size of the retrotransposition bridges involved and to the absence of substantial copy number changes associated to them ^10^. Our findings complete the picture of a hidden landscape of retrotransposition-mediated structural variation in human cancer.

## The tempo of somatic retrotransposition in human cancer

Previous cancer retrotransposition analyses have shown that somatic retrotranspositions can occur at different stages of tumour progression, including pretumoral lesions, primary tumours and metastases ^7,10,22,24,47,48^. However, the extent to which retrotransposition contributes to mutational burden throughout the evolution of tumours remains poorly understood. Current computational approaches for timing inference rely on the estimation of the variant allele frequency (VAF) and the local copy number state from short reads to classify variants as early or late events ^49,50^. To deal with retrotransposons, we first developed MEIGA-SR (MEIGA short-reads), an algorithm specifically designed for VAF estimation of retrotransposition events from Illumina sequencing. Then, our approach employed the estimated VAF, along with tumour purity and copy number information, to obtain a relative timing estimation of somatic retrotransposition events (details of the algorithm and its validation in **Supplementary Note** and **Supplementary Fig. 10a-c**).

We firstly ran our timing approach on the 6,266 canonical somatic retrotransposition events from the cohort, which provided a relative timing estimation for 74% (4,644/6,266) of the insertions. The results revealed that 64.1% (2975/4644) and 14.9% (692/4644) of the events are clonal early and clonal late, respectively, while 14.5% (674/4644) are clonal unassigned (NA) events (**Fig. 7a**). Of note, this analysis is biased towards detecting clonal events, as mutations are more easily detectable when they display higher clonality. Consequently, the percentages of timing categories are not directly comparable within a sample. Despite this constraint, we still find that 6.5 % (303/4644) are subclonal (late) events in our cohort (**Fig. 7a**). Our analysis shows a considerable variability between tumours, with clonal early events being the most frequent (ranging from 27.4% to 77.9%) in all but one sample (PD0287a), followed by clonal late (0.7%-34.8%), clonal NA (0%-35.1%), and subclonal (0.8%-14.6%) (**Fig. 7a**). This analysis confirms that retrotransposition is a mutational process active in both early and late stages of tumour progression.

**Figure 7.**
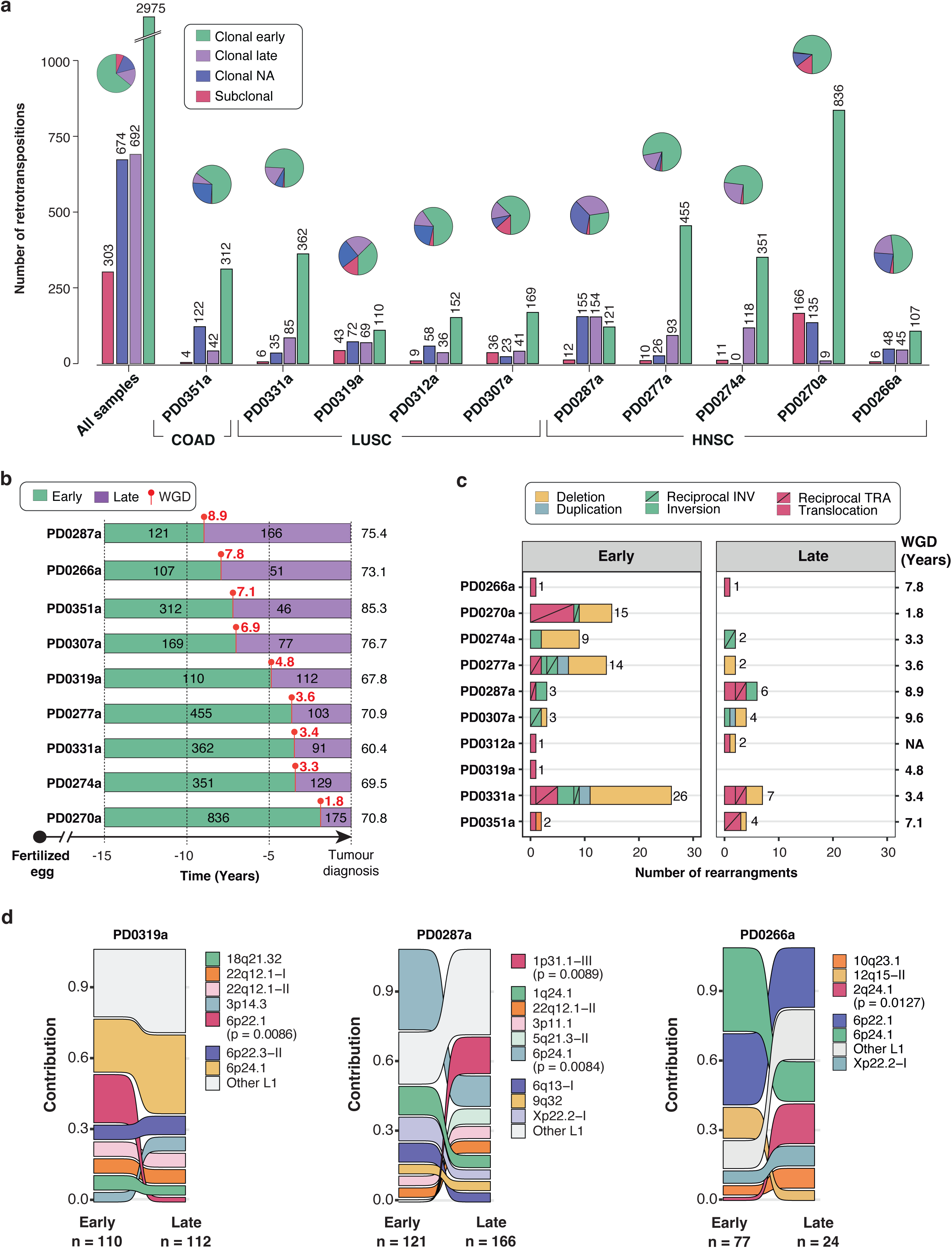
The tempo of somatic retrotransposition in human cancer. **(a)** Relative-time timing estimation of 4,644 canonical retrotransposition events in our cohort according to four timing categories: early clonal (green), late clonal (purple), clonal NA (blue), subclonal. **(b)** Real-time timing estimation of canonical retrotransposon insertions throughout patients’ lifetime in samples with whole-genome duplication (WGD) events. The X axis shows the time interval before (green) and after (purple) the WGD event. Red lollipops indicate the timing of WGD events with red numbers showing the estimated years before cancer diagnosis when the WGD occurred. Numbers within green and purple timelines represent the number of insertion events. Numbers at the end of the timeline represent patient age at biopsy. **(c)** Numbers and types of retrotransposon-mediated rearrangements per tumour, categorized by early (before WGD) and late (after WGD) timing. Numbers on the right side indicate years before cancer diagnosis when the WGD event occurred. **(d)** Relative contribution of each L1 source element to the retrotransposition burden during primary tumour progression in samples PD0319a, PD0287a, and PD0266a. Significant activity changes between early and late stages are shown with p-values. The total number of somatic insertion events in early and late stages is indicated below the timing labels.

To further characterize the tempo of somatic retrotransposition in tumorigenesis, we conducted a real-time timing estimation approach (**Supplementary Note**). The method relies on the analysis of mutational clock signatures that correlate with patient age at diagnosis, which can be used for timing of whole-genome doubling (WGD) events – often observed early in tumorigenesis ^51,52^ – and their associated variants ^49,50^. While all tumours in our study exhibited one WGD event, in tumour PD0312a we find more than one, confounding timing estimation. Consequently, this tumour was excluded from the analysis. In general, our approach determined that WGDs occurred at a median time of 4.77 years before diagnosis, ranging from 1.77 to 8.87 years, and placing 2,823 and 950 retrotranspositions events before and after a WGD event, respectively (**Fig. 7b**). Of note, all analysed tumours exhibit more than a hundred retrotransposition events before the whole-genome duplication. For instance, in tumour PD0287a, 121 out of 287 clonal events occurred more than 8 years before the patient was diagnosed with HNSC (**Fig. 7b**). Overall, these results indicate that activation of somatic retrotransposition is not merely the consequence of the genomic chaos typically governing in later stages of tumour progression, following the first WGD event. Instead, it appears to be a mutational process triggered early during tumour development, which can be very active.

Secondly, the analysis of 152 retrotransposon-mediated rearrangements provided a confident timing estimation for 103 retrotransposon bridges, of which three-quarters (n=75) were classified as early and one-quarter (n=28) as late events (**Fig. 7c**). For example, in PD0331a, we observe a total of 26 RT-RGs occurring 3.4 years before diagnosis (confident interval range = 2.5 – 5.8), including four reciprocal translocations and one reciprocal inversion. Notably, nine L1-mediated reciprocal translocations in the cohort occurred before the first WGD event. This includes one reciprocal translocation in PD0287a that took place more than 8.87 years before tumour diagnosis (confident interval range = 7.2 - 22.2; **Fig. 7c**). Overall, our work confirms that L1 elements are primary contributors of cancer genome architecture and identifies L1 activity as an early cause of the large-scale genome instability that typically characterizes tumorigenesis.

Finally, by utilizing the information from both transductions and derived solos, we investigated the activity of source elements throughout primary tumour evolution to unprecedented resolution ^7,10^. Overall, we observe a positive correlation between the number of derived insertions per sample and the number of active elements early in tumour evolution (Pearson correlation test, R=0.71, p=0.021; **Supplementary Fig. 10e**), a trend that persists in the later stages (R=0.8, p=0.0052). Of note, in general tumours have a substantial number of L1 loci already active at early stages (median = 23.5, ranging from 8 to 57; for late, median = 16, range = 3-30 **Supplementary Fig. 10e**). The most extreme example in our cohort is tumour PD0270a, with 56 active source elements, contributing to a total of 426 derived copies. Although in general the relative contribution of individual L1 source elements stays stable throughout primary tumour progression, we observe instances in which specific elements significantly changed their activity rates between timing categories. For example, in tumour PD0319a, the activity of a source element at 6p22.1 represented 20.41% (traced copies: 10/49) of the total retrotransposition burden in the early stage, ranking the second in the activity, but became nearly inactive in late stages of tumour progression, ranking the seventh (**Fig. 7d**). Conversely, in tumour PD0287a, we observe late activation of the source element at 1p31.1-III, ultimately becoming the most active source L1 element in the later stages of tumoral progression (15.8%; traced copies: 12/76) (**Fig. 7d**). Thus, our data confirm that L1 activity is a dynamic process in the primary tumours, which can be reversed through tumour progression ^7^, resulting in a variable relative contribution of L1 source elements to insertional mutagenesis along tumour progression.

## DISCUSSION

Here, we present an extensive set of somatic retrotransposition events, identified in human tumors with exceptionally high L1 activity using whole-genome long-read sequencing analysis. This strategy illuminates internal sequences of L1 source elements and their progeny, allowing unambiguous tracing of progenitor copies for many somatic L1 insertions. Our results confirm that only a limited number of L1 loci are enabled for retrotransposition in cancer genomes ^7,10,22,37,53^. We find that retrotransposition activity in a tumour is the result of the balance between two processes. On the one hand, the activity of a small set of full-length elements, poised for mobilization through promoter hypomethylation, which increase the number of somatic retrotransposition events. On the other hand, the operation of host and L1 replication processes that promote internal rearrangements during integration, rendering most L1 somatic copies inactive and thereby limiting the exponential escalation of retrotransposition. Our analysis unveiled the variety of mechanisms behind L1 integration, recapitulating findings from engineered L1 elements mobilized in transformed cultured cells ^2,16,17^. Additionally, we observed evidence for previously undescribed or poorly documented mechanisms, including twin-priming and switching, and second cDNA strand synthesis in trans. Together with L1-mediated deletions ^10^, these insertion-mediated processes significantly impact tumor genome architecture by driving balanced translocations, inversions, and even more complex rearrangements. Our data reveal a relatively frequent occurrence of retrotransposon-mediated rearrangements, estimated in one event in every 40 canonical insertions. We speculate that the spatiotemporal proximity of the involved loci might be facilitated by chromatin conformation potentially involving molecular condensates. Our findings depict a panorama where L1 retrotransposition acts as an early mutational process in tumour evolution, preceding the first whole-genome doubling event, which typically occurs in early cancerous lesions ^51,52^. Overall, these observations highlight L1 activity as a more significant player in tumor genome plasticity than previously anticipated, extending beyond simple insertional mutagenesis ^26^. To better understand the dynamics of somatic retrotransposition in cancer, we will need to study L1 regulation at the locus-level ^27,44,54–56^ and in a multidimensional scale, throughout different stages of malignant transformation– from normal tissues to metastases –, from the bulk tumour to the single-cell, and integrating genomics with transcriptomic and epigenomic data.

## SUPPLEMENTARY TABLES LEGENDS

**Supplementary Table 1.** Screening of somatic retrotranspositions in 137 cancer genomes

**Supplementary Table 2.** Annotation of 6,266 canonical somatic retrotransposition events identified in 10 tumours with high somatic retrotransposition rates

**Supplementary Table 3.** Annotation of 152 retrotransposition-mediated rearrangements identified in 10 tumours with high somatic retrotransposition rates

**Supplementary Table 4.** Annotation of source element activity inferred with the Source Inference method

**Supplementary Table 5.** ONT sequencing stats for 10 high retrotransposition rate tumours and adjacent tissues

**Supplementary Table 6.** Simulation of 6,420 retrotransposon insertions with MEIsimulator

**Supplementary Table 7.** Simulation of 70 retrotransposition-mediated rearrangements with MEIsimulator

**Supplementary Table 8.** Benchmarking stats for simulated canonical retrotransposition insertions including MEIGA and three additional callers

**Supplementary Table 9.** Recalling stats for simulated canonical retrotransposition insertions categorized by insertion type

**Supplementary Table 10.** Computational time and memory requirements for retrotransposition callers in detecting simulated insertions at different VAFs

**Supplementary Table 11.** Evaluation of MEIGA in sample PD0270a using additional sequencing technologies and pipelines

**Supplementary Table 12.** Genome-wide mean methylation estimated from ONT sequencing

**Supplementary Table 13.** Mean methylation per source element estimated from ONT sequencing

**Supplementary Table 14.** Timing of whole-genome doubling (WGD) events

**Supplementary Table 15.** Evaluation of retrotransposons timing approach in a multi-region setup

## METHODS

### Cancer datasets

#### Screening of high-retrotransposition rate tumours

We performed shallow sequencing with Illumina paired-ends, 350 bp insert size and 150 bp read length to obtain the whole genomes of 137 tumours, including head-and-neck squamous carcinoma (n = 37), lung squamous carcinoma (n = 50) and colorectal adenocarcinoma (n = 50). This approach resulted in an average coverage of 11.7x (95% confidence interval, ranging from 11.5x to 11.9x; **Supplementary Fig 1a-b**). Alignments to the reference human genome (assembly GRCh38) were performed using bwa mem v0.7.17 ^58^ and processed with samtools v1.12 ^59^. Duplicate reads were identified and removed from subsequent analyses using biobambam2 v2.0.87 ^60^. The resulting BAM files were then analysed using xTea ^61^ for detecting L1 retrotransposition events. To confidently identify somatic events, given the absence of genomic data from adjacent tissues at this stage, we genotyped the resulting L1 insertions using MEIGA-SR (i.e., MEIGA-Short Reads, described in section 5.1) and excluded those events shared by two or more donors within the tumoral cohort. Additionally, we excluded events documented in the database of retrotransposon polymorphisms generated within the framework of the 1000 Genomes Project ^8^. We then selected 10 tumours with a minimum of 100 somatic retrotranspositions for multiplatform whole-genome sequencing with short and long reads. See **Supplementary Table 1**.

#### Whole-genome sequencing of high-retrotransposition tumours with short and long reads

To achieve a comprehensive genomic characterization of the 10 tumours with high retrotranspositions rates selected in the screening stage, we performed whole-genome sequencing of these tumours and their matched-normal (non-tumoral adjacent tissues) counterparts with short (Illumina) and long (Oxford Nanopore Technologies, ONT) reads. For Illumina, we carried out paired-end libraries with a 350 bp insert size and 150 bp reads, each to a final coverage of ∼30x. Alignments to the reference human genome and marking of read duplicates were performed as described above. For single-molecule sequencing with ONT, we used the Short Read Eliminator XS buffer (Circulomics, Maryland, USA) to remove DNA fragments shorter than 5 kb. The DNA was then purified using Agencourt AMPure XP magnetic beads (Beckman Coulter, California, USA), following the manufacturer’s instructions. Libraries were constructed using the Oxford Nanopore Sequencing ligation library preparation kit (SQK-LSK109, Oxford Nanopore Technologies Ltd) according to the manufacturer’s protocol, including an initial DNA end-repair and dA-tailing step using the NEBNext End Repair/dA-tailing module (NEB). Then, libraries were loaded into MinION R9.4 flow cells (FLO-MIN106, Oxford Nanopore Technologies Ltd) for sequencing. MinION devices were controlled by the MinKNOW software (v21.02.1 to v22.10.10). High-accuracy base calling was performed using the GPU-dependent Guppy software v6.1.5 (Model: dna_r9.4.1_450bps_hac). The resulting fastq files were aligned to the reference genome (GRCh38) with minimap2 v2.24 ^6^, and the generated alignments underwent sorting and quality-based filtering with samtools v1.12. All the tumours were sequenced to a minimum coverage of 30x and resulted in a median N50 of 19.9 kb (Range = 13.9-24.7, **Supplementary Fig. 1c-d**; **Supplementary Table 5**). Additionally, the tumour with the highest rate of somatic retrotransposition (PD0270a) was further sequenced using an alternative long-read platform, namely PacBio HiFi, alongside its normal adjacent tissue. The sequencing coverage was 43x and 19x, with an N50 read length of 18.3 kb and 15.3 kb, for the tumour and normal adjacent tissue, respectively.

### Analysis of somatic retrotranspositions

#### Identification of somatic retrotranspositions with MEIGA

To find and characterize somatic retrotransposition in the long reads’ dataset, we developed a bioinformatics algorithm coined MEIGA (Mobile Element Integration Genome Analyzer). The method relies on the identification of read clusters indicative of structural variation breakpoints, followed by a reconstruction of such variants through local assembly and the identification of hallmarks of somatic retrotransposition. MEIGA can detect seven main types of somatic retrotranspositions according to the type of sequence retrotransposed. These types are solo retrotranspositions events (L1-solo, Alu-solo and SVA-solo), L1-partnered transductions, L1-orphan transductions, processed pseudogenes, and solitary polyadenylate [poly(A)] tracts. Notably, MEIGA not only detects canonical insertions but also retrotransposition-mediated rearrangements resulting from aberrant integration events. The algorithm is run in tumour-normal paired mode. See description and validation of the method in **Supplementary Note**; **Supplementary Fig. 2** and **Supplementary Fig. 3**.

#### MEIGA benchmarking

To establish a robust benchmarking framework for MEIGA, we developed MEIsimulator, a bioinformatics tool designed to generate sequencing data (short reads and long reads) from synthetic genomes that incorporate a wide range of randomly integrated retrotransposition events. MEIsimulator can also simulate aberrant retrotranspositions that result in SVs such as deletions, duplications, inversions, and translocations. Then, to evaluate MEIGA, we used MEIsimulator with default settings to simulate a total of 6,420 retrotransposon insertions (**Supplementary Table 6**). Additionally, we conducted simulations of L1-mediated genomic rearrangements, which included 20 deletions, 20 duplications, 20 inversions, and 10 translocations (**Supplementary Table 7**). These sets of retrotransposition events were replicated in both short-read and long-read sequencing datasets. In the short-read simulation, we set a sequencing depth of 30x and a VAF of 50%. For the long-read simulation, retrotransposition events were simulated at different VAF levels, including 0.15, 0.3 and 0,5, and a sequencing depth of 33.7x, which corresponds to the median depth of our ONT cohort. We then run MEIGA together with three additional reference pipelines to detect retrotranspositions in these mock genomes, including rMETL v1.0.4 ^63^, xTea v0.1.9 ^61^, and PALMER v2.0.0 ^64^ (**Fig. 1b**; **Supplementary Fig. 2c-d**; **Supplementary Table 8 and 9**). In addition, we used Blast v2.11.0 ^65^ to evaluate the performance of MEIGA to reconstruct the nucleotide sequences of the simulated insertions from above, by analysing the size and the nucleotide identity between observed and expected sequences (**Supplementary Fig. 2e**).

#### Validation of MEIGA retrotransposition calls

We ran MEIGA on the long-read sequencing data from the 10 relevant tumours and their adjacent tissues. To validate our results, we carried out further sequencing in the tumour with the highest number of canonical somatic retrotranspositions (PD0270a), including two additional technologies (Illumina paired-ends and PacBio HiFi). Then, we analysed somatic retrotransposition with other relevant pipelines, including xTea-SR for the Illumina short reads and xTea-LR for the ONT and PacBio long reads. The intersection of the insertions called by each method is shown in **Supplementary Fig. 3a** and **Supplementary Table 11**. Our results find that 92.5% (1,193/1,290) of MEIGA calls for ONT, are validated by at least one external method (xTea-SR or xTea-LR) or an additional technology (Illumina paired-ends or PacBio HiFi). To validate the fraction of MEIGA private calls (i.e., those events that do not intersect with any other pipeline nor technology; accounting for 97 calls on the ONT dataset, 109 calls on the PB dataset, and 112 calls on both ONT and PB), we carried out visual inspection with Integrative Genomics Viewer (IGV) of a random selection of 150 insertions (50 MEIGA private calls from the ONT dataset, 50 private calls from the PB dataset, and 50 private calls from the ONT+PB dataset). This analysis confirmed a true positive rate of 94.7%. (142/150) (**Supplementary Fig. 3b**). More benchmarking details in **Supplementary Note** and **Supplementary Fig. 3c-d**.

#### Annotation of the internal structure of L1 insertions

Somatic L1 insertion calls are subject to an additional annotation procedure to analyse their structural configuration. First, the sequence corresponding to the target site duplication was detected and trimmed from either the 5’ or 3’ end of the L1 insert and insertions in the minus strand were reverse complemented. Then, poly(A) tails were detected, with L1 insertions containing a single poly(A) tract being classified as solo events, while those containing two tracts were annotated as 3’ transduction candidates. To trace transductions to their full-length L1 progenitor, transduced sequences were aligned with BWA-MEM 0.7.17-r1188 to a patient specific database containing the sequences corresponding to the 10 kb genomic intervals downstream to each of the full-length L1s (germline plus somatic) detected for a given patient. The inserted sequences were then further trimmed by removing the poly(A) tails and transduced sequences, resulting in trimmed sequences corresponding to L1 sequences alone. Trimmed inserts were aligned using BWA-MEM into a consensus L1 sequence and classified as full-length, 5’ truncated, 5’ inverted or complex L1 insertion events, based on the alignment hits. As a last step, the inversion junction conformation for every twin priming event was determined based on the alignment position over the consensus for the inverted and non-inverted L1 pieces. More details in **Supplementary Note**.

#### Haplotype phasing of long reads

We carried out the phasing of germline polymorphisms and reconstructed the parental haplotypes from long reads, which will be later employed in the characterization of L1 source candidates (see next section). First, we run Freebayes v1.3.6 [arXiv:1207.3907] on the Illumina short-read sequencing data obtained from matched-normal samples to identify germline polymorphisms (SNPs and short indels). Then, we run Whatshap v1.2.1 [doi:10.1101/085050] on the ONT long-read sequencing data to first construct a graph that captures the different ways the germline variants could be combined, and then performed phasing inference by assigning the most likely haplotype to each parental chromosome. In a final step, we employed WhatsHap to add phasing tags to each alignment that could be confidently associated with either parental haplotype. This step was performed on the alignments included in the ONT long-read sequencing data derived from both the tumour and matched normal tissues. Of note, the phasing strategy obtained haplotype blocks longer than 1.5 Mb for both the adjacent tissues and the tumours (**Supplementary Fig. 1e-f**).

#### Identification of L1 source candidates

We aimed to identify all potentially active L1 loci in the cohort achieving haplotype-level resolution, from a dataset of 693 L1 candidates that have a complete length and structure initially described in PCAWG ^10^ and HGSVC2 ^8^. To accomplish this, we used the ONT phased BAM files obtained from matched-normal samples. We employed two different approaches depending on whether the source elements were present or absent on the reference genome. First, for the L1 elements present in the reference genome, reconstruction of the entire sequence was performed using the ‘refSrc_seqs.py’ algorithm included in the MEIGA suite. Second, for the non-reference source L1 elements, we adopted a distinct approach. As these elements manifest as insertions in the genome, we employed MEIGA, using default parameters, to identify full-length solo L1 insertions in the matched-normal tissues. In cases of homozygous non-reference L1 loci, we re-ran MEIGA on the target regions employing the ‘--hpTag’ parameter for the reconstruction of haplotype-specific sequences. For heterozygous non-reference L1s, we relied on the insertion sequences initially reconstructed by MEIGA.

#### The Source Inference method

Our source inference method elucidates the element-of-origin of somatic L1s through the identification of specific SNVs shared between these L1 sources and their derived insertions. The method starts by genotyping all germinal full-length L1 loci within each patient’s normal genome to identify the diagnostic SNVs that characterize each L1 source. Then, the algorithm identifies SNVs in the repertoire of L1 somatic copies in the patient’s tumour genome. Finally, the element-of-origin of each individual somatic copy is inferred by comparing the set of SNVs present in each somatic copy with the set of diagnostic SNVs that characterize each source L1 A description of the workflow is available in **Supplementary Note**.

To evaluate our method, we simulated the mobilization of all potential source elements (n=266) from the patient in the cohort with the highest L1 activity rate (PD0270), assuming a homogeneous activity of 20 retrotranspositions for each one of the 266 elements and with the derived copies showing the same insertion size distribution as in 2954 cancer genomes from the PCAWG project ^10^. The results showed a high specificity of the pipeline (>99%) and indicated an overall sensitivity of 47.6% (2530/5320), with a drop in sensitivity when somatic L1s are derived from younger source elements in relation with a decrease of genetic divergence (sensitivity = 59.2% [1042/1660] for pre-Ta; 47.0% [583/1240] for Ta-0; 36.8% [735/2000] for Ta-1). We also observe that the performance of the method is notably affected by the size of the derived copies, with a reduction in the sensitivity inferred from smaller derived copies, particularly in those copies derived from Ta-1 source elements (**Supplementary Fig. 5**). Further evaluation in **Supplementary Note**.

#### Identification of hot L1 elements and L1 phylogenetic analysis

Hot L1 elements must meet a minimum of one of the following conditions: (i) L1 activity rate, defined as the average number of mobilizations mediated by a source element when found activated) is equal or more than 10; (ii) the relative contribution to the total amount of traced mobilizations in the cohort is equal or more than 1.5%. This analysis finds 27 hot L1 germline in our cohort (**Supplementary Fig. 6c**). These hot sources were aligned with muscle v3.36.0 ^66^. And a L1 phylogenetic tree was built using the following R packages: Ape v5.6-2 ^67^ and Phangorn v2.8.1 ^68^. Nucleotide substitution model selection was performed with phangorn::modelTest ^69^. GRT + G + I model was selected based on AIC (Akaike Information Criterion) score. A maximum likelihood tree was estimated with the phangorn::optim.pml function, starting from a distance-based tree. Tree topology was adjusted via nearest neighbour interchange ^70^. After maximum likelihood optimization, midpoint rooting was applied, rendering L1Pt as the expected tree outgroup. Non-parametric bootstrap analysis with 1000 replicates was done to determine the node consistency along the inferred tree. For display purposes, only values informative of high support (≥80%) were labelled to the tree nodes. L1Hs subfamily assignment was performed through the identification of subfamily diagnostic nucleotide positions on their 3’ end of the L1 sequence ^71^ (see **Supplementary Note**).

#### Analysis of methylation profiles

Estimates of methylation content for all CpG sites were calculated with Megalodon26 (v2.5.0) using the Remora neuronal network model dna_r9.4.1_e8. The configuration for the remora-modified-bases was set with the flags: ‘dna_r9.4.1_e8 hac 0.0.0 5hmc_5mc CG 0’. Epigenetic profiles for individual L1 loci were established by averaging the DNA methylation levels across CpG dinucleotides within the bodies, promoters, and adjacent genomic regions, spanning 20 kb upstream and downstream, using BEDtools v2.31.027. Disparities between tumour samples and their corresponding adjacent healthy tissues were quantified using Gtools v3.9.428, and statistical significance was determined employing two-sided Mann-Whitney-Wilcoxon tests. See **Supplementary Fig. 7** and **Supplementary Tables 12 and 13**.

#### Detection of polyadenylation signals downstream L1 loci

APARENT ^43^ is a deep neural network designed to detect and score all polyadenylation sites within a given DNA sequence. In this study, we employed this software to identify both canonical and alternative polyadenylation signals and to estimate their relative strengths. For each tumour sample, APARENT was applied to the reconstructed sequences of source L1 elements along with their unique downstream sequences. Subsequently, we matched this information with the rates of solo insertions and transductions.

#### Identification of retrotransposon-mediated rearrangements

Rearrangements mediated by retrotransposition manifest as junctions between two distant breakpoints in the genome with retrotransposon bridges connecting them. MEIGA identified a total of 152 retrotransposition-mediated junctions in the 10 tumours cohort. These junctions were classified into four primary categories, determined by (i) whether they were interchromosomal or intrachromosomal events and (ii) the orientation of their breakpoints (note that we have not considered associated copy-number changes in this classification). For intrachromosomal junctions, our naming convention was as follows: deletion-like for [+/-], duplication-like for [-/+] and inversion-like, which encompasses both head-to-head [+/+] and tail-to-tail [-/-] inversions. Interchromosomal junctions were categorized as translocation-like. See **Supplementary Note**.

#### Validation of L1-mediated reciprocal translocations by cytogenetics

To validate our findings, we included an additional lung adenocarcinoma cell line, NCI-H2009, and its corresponding normal counterpart, NCI-BL2009, in our study. We conducted FISH experiments on the NCI-H2009 cell line to assess the occurrence of a reciprocal translocation between chromosomes 3 and 6 mediated by the somatic integration of a L1. We employed commercially available whole chromosome painting (WCP) probes targeting chromosomes 3 and 6 labelled with either green or red fluorescent dyes (FWCP-03 and FWCP-06, respectively; Creative Bioarray). Metaphase spreads were obtained from NCI-H2009 cells following standard techniques, pre-treated with RNAse and pepsin and subjected to dual FISH experiments as per the provider’s recommendations. After a counterstaining with DAPI (0.14 pg/ml), photographs were taken for each individual colour with a Nikon Eclipse-800 fluorescence microscope (Tokyo, Japan) equipped with a DS-Qi1Mc CCD camera (Nikon) and controlled the using NIS-Elements software (Nikon). The resulting images were merged and processed using Adobe Photoshop CS6 (San Jose, CA, USA). See **Supplementary Fig. 8c-e**.

#### Validation of L1-mediated reciprocal translocations by chromatin conformation capture

We carried out Micro-C to detect 3D chromatin contacts at genome-wide level in the high-retrotransposition rate cancer cell line NCI-H2009 ^7^. This is a 3C-based technique that employs an MNase to digest the genome at mononucleosome level to generate genome-wide contact maps with high resolution. We prepared 4 Micro-C libraries for NCI-H2009 following Dovetail Micro-C Kit protocol (Dovetail Genomics). Final libraries were sequenced at 150bp PE by an external service (Macrogen Inc.) using the HiSeqX sequencer from Illumina. Then, we aligned our Micro-C library onto the GRCh38 genome assembly using bwa-mem. Following the recommended Micro-C procedure, we used the parse module of the pairtools pipeline to find ligation junctions and other proximity ligations in our libraries. After sorting the parsed pairs, a deduplication step is performed, where pairtools dedup is used to detect molecules that could be formed via PCR duplication. These pairs are marked and excluded from downstream analysis. Finally, a 5 kb resolution contact map is generated using cooler, a widely used tool for managing storage and manipulation of matrices in the form of contact maps. See **Supplementary Fig. 8f**.

### Timing approaches

#### Real Time WGD Timing

The whole genome duplications (WGDs) were timed using an approach similar to that outlined in ^49^. The proportion of clonal clocklike CpG > TpG SNVs on different numbers of allelic copies, known as their multiplicities, was used to infer the WGD timing. We only considered genomic regions where the highest number of copies of either parental allele, known as the major copy number, was two. This is because all gains in these regions are assumed to occur through the WGD. We assumed that a linear acceleration of 5x occurred in the CpG > TpG mutation rate at a point uniformly distributed 1 and 15 years before diagnosis. Given a sampled acceleration timepoint, we found the WGD timing that corresponded to the maximum likelihood of the proportion of multiplicity one and two clocklike SNVs in genomic region. We marginalized the likelihood over the fraction of multiplicity one mutations that were subclonal and corrected the multiplicity proportions to account for our lack of power to detect SNVs with less than three reads. The uncertainty in our WGD timing estimates was estimated by bootstrapping over the SNVs in the tumour and resampling the acceleration timepoint. See **Supplementary Table 14**.

#### Timing of canonical retrotranspositions

Retrotransposition events were timed using MutationTimeR v 1.00.2 ^49^, using Illumina read counts, copy number profiles derived using Battenberg ^72^ and our pseudo-subclone structures. As with the real time WGD timing analysis using SNVs, we assumed the presence of a pseudo subclone corresponding to 30% of all biopsied tumour cells and accounting for 10% of all detected retrotranspositions. We then used phasing information from Nanopore sequencing to revise maximum likelihood timing classifications produced from MutationTimeR. Clonal retrotransposition events on one allelic copy in copy number regions with major copy number greater than one and minor copy number equal to one, were timed as clonal NA in MutationTimeR, as it is unclear whether they occurred after the gain on the major allele or at any time in the clonal period on the minor allele. We used a binomial model to measure the probability that the allele that the retrotransposition was phased to, was the major allele. If the maximum likelihood was that the retrotransposition occurred on the major allele, its timing was reclassified as clonal late. Only retrotransposition events exclusively phased to one allele and with less than 10% unphased reads were revaluated using this method. See **Supplementary Note** and **Supplementary Table 15**.

#### Timing of retrotransposon-mediated rearrangements

VAF estimation of retrotransposon insertions associated with rearrangements is more challenging than for canonical insertions. Here, the two breakpoints of a given rearrangement can vary in copy number, which may affect the analysis. Thus, we only considered those retrotransposon-mediated rearrangements whose breakpoints exhibited consistent timing categories. To assess the performance of our approach, we conducted a simulation involving 20 deletions, 20 duplications, 20 inversions, and 10 translocations mediated by L1, all with a VAF of 0.5. While we observed a larger dispersion of the VAF estimations compared to canonical insertions, our method consistently estimated VAFs centred at 0.5, with no significant differences observed among the various types of rearrangements studied (**Supplementary Fig. 10b**). Of note, the dispersion of estimations appeared to be notably higher for duplications, a difference that may be attributable to the fact that duplications double the number of reference reads, introducing a larger variability in the inferences.

## DATA AVAILABILITY

Fastq files corresponding to 137 tumours and their adjacent tissues will be available at the European Genome-Phenome Archive (EGA) before publication. This includes Illumina paired-ends, ONT, PacBio sequences. Links to these files will be provided before review (dataset are currently been uploaded to the database).

## CODE AVAILABILITY

The primary algorithm used for identifying somatic retrotransposition events in long-read sequencing data, MEIGA, is publicly accessible at:

https://gitlab.com/mobilegenomesgroup/MEIGA_LR (version 2.1.0).

Additional complementary methods utilized in this study are also publicly available at the following locations:

MEIsimulator (v1.0): https://gitlab.com/mobilegenomesgroup/MEIsimulator

MEIGA-SR (v1.1.8): https://gitlab.com/mobilegenomesgroup/MEIGA-SR

RT-Structure annotation (v1.0): https://gitlab.com/mobilegenomesgroup/rt-structure-annotation Source Inference (v1.0): https://gitlab.com/mobilegenomesgroup/SourceInference

### ACKNOWLEDGEMENTS

J.M.C.T. was supported by European Research Council (ERC) Starting Grant 716290 ‘SCUBA CANCERS’, Spanish Association for Cancer Research (AECC) grant LABAE20053TUBI, and Spanish Government grant PGC2018-102245-B-100. Currently, he is supported by Fundacion La Caixa grant HR22-00529, under the CaixaResearch Health Programme, and the Spanish Government grant PID2021-126493OB-100. B.R.-M. was supported by a PhD fellowship from Xunta de Galicia (ED481A-2016/151) and by a Bridging Excellence Fellowship provided by the Life Science Alliance. Currently, he is supported by the Spanish Ministry of Science and Innovation through the Centro de Excelencia Severo Ochoa (grants CEX2020-001049-S, MCIN/AEI /10.13039/501100011033), and the Generalitat de Catalunya through the CERCA programme. S.Z. was supported by Xunta de Galicia fellowship number ED481A-2018/199. M.S. was supported by Xunta de Galicia fellowship number ED481A-2017/306. N.P.E is supported by the Spanish Government grant PRE2022-103126. D.G-S. is supported by Xunta de Galicia grant number ED481D-2022-001. E.G.A. was supported by Spanish Government grant FPU17/05396. A.O. was supported by Xunta de Galicia fellowship number ED481A-2020/214. P.O. was supported by the Spanish Government grant FPU18/03421. I.O. is supported by Spanish Government fellowship FPU21/02665. P.V.L. and T.M.B. were supported by the Francis Crick Institute which receives its core funding from Cancer Research UK (CC2008), the UK Medical Research Council (CC2008), and the Wellcome Trust (CC2008). T.M.B. was supported by a PhD fellowship from Boehringer Ingelheim Fonds. P.V.L. is a CPRIT Scholar in Cancer Research and acknowledges CPRIT grant support (RR210006). M.M.-F. was supported by the Spanish Association Against Cancer (AECC) Scientific Foundation (grant INVES207MART) and is currently supported by the Miguel Servet program (grant CP20/00188) from the Instituto de Salud Carlos III (ISCIII) and the European Social Fund. GC is supported by Agence Nationale de la Recherche (ANR-15-IDEX-0001, ANR-11-LABX-0028) and CNRS (GDR 3546). We thank Dr David Posada and Dr Joao Alves for discussion on clonality analyses. We thank the Basque Biobank for sampling support. We thank the Galician Supercomputing Centre (CESGA) for computational resources. We thank Dr Ana Igea for management support. We thank the patients and their families for their participation in the project.

## AUTHOR CONTRIBUTIONS

J.M.C.T., B.R.-M., and S.Z. designed the study. J.M.C.T. and S.Z. wrote the manuscript with assistance from G.C., P.V.L., M.S. and B.R.-M. S.Z., B.R.-M., M.S. and E.G.A. designed somatic retrotransposition algorithms. S.Z. and N.P.E. performed computational analyses with support from D.G.S., J.T., I.O., P.O. Samples management, quality control and library preparation by A.P.-V., J.R.-C., A.O., M.M.-F., D.G.-S., I.D.-A. J.T. and D.G.-S. performed methylation analyses. D.G.-S. performed cytogenetics analysis. A.O. and I.O. carried out chromatin conformation analysis. L1 internal rearrangements analysis by S.Z., B.R.-M., J.T. and N.P.E. L1-rearrangements analysis by S.Z. Source elements inference by M.S. with support from S.Z. and B.R.-M. Timing approaches by S.Z., T.M.B. and P.V.L. Pipelines benchmarking by M.S., N.P.E. and S.Z..

## REFERENCES

1. Lander, E.S. et al. Initial sequencing and analysis of the human genome. Nature 409, 860–921 (2001).

2. Gilbert, N., Lutz, S., Morrish, T.A. & Moran, J.V. Multiple fates of L1 retrotransposition intermediates in cultured human cells. Mol Cell Biol 25, 7780–95 (2005).

3. Ostertag, E.M. & Kazazian, H.H., Jr. Twin priming: a proposed mechanism for the creation of inversions in L1 retrotransposition. Genome Res 11, 2059–65 (2001).

4. Brouha, B. et al. Hot L1s account for the bulk of retrotransposition in the human population. Proc Natl Acad Sci U S A 100, 5280–5 (2003).

5. Beck, C.R. et al. LINE-1 retrotransposition activity in human genomes. Cell 141, 1159–70 (2010).

6. Sassaman, D.M. et al. Many human L1 elements are capable of retrotransposition. Nat Genet 16, 37–43 (1997).

7. Tubio, J.M.C. et al. Mobile DNA in cancer. Extensive transduction of nonrepetitive DNA mediated by L1 retrotransposition in cancer genomes. Science 345, 1251343 (2014).

8. Ebert, P. et al. Haplotype-resolved diverse human genomes and integrated analysis of structural variation. Science 372(2021).

9. Beck, C.R., Garcia-Perez, J.L., Badge, R.M. & Moran, J.V. LINE-1 elements in structural variation and disease. Annu Rev Genomics Hum Genet 12, 187–215 (2011).

10. Rodriguez-Martin, B. et al. Pan-cancer analysis of whole genomes identifies driver rearrangements promoted by LINE-1 retrotransposition. Nat Genet 52, 306–319 (2020).

11. Kazazian, H.H., Jr. & Moran, J.V. Mobile DNA in Health and Disease. N Engl J Med 377, 361–370 (2017).

12. Kazazian, H.H., Jr., et al. Haemophilia A resulting from de novo insertion of L1 sequences represents a novel mechanism for mutation in man. Nature 332, 164–6 (1988).

13. Burns, K.H. Transposable elements in cancer. Nat Rev Cancer 17, 415–424 (2017).

14. Shukla, R. et al. Endogenous retrotransposition activates oncogenic pathways in hepatocellular carcinoma. Cell 153, 101–11 (2013).

15. Cajuso, T. et al. Retrotransposon insertions can initiate colorectal cancer and are associated with poor survival. Nat Commun 10, 4022 (2019).

16. Gilbert, N., Lutz-Prigge, S. & Moran, J.V. Genomic deletions created upon LINE-1 retrotransposition. Cell 110, 315–25 (2002).

17. Symer, D.E. et al. Human l1 retrotransposition is associated with genetic instability in vivo. Cell 110, 327–38 (2002).

18. Meyer, T.J., Srikanta, D., Conlin, E.M. & Batzer, M.A. Heads or tails: L1 insertion-associated 5’ homopolymeric sequences. Mob DNA 1, 7 (2010).

19. Myers, J.S. et al. A comprehensive analysis of recently integrated human Ta L1 elements. Am J Hum Genet 71, 312–26 (2002).

20. Gardner, E.J. et al. The Mobile Element Locator Tool (MELT): population-scale mobile element discovery and biology. Genome Res 27, 1916–1929 (2017).

21. Porubsky, D. et al. Recurrent inversion polymorphisms in humans associate with genetic instability and genomic disorders. Cell 185, 1986–2005 e26 (2022).

22. Scott, E.C. et al. A hot L1 retrotransposon evades somatic repression and initiates human colorectal cancer. Genome Res 26, 745–55 (2016).

23. Solyom, S. et al. Extensive somatic L1 retrotransposition in colorectal tumors. Genome Res 22, 2328–38 (2012).

24. Rodic, N. et al. Retrotransposon insertions in the clonal evolution of pancreatic ductal adenocarcinoma. Nat Med 21, 1060–4 (2015).

25. Pradhan, B. et al. Detection of subclonal L1 transductions in colorectal cancer by long-distance inverse-PCR and Nanopore sequencing. Sci Rep 7, 14521 (2017).

26. Lee, E. et al. Landscape of somatic retrotransposition in human cancers. Science 337, 967–71 (2012).

27. Ewing, A.D. et al. Nanopore Sequencing Enables Comprehensive Transposable Element Epigenomic Profiling. Mol Cell 80, 915–928 e5 (2020).

28. Shiraishi, Y. et al. Precise characterization of somatic complex structural variations from tumor/control paired long-read sequencing data with nanomonsv. Nucleic Acids Res 51, e74 (2023).

29. Luan, D.D., Korman, M.H., Jakubczak, J.L. & Eickbush, T.H. Reverse transcription of R2Bm RNA is primed by a nick at the chromosomal target site: a mechanism for non-LTR retrotransposition. Cell 72, 595–605 (1993).

30. Moran, J.V. et al. High frequency retrotransposition in cultured mammalian cells. Cell 87, 917–27 (1996).

31. Cost, G.J., Feng, Q., Jacquier, A. & Boeke, J.D. Human L1 element target-primed reverse transcription in vitro. EMBO J 21, 5899–910 (2002).

32. Monot, C. et al. The specificity and flexibility of l1 reverse transcription priming at imperfect T-tracts. PLoS Genet 9, e1003499 (2013).

33. Thawani, A., Ariza, A.J.F., Nogales, E. & Collins, K. Template and target-site recognition by human LINE-1 in retrotransposition. Nature 626, 186–193 (2024).

34. Zingler, N. et al. Analysis of 5’ junctions of human LINE-1 and Alu retrotransposons suggests an alternative model for 5’-end attachment requiring microhomology-mediated end-joining. Genome Res 15, 780–9 (2005).

35. Baldwin, E.T. et al. Structures, functions and adaptations of the human LINE-1 ORF2 protein. Nature 626, 194–206 (2024).

36. Rangwala, S.H., Zhang, L. & Kazazian, H.H., Jr. Many LINE1 elements contribute to the transcriptome of human somatic cells. Genome Biol 10, R100 (2009).

37. Philippe, C. et al. Activation of individual L1 retrotransposon instances is restricted to cell-type dependent permissive loci. Elife 5(2016).

38. Lanciano, S. & Cristofari, G. Measuring and interpreting transposable element expression. Nat Rev Genet 21, 721–736 (2020).

39. Moran, J.V., DeBerardinis, R.J. & Kazazian, H.H., Jr. Exon shuffling by L1 retrotransposition. Science 283, 1530–4 (1999).

40. Goodier, J.L., Ostertag, E.M. & Kazazian, H.H., Jr. Transduction of 3’-flanking sequences is common in L1 retrotransposition. Hum Mol Genet 9, 653–7 (2000).

41. Pickeral, O.K., Makalowski, W., Boguski, M.S. & Boeke, J.D. Frequent human genomic DNA transduction driven by LINE-1 retrotransposition. Genome Res 10, 411–5 (2000).

42. Consortium, I.T.P.-C.A.o.W.G. Pan-cancer analysis of whole genomes. Nature 578, 82–93 (2020).

43. Bogard, N., Linder, J., Rosenberg, A.B. & Seelig, G. A Deep Neural Network for Predicting and Engineering Alternative Polyadenylation. Cell 178, 91–106 e23 (2019).

44. Lanciano, S. et al. Locus-level L1 DNA methylation profiling reveals the epigenetic and transcriptional interplay between L1s and their integration sites. Cell Genom 4, 100498 (2024).

45. Sil, S. et al. Condensation of LINE-1 is critical for retrotransposition. Elife 12(2023).

46. Li, Y. et al. Patterns of somatic structural variation in human cancer genomes. Nature 578, 112–121 (2020).

47. Doucet-O’Hare, T.T. et al. LINE-1 expression and retrotransposition in Barrett’s esophagus and esophageal carcinoma. Proc Natl Acad Sci U S A 112, E4894–900 (2015).

48. Nguyen, T.H.M. et al. L1 Retrotransposon Heterogeneity in Ovarian Tumor Cell Evolution. Cell Rep 23, 3730–3740 (2018).

49. Gerstung, M. et al. The evolutionary history of 2,658 cancers. Nature 578, 122–128 (2020).

50. Jolly, C. & Van Loo, P. Timing somatic events in the evolution of cancer. Genome Biol 19, 95 (2018).

51. Gemble, S. et al. Genetic instability from a single S phase after whole-genome duplication. Nature 604, 146–151 (2022).

52. Davoli, T. & de Lange, T. The causes and consequences of polyploidy in normal development and cancer. Annu Rev Cell Dev Biol 27, 585–610 (2011).

53. Stow, E.C. et al. Organ-, sex- and age-dependent patterns of endogenous L1 mRNA expression at a single locus resolution. Nucleic Acids Res 49, 5813–5831 (2021).

54. Gerdes, P. et al. Locus-resolution analysis of L1 regulation and retrotransposition potential in mouse embryonic development. Genome Res 33, 1465–1481 (2023).

55. Freeman, B. et al. Analysis of epigenetic features characteristic of L1 loci expressed in human cells. Nucleic Acids Res 50, 1888–1907 (2022).

56. Taylor, D. et al. Locus-specific chromatin profiling of evolutionarily young transposable elements. Nucleic Acids Res 50, e33 (2022).

57. Espejo Valle-Inclan, J. & Cortes-Ciriano, I. ReConPlot: an R package for the visualization and interpretation of genomic rearrangements. Bioinformatics 39(2023).

58. Li, H. & Durbin, R. Fast and accurate short read alignment with Burrows-Wheeler transform. Bioinformatics 25, 1754–60 (2009).

59. Li, H. et al. The Sequence Alignment/Map format and SAMtools. Bioinformatics 25, 2078–9 (2009).

60. Tischler, G. & Leonard, S. biobambam: tools for read pair collation based algorithms on BAM files. Source Code for Biology and Medicine 9, 13 (2014).

61. Chu, C. et al. Comprehensive identification of transposable element insertions using multiple sequencing technologies. Nat Commun 12, 3836 (2021).

62. Li, H. Minimap and miniasm: fast mapping and de novo assembly for noisy long sequences. Bioinformatics 32, 2103–10 (2016).

63. Jiang, T., Liu, B., Li, J. & Wang, Y. rMETL: sensitive mobile element insertion detection with long read realignment. Bioinformatics 35, 3484–3486 (2019).

64. Zhou, W. et al. Identification and characterization of occult human-specific LINE-1 insertions using long-read sequencing technology. Nucleic Acids Res 48, 1146–1163 (2020).

65. Camacho, C. et al. BLAST+: architecture and applications. BMC Bioinformatics 10, 421 (2009).

66. Edgar, R.C. MUSCLE: multiple sequence alignment with high accuracy and high throughput. Nucleic Acids Res 32, 1792–7 (2004).

67. Paradis, E. & Schliep, K. ape 5.0: an environment for modern phylogenetics and evolutionary analyses in R. Bioinformatics 35, 526–528 (2019).

68. Schliep, K.P. phangorn: phylogenetic analysis in R. Bioinformatics 27, 592–3 (2011).

69. Posada, D. & Crandall, K.A. MODELTEST: testing the model of DNA substitution. Bioinformatics 14, 817–8 (1998).

70. Nguyen, L.T., Schmidt, H.A., von Haeseler, A. & Minh, B.Q. IQ-TREE: a fast and effective stochastic algorithm for estimating maximum-likelihood phylogenies. Mol Biol Evol 32, 268–74 (2015).

71. Boissinot, S., Chevret, P. & Furano, A.V. L1 (LINE-1) retrotransposon evolution and amplification in recent human history. Mol Biol Evol 17, 915–28 (2000).

72. Nik-Zainal, S. et al. The life history of 21 breast cancers. Cell 149, 994–1007 (2012).

