## Supplementary text, figures and tables for "Synchronous L1 retrotransposition events promote chromosomal crossover early in human tumorigenesis": Zumalave&Tubio_STEXT_20240530_SUBMITTED copy.docx

1. The landscape of cancer retrotransposition in the light of long reads
   1.1. Screening of high-retrotransposition rate tumours

We performed shallow sequencing with Illumina paired-ends, 350 bp insert size and 150 bp read length to obtain the whole genomes of 137 tumours, including head-and-neck squamous carcinoma (n = 37), lung squamous carcinoma (n = 50) and colorectal adenocarcinoma (n = 50). This approach resulted in an average coverage of 11.7x (95% confidence interval, ranging from 11.5x to 11.9x; **Supplementary Fig 1a-b**). Alignments to the reference human genome (assembly GRCh38) were performed using bwa mem v0.7.17 ^1^ and processed with samtools v1.12 ^2^. Duplicate reads were identified and removed from subsequent analyses using biobambam2 v2.0.87 ^3^. The resulting BAM files were then analysed using xTea ^4^ for detecting L1 retrotransposition events. To confidently identify somatic events, given the absence of genomic data from adjacent tissues at this stage, we genotyped the resulting L1 insertions using MEIGA-SR (i.e., MEIGA-Short Reads, described in section 5.1) and excluded those events shared by two or more donors within the tumoral cohort. Additionally, we excluded events documented in the database of retrotransposon polymorphisms generated within the framework of the 1000 Genomes Project ^5^. We then selected 10 tumours with a minimum of 100 somatic retrotranspositions for multiplatform whole-genome sequencing with short and long reads. See **Supplementary Table 1**.

1.2. Whole-genome sequencing of high-retrotransposition tumours with short and long reads

To achieve a comprehensive genomic characterization of the 10 tumours with high retrotranspositions rates selected in the screening stage, we performed whole-genome sequencing of these tumours and their matched-normal (non-tumoral adjacent tissues) counterparts with short (Illumina) and long (Oxford Nanopore Technologies, ONT) reads. For Illumina, we carried out paired-end libraries with a 350 bp insert size and 150 bp reads, each to a final coverage of ~30x. Alignments to the reference human genome and marking of read duplicates were performed as described above. For single-molecule sequencing with ONT, we used the Short Read Eliminator XS buffer (Circulomics, Maryland, USA) to remove DNA fragments shorter than 5 kb. The DNA was then purified using Agencourt AMPure XP magnetic beads (Beckman Coulter, California, USA), following the manufacturer’s instructions. Libraries were constructed using the Oxford Nanopore Sequencing ligation library preparation kit (SQK-LSK109, Oxford Nanopore Technologies Ltd) according to the manufacturer’s protocol, including an initial DNA end-repair and dA-tailing step using the NEBNext End Repair/dA-tailing module (NEB). Then, libraries were loaded into MinION R9.4 flow cells (FLO-MIN106, Oxford Nanopore Technologies Ltd) for sequencing. MinION devices were controlled by the MinKNOW software (v21.02.1 to v22.10.10). High-accuracy base calling was performed using the GPU-dependent Guppy software v6.1.5 (Model: dna_r9.4.1_450bps_hac). The resulting fastq files were aligned to the reference genome (GRCh38) with minimap2 v2.24 ^6^, and the generated alignments underwent sorting and quality-based filtering with samtools v1.12. All the tumours were sequenced to a minimum coverage of 30x and resulted in a median N50 of 19.9 kb (Range = 13.9-24.7, **Supplementary Fig. 1c-d**; **Supplementary Table 5**). Additionally, the tumour with the highest rate of somatic retrotransposition (PD0270a) was further sequenced using an alternative long-read platform, namely PacBio HiFi, alongside its normal adjacent tissue. The sequencing coverage was 43x and 19x, with an N50 read length of 18.3 kb and 15.3 kb, for the tumour and normal adjacent tissue, respectively.

1.3. Identification of somatic retrotranspositions with MEIGA

To find and characterize somatic retrotransposition in the long reads’ dataset, we developed the bioinformatic algorithm MEIGA: Mobile Element Integration Genome Analyzer. The method hinges on identifying read clusters indicative of structural variation breakpoints (**Supplementary Fig- 2a-b**), followed by reconstructing these variants through local assembly to precisely identify hallmarks of somatic retrotransposition. MEIGA can discern eight primary types of somatic retrotranspositions based on the transposed sequence: solo-retrotranspositions encompassing full or partial sequences of L1, Alu and SVA; partnered transductions, where an L1 sequence together with a non-repetitive downstream sequence of an active L1 retrotransposition are retrotransposed; orphan transductions, involving only the non-repetitive downstream sequence without the cognate L1; processed pseudogenes, comprising events where mRNA from nuclear genes is retrotransposed; and solitary poly-adenylate [poly(A)] tracts from severely truncated retrotranspositions. Depending on the type of structural variation generated by the integration process, MEIGA can identify insertions resulting from canonical integration events, as well as other types of rearrangements resulting from aberrant integrations.

MEIGA can be run in paired (tumour-normal) mode to detect variants acquired somatically. Input files are two BAM files (tumour and normal) containing aligned reads on a reference genome, specifically generated from long-read sequencing platforms like ONT and PacBio. The pipeline is tailored to work using the GRCh38 human reference assembly, although it can be adapted for different genome assemblies if repeats’ annotations are available. The MEIGA workflow involves five steps for accurately identifying retrotransposition events, as follows. Note that most of the parameters mentioned below are “default” variables that can be adjusted by the user.

**a) Recruitment of supporting reads**

MEIGA starts by identifying reads that provide evidence for various types of SVs relative to the reference genome. For this purpose, MEIGA screens the CIGAR strings of BAM alignments and distinguishes between two types of reads (**Supplementary Fig- 2a-b**): (1) “spanning reads”, which are defined as those reads that cover the full size of an insertion or SV, and (2) “split reads”, which are those that cover one of the two breakpoints delimiting an insertion or other type of SV. In the split reads, two alignment segments are identified: the anchor segment, which aligns onto the region of the reference genome immediately adjacent to the relevant SV and the clipped segment, which supports either an insertion larger than the clipping length or another type of SV. To ensure the precision of MEIGA calls, using default parameters, only read alignments with a mapping quality (MAPQ) higher than 20 are considered. The algorithm is preconfigured by default to identify SVs larger than 50 bp, which helps to reduce the error rate associated with the loss of base-calling accuracy typically observed for short indels in ONT data. When MEIGA is executed in paired mode, recruitment of supporting reads is simultaneously performed in both the tumour and matched-normal sequencing data.

**b) Read clustering**

The read clustering process involves two steps, namely primary clustering and meta-clustering. In the primary clustering, split reads are grouped to conform a “split clusters” of at least two reads, which must accomplish two conditions: first, their anchor segments share the same clipping orientation; second, their distance to the closest breakpoint within the cluster is ≤50 bp. Similarly, spanning reads are clustered together to conform a “spanning cluster” if their embedded insertions are closer than 250 bp. Additionally, spanning clusters undergo a correction step to fix insertion fragmentation caused by alignment contiguity defects. This correction process includes merging shorter insertions and redefining insertion boundaries for supporting reads displaying insertion fragmentation. In a second step, split clusters and spanning clusters undergo a meta-clustering process with the idea of grouping those primary clusters supporting the same SV. To conform a metacluster, it is required a reciprocal distance of 500 bp between clusters and a minimum of three supporting reads in total. Each metacluster is defined by two genomic coordinates that demarcate the beginning and the end of the genomic region encompassed by that metacluster. The meta-clustering process yields two types of metaclusters: insertion (INS) and break-ends (BND) metaclusters. INS metaclusters provide support for genomic insertions entirely encompassed by the reads supporting the metacluster, whereas BND metaclusters offer support for either long insertions not fully covered by those reads, or an alternative type of SV. At this step, a set of default filters is applied to refine the metacluster calls, which include (i) setting a maximum limit of 500 reads that can compose a metacluster, and (ii) excluding metaclusters with varying insertion lengths (i.e., coefficient of variation greater than 40%).

**c) Identifying retrotransposon insertions**

From each INS metacluster, MEIGA extracts the nucleotide sequence of the relevant insertion, along with the 2.5 kb-long nucleotide sequences from the flanking regions immediately upstream and downstream. Subsequently, MEIGA employs wtdbg2 (v2.5) ^7^ to derive a consensus from the extracted sequences, which is then polished using racon (v1.5.0) ^8^. If consensus construction fails, MEIGA restarts the polishing step by utilizing the inserted sequence from the read containing the insertion whose length is the closest to the median. Next, to precisely define the insertion sequence and the insertion-genome junctions, the polished sequence is aligned to the reference region where the insertion was located using minimap2 (v 2.24). In a subsequent step, MEIGA aims to determine the identity (i.e., the retrotransposition type) of the insertions by performing a whole-genome alignment, followed by an annotation step. For the whole-genome alignment, MEIGA maps the predicted inserted sequences using both bwa-mem and minimap2. We employ this dual approach because bwa-mem excels at characterizing short alignments that may be missed by minimap2, while minimap2 is better suited for handling longer alignments that may become fragmented when using bwa-mem. Next, MEIGA inspects the genomic environment linked to the alignment positions retrieved for the inserted sequences, to identify and annotate relevant genomic features, including retrotransposon-like repeats, poly(A/T) repeats, sequences positioned downstream of source L1 elements, which may serve as indicators of transductions and coding regions, which may point to pseudogene mobilizations. Only INS metaclusters that fall into one of the mentioned categories are retained for further steps.

**d) Identifying retrotransposition-mediated junctions**

BND metaclusters can be suggestive of rearrangements mediated by retrotransposition, which are presented as junctions between two distant breakpoints in the genome, with retrotransposition bridges connecting them. These retrotransposition bridges are expected to be supported by a single BND metacluster at each genomic breakpoint. To identify these events, MEIGA first examines the longest clipping that supports each metacluster and performs a whole-genome alignment using minimap2. Subsequently, our algorithm inspects the alignments adjacent to the beginning of the clipping segment to identify the presence of a retrotransposition bridge, by using annotations mentioned in the previous step. Only BND metaclusters in which a candidate bridge is identified are retained for further analysis. Then, MEIGA continues to inspect the alignments next to the candidate bridge – referred to as anchor alignments – with the idea of finding pairs of BND metaclusters in which the coordinates of one metacluster overlap with the coordinates of the anchor alignments from the other metacluster. When these metaclusters are identified, they are classified as junctions and are retained for further analysis if they satisfy the following two conditions: (i) sharing a minimum of two reads, and (ii) their respective bridges exhibit the same identities. Junctions can be: intrachromosomal, which are often associated to DNA loss or gain; or interchromosomal, suggestive of a translocation event. Finally, MEIGA aims to accurately reconstruct the nucleotide sequence from each junction’s bridge, following an approach similar to that described for the INS metaclusters, as follows. First, MEIGA gathers the supporting reads and extracts bridge sequences together with up to 2.5 kb of the flanking regions. Next, using wtdbg2, MEIGA creates a consensus sequence, which is further refined with racon. If consensus construction is unsuccessful, the process begins by extracting the bridge sequence from the event that has both the longest clipped and anchor segments. Then, MEIGA accurately determines the breakpoints of the bridge by executing a minimap2 alignment targeting both genomic sides of the junction.

**e) Refining insertion and bridge structures**

To acquire a more detailed understanding of the structural features of candidate retrotransposition events, MEIGA employs minimap2 to align the nucleotide sequences of insertions and junctions’ bridges with the consensus sequence of the candidate retrotransposon family that has been preassigned for each event. This approach provides a precise means of assessing various structural attributes, such as retrotransposon subfamily, internal rearrangements (e.g., inversions and deletions), event size, presence of poly(A/T) tail, insertion mechanism and other relevant characteristics. Then, MEIGA conducts an additional read gathering step with less stringent criteria to refine the number of supporting reads. Subsequently, candidate retrotransposition events are categorized as somatic when no supporting reads are found in the matched normal sample. Otherwise, they are classified as germline. Finally, MEIGA proceeds to annotate the genomic regions where the retrotransposition events are located, which is required for the final filtering step.

**f) Applying filtering criteria**

After full characterization of the retrotransposon sequence, a final set of filters is applied to ensure specificity. Candidate insertions and junctions are retained if the following conditions are met: (i) the minimum and maximum number of supporting reads ranges from 3 to 500 reads, (ii) the insertion sequence is not entirely composed of expanded repeats/low-complexity sequences, except for poly(A/T) tracks, (iii) there are no ambiguous retrotransposon families assigned to the same insertion, (iv) the proportion of the insertion sequence with an assigned identity exceeds 40%, and (v) the ratio of spanning reads versus clipping reads is consistent given the insertion size. Finally, retrotransposon insertions are reported in VCF format.

1.4. MEIGA benchmarking

To establish a robust benchmarking framework for MEIGA, we developed MEIsimulator, a bioinformatics tool designed to generate sequencing data from synthetic genomes that incorporate a wide range of randomly integrated retrotransposition events. Briefly, the algorithm starts by seeding retrotransposon sequences at randomly predetermined positions within a reference genome to generate a custom mock genome and subsequently employs ART v2.5.8 ^9^ or PBSIM v1.0.4 ^10^ to generate, respectively, Illumina short reads or ONT long reads on the new genome configuration. MEIsimulator can also simulate aberrant retrotranspositions that result in SVs such as deletions, duplications, inversions and translocations.

To evaluate MEIGA, we used MEIsimulator with default settings to simulate a total of 6,420 retrotransposon insertions, including 1,000 Alu-solo; 1,000 SVA-solo; 1,000 L1-solo; 1,710 L1-partnered and 1,710 L1-orphan. Specifically, for each one of the 114 L1 source elements found to be active in the PCAWG dataset ^11^, we simulated 15 L1-partnered and 15 L1-orphan transduction events (**Supplementary Table 6**). Additionally, we conducted simulations of L1-mediated genomic rearrangements, which included 20 deletions, 20 duplications, 20 inversions and 10 translocations (**Supplementary Table 7**). These sets of retrotransposition events were replicated in both short-read and long-read sequencing datasets. In the short-read simulation, we set a sequencing depth of 30x and a VAF of 50%. For the long-read simulation, retrotransposition events were simulated at different VAF levels, including 0.15, 0.3 and 0,5 and to a sequencing depth of 33.7x, which corresponds to the median depth of our ONT cohort. We then run MEIGA together with three additional reference pipelines to detect retrotranspositions in these mock genomes, including rMETL v1.0.4 ^12^, xTea v0.1.9 ^4^ and PALMER v2.0.0 ^13^ Each caller was executed using default parameters to ensure a consistent evaluation. Key metrics such as precision, recall and computational efficiency were assessed to evaluate the performance of each tool. Precision metrics did not account for retrotransposition-type assignment, as it was not specified by all callers, and a 150 bp offset was allowed for intersecting the calls. The results showed a higher precision and recall of MEIGA for overall insertions (f1-score ranges from 83.70 to 99.55 for VAFs from 0.15 to 0.5, respectively) relative to the other three pipelines employed (**Fig. 1b**; **Supplementary Fig. 2c-d**; **Supplementary Tables 8-10**).

In addition, we used Blast v2.11.0 ^14^ to evaluate the performance of MEIGA to reconstruct the nucleotide sequences of the simulated insertions from above, by analysing the size and the nucleotide identity between observed and expected sequences. Here, our algorithm successfully reconstructed the size of the simulated insertions, showing an average observed to expected size ratio O/E = 0.99 and a pairwise average nucleotide identity ANI = 99.93% (**Supplementary Fig. 2e**).

1.5. Validation of MEIGA retrotransposition calls

We ran MEIGA on the long-read sequencing data from the 10 relevant tumours and their adjacent tissues. The analysis retrieved a total of 6,418 retrotranspositions acquired somatically and confirmed that all tumours have high rates of somatic retrotransposition (median = 622.5, range [296-1,311]). To validate our results, we carried out further sequencing in the tumour with the highest number of canonical somatic retrotranspositions (PD0270a, n = 1,290; note that are excluding 21 retrotransposition-mediated rearrangements at this step), including two additional technologies (Illumina paired-ends and PacBio HiFi). Then, we analysed somatic retrotransposition with other relevant pipelines, including xTea-SR for the Illumina short reads and xTea for the ONT and PacBio long reads. The intersection of the insertions called by each method is shown in **Supplementary Fig. 3a** and **Supplementary Table 11**. Of note, since xTea does not support somatic calling, xTea private calls have been excluded from the analysis because most corresponded to germline variants. Our results find that 92.5% (1,193/1,290) of MEIGA calls for ONT, are validated by at least one external method (xTea-SR or xTea) or an additional technology (Illumina paired-ends or PacBio HiFi). Furthermore, we find that 91.6% (1,183/1,292) of MEIGA calls for PacBio are also validated, indicating a good performance of MEIGA for both ONT and PacBio HiFi datasets. Of note, to validate the fraction of MEIGA private calls (i.e., those events that do not intersect with any other pipeline nor technology; accounting for 97 calls on the ONT dataset, 109 calls on the PB dataset and 112 calls on both ONT and PB), we carried out visual inspection with Integrative Genomics Viewer (IGV) of a random selection of 150 insertions (50 MEIGA private calls from the ONT dataset, 50 private calls from the PB dataset and 50 private calls from the ONT+PB dataset). The following criteria had to be accomplished for a candidate insertion to be classified as a real somatic retrotransposition: (i) presence of three or more supporting reads in the tumour long-read data; (ii) lack of any supporting reads in the matched normal long-read data; (iii) presence of at least one of these retrotransposition hallmarks: TSD, poly(A/T) tail or retrotransposon sequence. This analysis confirmed a true positive rate of 94.7%. (142/150) (**Supplementary Fig. 3b**). Overall, our analysis reveals that long reads provide ~32.4% more somatic insertion events than with short reads only (**Supplementary Fig. 3c**). Finally, to evaluate the relative sensitivity of both long-read methods employed here, MEIGA and xTea, we estimated the fraction of somatic events that those two methods can recall from a dataset of 943 somatic calls with short reads. We observe that MEIGA can call ~12.9% and 12.8% more somatic events than xTea on the ONT and PacBio datasets, respectively (**Supplementary Fig. 3d**).

2. Long reads reveal the patterns and mechanisms of L1 internal rearrangements
   2.1. Annotation of the internal structure of L1 insertions

Somatic L1 insertion calls are subjected to an additional annotation procedure to analyse their structural configuration. First, the sequence corresponding to the target site duplication was detected and trimmed from either the 5’ or 3’ end of the L1 insert through the identification of exact matches with the genomic sequence at the integration position. Then, to have all L1 inserts in forward (5’ to 3’) orientation, the reverse complement sequence for every L1 insertion occurring in the minus strand was obtained. Then, poly(A) tails were detected for every insertion event, requiring poly(A) monomers to have a minimum size of 15 bp and a minimum purity of 90%, and be located at a maximum distance of 30 bp relative to the insertion end. L1 insertions containing a single poly(A) tract were classified as solo events, while those containing two tracts were annotated as 3’ transduction candidates with the sequence in between the poly(A) stretches corresponding to the transduced bit of DNA. To trace transductions to their full-length L1 progenitor, transduced sequences were aligned with bwa-mem v0.7.17-r1188 to a patient specific database containing the sequences corresponding to the 10 kb genomic intervals downstream to each of the full-length L1s (germline plus somatic) detected for a given patient. To maximize sensitivity for particularly short transduction events a minimum seed length (*-k*) of 8 bp and a minimum score (*-T*) of 0 were used. Alignment hits were further filtered by requiring a minimum mapping quality of 20. The inserted sequences were then further trimmed by removing the poly(A) tails and transduced sequences, resulting in trimmed sequences corresponding to L1 sequences alone. Trimmed inserts were aligned using bwa-mem (same parameters mentioned above) into a consensus L1 sequence derived from the 632 FL-L1 insertions included in the HGSVC call set ^5^. Alignment hits over the L1 consensus were chained based on complementarity to identify the minimum set of nonoverlapping alignments that span the maximum percentage of the inserted sequence. Based on the alignment chains, L1s are classified as full-length (single hit spanning >95% of the consensus L1), 5' truncated (single hit spanning ≤95% of the consensus L1), 5' inverted or twin priming (two hits with the first in reverse while the second in forward orientation), interstitial deleted (two hits in forward orientation with the distance in between corresponding to the deletion length) and complex (more than 2 hits; of note, all complex insertions were visually inspected and manually reannotated). Then, the inversion junction conformation for every twin priming event is determined based on the alignment position over the consensus for the inverted and non-inverted L1 pieces. Blunt joints are characterized by perfect complementary alignments, while overlapping and discontinuous alignments define duplications and deletions at joints, respectively.

2.2. The insides of L1-mediated transductions

Among the many ways in which L1 activity has shaped the human genome is the mobilization of unique DNA sequences downstream of the L1 element, a process known as L1-mediated 3’-transduction. In our cohort, MEIGA identified 1,535 partnered and 705 orphan transductions with a median size of 1,093 bp (range = 112-7,421 bp) and 464 bp (range = 56-3,861 bp), respectively (**Fig. 2a**). The median size of the transduced regions is 0.1 kb and 0.4 kb, respectively for partnered and orphan transductions (**Supplementary Fig. 4d**). As for the internal rearrangements of L1-partnered transductions, we find ~0.4% (6/1,535) insertions in which the companion L1 is full-length element likely competent for retrotransposition; 37% (570/1,535) of events have a 5’-deletion only; and 56% (865/1,535) events bear a 5’ deletion together with a 5’ inversion. See details in **Supplementary Fig. 4e**.

In our cohort, L1-transductions represent 38% from the total L1-derived somatic insertions, which represents a relevant increment relative to previous cancer retrotransposition studies that reported a transduction frequency of 19%-24% using paired-ends ^15,16^. We observe that this discrepancy is due to the existence of partnered transductions bearing transduced regions shorter than ~50 bp, which were misinterpreted as solo-L1s in previous retrotransposition analyses. In our cohort, these “microtransductions” have a median size of 24 bp (range = 10-49 bp) and represent 28% (627/2,240) of all the L1-mediated transductions (**Supplementary Fig. 4f**). Some microtransductions are particularly frequent, which reveals the activity of specific L1 loci. In one relevant example, a germline L1 element located at chromosome Xp22.2 gave rise to 537 microtransductions 15-39 bp-long (median = 24 bp) region downstream. Another L1 element located at chromosome 6p24.1 promoted 168 microtransductions that are 14 to 47 bp-long (median = 23 bp).

2.3. The insides of processed pseudogenes

Our method identified 133 insertions of processed pseudogenes. These are by-products of L1 activity generated when the mRNA molecules from nuclear genes are retrotranscribed and integrated using the L1 protein machinery. The structural analysis shows that median length of the pseudogene insertions in the cohort is 574 bp, ranging from 96 to 3,646 bp (**Fig. 2a**). In a remarkable example from tumour PD0307a, we found the largest pseudogene insertion: a 3,646 bp-long event containing 15 different exons from the gene *SIPA1L2*. Overall, we find that 4.5% (6/133) are full-length insertions that keep all the coding exons and UTRs and preserve their full structure. The remaining 127 events present a structure that is truncated by 5’-deletions of different size: 10% (13/127) of them represent “complete” insertions that retain all the coding exons and UTRs but exhibit short 5’-deletions in the 5’-UTR; 35% (44/127) are insertions that preserve the 3’-UTR together with one or more coding exons (median of retained coding exons is 2, range = 1-14); and a majority (54%, 68/127) of insertions are represented by a solitary 3’-UTR with no companion coding exon. 24% (31/127) of the events with 5´-deletion presented co-occurrence of inversion of the 5´-extreme, indicating that one-quarter of the pseudogene insertions followed twin priming retrotransposition ^17^, of which most have nucleotides at the point of inversion that are either deleted (58%, 18/31) or duplicated (35%, 11/31), and two of them show both deletion and duplication. Here, duplications have a median size of 9 nucleotides (range = 1-28) and deletions of 12.5 (range = 1-632). See **Supplementary Fig. 4g-h**.

3. A novel panorama of source elements activity in the light of long reads
   3.1. Haplotype phasing of long reads

We carried out the phasing of germline polymorphisms and reconstructed the parental haplotypes from long reads, which will be later employed in the characterization of L1 source candidates (see next section). First, we run Freebayes v1.3.6 [arXiv:1207.3907] on the Illumina short-read sequencing data obtained from matched-normal samples to identify germline polymorphisms (SNPs and short indels). Then, we run WhatsHap v1.2.1 [doi:10.1101/085050] on the ONT long-read sequencing data to first construct a graph that captures the different ways the germline variants could be combined, and then performed phasing inference by assigning the most likely haplotype to each parental chromosome. In a final step, we employed WhatsHap to add phasing tags to each alignment that could be confidently associated with either parental haplotype. This step was performed on the alignments included in the ONT long-read sequencing data derived from both the tumour and matched normal tissues. Of note, the phasing strategy obtained haplotype blocks larger than 1.5 Mb for both the adjacent tissues and the tumours (**Supplementary Fig. 1e-f**).

3.2. Identification of L1 source candidates

We aimed to identify all potentially active L1 loci in the cohort achieving haplotype-level resolution, from a dataset of 693 L1 candidates that have a complete length and structure initially described in PCAWG ^16^ and HGSVC2 ^5^. To accomplish this, we used the ONT phased BAM files obtained from matched-normal samples. We employed two different approaches depending on whether the source elements were present or absent on the reference genome. First, for the L1 elements present in the reference genome, reconstruction of the entire sequence was performed using the ‘*refSrc_seqs.py*’ algorithm included in the MEIGA suite. In brief, we first collected alignments that spanned an L1 element and its adjacent regions. These reads were then segregated based on their haplotype tags and mapped to the L1Hs consensus sequence. For each distinct haplotype, the resulting alignments were aggregated and assembled using wtdbg2 (v2.5). Subsequently, the assembled sequences underwent a polishing step with racon (v1.5.0) and were finally realigned to the L1Hs consensus to identify the precise breakpoints of the source L1 element. Second, for the non-reference source L1 elements, we adopted a distinct approach. As these elements manifest as insertions in the genome, we employed MEIGA, using default parameters, to identify full-length solo L1 insertions in the matched-normal tissues. In cases of homozygous non-reference L1 loci, we re-ran MEIGA on the target regions employing the *‘--hpTag’* parameter for the reconstruction of haplotype-specific sequences. For heterozygous non-reference L1s, we relied on the insertion sequences initially reconstructed by MEIGA. Using the haplotype-aware version of MEIGA could have resulted in reduced sensitivity, due to the exclusion of reads where haplotype assignment is uncertain.

3.3. The Source Inference method

Our source inference method elucidates the element-of-origin of somatic L1s through the identification of specific SNVs shared between these L1 sources and their derived insertions. The method starts by genotyping all germinal full-length L1 loci within each patient’s normal genome to identify the diagnostic SNVs that characterize each L1 source. Then, the algorithm identifies SNVs in the repertoire of L1 somatic copies in the patient’s tumour genome. Finally, the element-of-origin of each individual somatic copy is inferred by comparing the set of SNVs present in each somatic copy with the set of diagnostic SNVs that characterize each source L1. The algorithm is available at <https://gitlab.com/mobilegenomesgroup/SourceInference>. The workflow is as follows:

**a) Identification of diagnostic nucleotides in L1 germline loci**

All candidate L1 loci retrieved from the genotyping of the matched normal genome are aligned using minimap2 v2.24 ^18^ to the L1HS consensus sequence with NCBI accession number L19088.1. SNVs were then called from the resulting alignment using Rsamtools v2.10.0 pileup (R package version 2.18.0, https://bioconductor.org/packages/Rsamtools). Diagnostic SNVs are defined here as those private to a specific L1 locus, meaning they were not present in any other source element locus within the same donor. SNVs shared by several candidate elements were classified as “regular”. Source elements exhibiting more than 1% sequence divergence to the L1 reference sequence, as determined by the detected SNVs, were excluded from analysis for that specific patient. If a given source element fails to meet this criterion in two or more donors, it is filtered out from the entire cohort.

**b) Identification of SNVs in somatic L1s**

The identification of SNVs on L1 somatic insertions involves an iterative pairwise alignment to the reference L1 sequence by means of the Smith-Waterman algorithm as implemented in Biostrings v2.62.0 R package. To address somatic insertions containing internal rearrangements, we first reconfigured them to their primary orientation before alignment to ensure accurate mapping. Then, SNV calling was performed individually for each L1 insertion using the mismatchTable function within Biostrings.

**c) Inference of source elements**

Each individual somatic L1 insertion is compared to the repertoire of candidate source elements identified in the matched-normal genome, with the aim of identifying regular and diagnostic SNVs between them and using this information to assign a somatic L1 to a specific source element. The following conditions must be accomplished: (1) there is a 100% of diagnostic variants shared between the somatic insertion and the potential element-of-origin; (2) there is no other candidate source element sharing 100% of diagnostic variants with the same insertion; (3) There is a ≥75% of total SNVs (diagnostic + regular) shared between the relevant somatic insertion and the candidate source element.

We run Source Inference on the 10 tumours from our cohort. The algorithm identified 46 cryptic elements that emerged as donors of somatic solo-L1s using this alternative strategy that were not identified with a transduction-based approach. This makes a total of 197 source elements (46 elements obtained with the diagnostic nucleotides strategy and 151 with the transduction-based approach) with proven activity in the cohort. Overall, our algorithm attributed 899 somatic L1-solos to a particular donor element, an addition that had a substantial impact in the patterns of source element activity (**Supplementary Table 4**; **Supplementary Fig. 6a-b**).

3.4. Evaluation of the Source Inference method

To evaluate our method, we simulated the mobilization of all potential source elements (n = 266) from the patient in the cohort with the highest L1 activity rate (PD0270), assuming a homogeneous activity of 20 retrotranspositions for each one of the 266 elements and with the derived copies showing the same insertion size distribution as in 2954 cancer genomes from the PCAWG project ^16^. The results showed a high specificity of the pipeline (>99%) and indicated an overall sensitivity of 47.6% (2,530/5,320), with a drop in sensitivity when somatic L1s are derived from younger source elements in relation with a decrease of genetic divergence (sensitivity = 59.2% [1042/1,660] for pre-Ta; 47.0% [583/1,240] for Ta-0; 36.8% [735/2,000] for Ta-1). We also observe that the performance of the method is notably affected by the size of the derived copies, with a reduction in the sensitivity inferred from smaller derived copies, particularly in those copies derived from Ta-1 source elements (**Supplementary Fig. 5**).

To further evaluate our method, we employed a set of 388 partnered transductions that were confidently preassigned to its source element of origin to analyse the concordance between source element inference derived from transductions and from L1 diagnostic nucleotides. These partnered transductions represent retrotranspositions that preserve both the L1 sequence and the downstream transduced region, making them a close-to-ideal scenario to test our algorithm. This analysis shows that 356 out of 388 partnered transductions were assigned to the same source element using both methods in an independent manner, being 91.7% the concordance between both strategies across the ten samples. We observed, however, that concordance varies with the length of the interrogated insertions, dropping off to 86% (117/136) when the analysis is restricted to partnered transductions with transduced regions below 200 bp, and increasing up to 94.8% (239/252) when delimiting the analysis to partnered transductions with transduced regions of 200 bp or longer.

3.5. Identification of hot L1 elements and L1 phylogenetic analysis

Hot L1 elements must meet a minimum of one of the following conditions: (i) L1 activity rate, defined as the average number of mobilizations mediated by a source element when found activated) is equal or more than 10; (ii) the relative contribution to the total amount of traced mobilizations in the cohort is equal or more than 1.5%. This analysis finds 27 hot L1 germline in our cohort (**Supplementary Fig. 6c**). These hot sources were used to build a L1 phylogeny, as follows. Consensus sequences from 27 L1 hot elements and L1Pt (L1 *Pan troglodytes*) sequence [GenBank: KF661301.1] were aligned with muscle v3.36.0 ^19^. The L1 phylogenetic tree was built using the following R packages: Ape v5.6-2 ^20^ and Phangorn v2.8.1 ^21^. Nucleotide substitution model selection was performed with phangorn::modelTest ^22^. GRT + G + I model was selected based on AIC (Akaike Information Criterion) score. A maximum likelihood tree was estimated with the phangorn::optim.pml function, starting from a distance-based tree. Tree topology was adjusted via nearest neighbour interchange ^23^. After maximum likelihood optimization, midpoint rooting was applied, rendering L1Pt as the expected tree outgroup. Non-parametric bootstrap analysis with 1000 replicates was done to determine the node consistency along the inferred tree. For display purposes, only values informative of high support (≥80%) were labelled to the tree nodes.

L1Hs subfamily assignment was performed through the identification of subfamily diagnostic nucleotide positions on their 3’ end of the L1 sequence ^24^. FL-L1 source elements bearing the diagnostic “ACG” or “ACA” triplet at 5,929-5,931 position were classified as “pre-Ta” and “Ta”, respectively. Ta elements were subclassified into “Ta-0” or “Ta-1" according to diagnostic nucleotides at 5,535 and 5,538 positions (Ta-0: G and C; Ta-1: T and G). Source elements that did not display any of these diagnostic profiles could not be assigned to a particular category, and their subfamily status remained undetermined.

3.6. Analysis of methylation profiles

Estimates of methylation content for all CpG sites were calculated with Megalodon (v2.5.0) ^25^ using the Remora neuronal network model dna_r9.4.1_e8. The configuration for the remora-modified-bases was set with the flags: ‘dna_r9.4.1_e8 hac 0.0.0 5hmc_5mc CG 0’. Epigenetic profiles for individual L1 loci were established by averaging the DNA methylation levels across CpG dinucleotides within the bodies, promoters and adjacent genomic regions, spanning 20 kb upstream and downstream, using BEDtools v2.31.027. Disparities between tumour samples and their corresponding adjacent healthy tissues were quantified using Gtools v3.9.428, and statistical significance was determined employing two-sided Mann-Whitney-Wilcoxon tests. See **Supplementary Fig. 7**. See **Supplementary Tables 12 and 13**.

3.7. Detection of polyadenylation signals downstream L1 loci

APARENT ^26^ is a deep neural network designed to detect and score all polyadenylation sites within a given DNA sequence. In this study, we employed this software to identify both canonical and alternative polyadenylation signals and to estimate their relative strengths. For each tumour sample, APARENT was applied to the reconstructed sequences of source L1 elements along with their unique downstream sequences. Subsequently, we matched this information with the rates of solo insertions and transductions (**Fig. 3d-e**).

4. A hidden landscape of genomic rearrangements mediated by retrotransposition
   4.1. Identification of retrotransposon-mediated rearrangements

Rearrangements mediated by retrotransposition manifest as junctions between two distant breakpoints in the genome with retrotransposon bridges connecting them. MEIGA identified a total of 152 retrotransposition-mediated junctions in the 10 tumours cohort. These junctions were classified into four primary categories, determined by (i) whether they were interchromosomal or intrachromosomal events and (ii) the orientation of their breakpoints (note that we have not considered associated copy-number changes in this classification). For intrachromosomal junctions, our naming convention was as follows: deletion-like for [+/-], duplication-like for [-/+] and inversion-like, which encompasses both head-to-head [+/+] and tail-to-tail [-/-] inversions. Interchromosomal junctions were categorized as translocation-like.

Deletions-like junctions were the most prevalent category, comprising 43% (n = 66) of the total, followed by translocations-like (30%, n = 45), inversions-like (18%, n = 27) and duplications-like at (9%, n = 14). Notably, these numbers significantly differ from those reported in the PCAWG study ^11^, where only 93 retrotransposon-mediated rearrangements were reported, including 90 deletions, one duplication, one translocation and one inversion were detected across 2,954 tumours. This suggests that all categories except deletions were likely underestimated in prior short-read studies.

Our analysis shows that 90% of the deletions mediated by retrotransposition in our cohort span regions between 0.1 and 10 kb, resembling the size distribution observed for deletions in the PCAWG dataset (Wilcoxon rank sum test, *p* = 0.42). However, inversions and duplications did not exhibit such patterns **(Supplementary Fig. 8a**).

4.2. Validation of L1-mediated reciprocal translocations by cytogenetics

To validate our findings, we included an additional lung adenocarcinoma cell line, NCI-H2009, and its corresponding normal counterpart, NCI-BL2009, in our study. We conducted dual FISH experiments on the NCI-H2009 cell line to assess the occurrence of a reciprocal translocation between chromosomes 3 and 6 mediated by the somatic integration of a L1. We employed commercially available whole chromosome painting (WCP) probes targeting chromosomes 3 and 6 labelled with either green or red fluorescent dyes (FWCP-03 and FWCP-06, respectively; Creative Bioarray). Metaphase spreads were obtained from NCI-H2009 cells following standard techniques, pre-treated with RNAse and pepsin and subjected to dual FISH experiments as per the provider’s recommendations. After a counterstaining with DAPI (0.14 pg/ml), photographs were taken for each individual channel with a Nikon Eclipse-800 fluorescence microscope (Tokyo, Japan) equipped with a DS-Qi1Mc CCD camera (Nikon) and controlled the using NIS-Elements software (Nikon). The resulting images were merged and processed using Adobe Photoshop CS6 (San Jose, CA, USA).

4.3. Validation of L1-mediated reciprocal translocations by chromatin conformation capture

We carried out Micro-C to detect 3D chromatin contacts at a genome-wide level in the high-retrotransposition rate cancer cell line NCI-H2009 ^15^. Micro-C libraries were prepared following the Dovetail Micro-C Kit protocol (Dovetail Genomics). Sequencing of the libraries was performed at 150 bp paired-end by an external service (Macrogen Inc.) using the HiSeqX sequencer from Illumina. The Micro-C library was then aligned to the GRCh38 genome assembly using bwa-mem v0.7.17. Following the recommended Micro-C procedure, we used the parse module of pairtools v0.3.0 [doi: https://doi.org/10.1101/2023.02.13.528389] to identify ligation junctions in our libraries. Samtools v1.14 was employed to sort the parsed pairs and pairtools dedup was used to mark PCR duplicates, which were subsequently excluded from downstream analysis. Finally, a 5 kb resolution contact map was generated using cooler v0.8.11 ^27^. See **Supplementary Fig. 8b**.

5. The tempo of somatic retrotransposition in human cancer

Current computational approaches for timing inference rely on the estimation of the variant allele frequency (VAF) and the local copy number state from short reads to classify variants as early or late events ^28,29^. We adapted these timing pipelines to deal with canonical retrotranspositions and retrotransposon-mediated rearrangements. Briefly, we first developed an algorithm specifically designed for VAF estimation of retrotransposition events from short reads; then our timing approach employed the estimated VAF, along with tumour purity and copy number information estimated from Battenberg ^30^, to obtain a relative timing estimation of somatic retrotranspositions.

5.1. Analysis of retrotransposition VAFs using MEIGA-SR

While tools like SVclone ^31^ are available for inferring allele frequencies of classical SVs in short-read sequencing data, they encounter limitations when dealing with retrotransposition events. Unlike classical SVs, retrotransposon insertions do not exhibit the characteristic formation of two well-defined clusters of discordant pairs—one on each side of the SV junction. Instead, these events are supported by discordant pairs where one read of the pair accurately clusters near the insertion breakpoint, while their mates disperse throughout the genome, mapping to multiple locations where a retrotransposon of the same class is present. Additionally, as reference reads precisely match the reference genome, they inherently possess a higher likelihood of alignment compared to alternate reads supporting an SV. Consequently, when computing allele frequencies of retrotransposition events, there is a risk for reference reads to be overrepresented. Here, we leveraged the coding developed for MEIGA and developed a targeted genotyping strategy, named MEIGA-SR, to effectively address this challenge. This method was specifically designed to analyse short-read sequencing data (Illumina paired-end sequencing of 150-bp reads) and accurately estimate the VAFs of previously identified somatic retrotransposition events. The workflow involves the following three steps (see **Supplementary Fig. 10a**):

**a) Identification of Target Site Duplications (TSDs):**

MEIGA-SR begins the analysis by processing BAM/CRAM alignments from tumour samples. During this step, MEIGA-SR collects clipped reads, specifically those with soft or hard-clipped segments exceeding 5 bp, located within ±30 bp of the retrotransposition insertion. No MAPQ filter is applied at this stage but reads marked as duplicates or with clipped segments at both ends of the read are excluded. Subsequently, two alignment segments are defined within each clipped read: the anchor segment, which aligns with the region of the reference genome immediately adjacent to the candidate insertion and the clipped segment, which encompasses the inserted sequence. Furthermore, clipped reads are classified as right or left clipped, depending on the position of the clipped segments on the read. For right-clipped reads, the most common coordinate where the anchor segment ends is defined as the right breakpoint of the TSD. For left-clipped reads, the most common coordinate where the anchor segment begins is defined as the left breakpoint of the TSD. If both breakpoints of the TSD are identified—with the left breakpoint smaller than the right—and they are separated by less than the read size (e.g., 150 bp), the method will then proceed to assess the allele frequency. Otherwise, no estimation will be provided for that specific event.

**b) Recruitment of supporting reads:**

MEIGA-SR analyses read pairs within a specified interval surrounding the TSD for each event designated for genotyping. The interval size is determined by the median fragment size and the read length of the libraries used in the study (e.g., Median fragment size (m) ~450 bp; Read length (r) = 150 bp; m – r / 2 = 325 bp). In our context, MEIGA-SR defines a region that starts 325 bp upstream from the beginning of the TSD and extends to 325 bp downstream from the end of the TSD for each event. Within these regions, MEIGA-SR examines the read pairs, categorizing them as either alternate or reference supporting read pairs based on their alignment characteristics.

There are two types of read pairs that can provide support for a retrotransposition event: (i) discordant read pairs and (ii) clipped read pairs. Discordant read pairs fail to align to the reference genome within the expected distance or orientation. In the event of a retrotransposition, discordant reads form two distinct clusters on either side of the insertion site—one with forward-facing reads upstream and another with reverse-facing reads downstream—while their paired mates align to distant genomic regions containing retrotransposons from the same family. Consequently, pairs with forward-oriented discordant reads that terminate before the end of the TSD are identified as alternate supporting read pairs for the upstream cluster. Similarly, pairs with reverse-oriented discordant reads that begin after the start of the TSD are classified as alternate supporting read pairs for the downstream cluster. On the other hand, clipped read pairs are identified as alternate supporting pairs if any read in the pair features a soft or hard-clipped segment larger than 5 bp, located within ±2 bp of the defined TSD breakpoints. Right-clipped reads are classified as providing upstream support, whereas left-clipped reads are classified as providing support to the downstream cluster.

Ultimately, read pairs where neither read supports the alternate allele and both have an alignment score (AS) greater than 120 and a MAPQ exceeding 30 are classified as reference supporting read pairs. However, read pairs that begin aligning after the start of the TSD or end before its conclusion are excluded from consideration. In such cases, it becomes challenging to determine whether these read pairs support the reference or the alternate sequence. Of note, for both reference and alternate categories, read pairs marked as duplicates or with clipped segments at both ends are excluded from the analysis.

**c) Assessment of allele frequencies:**

Following the classification of read pairs, MEIGA-SR utilizes the maximum count of observed alternate read pairs within the upstream and downstream clusters to estimate the allelic frequency of each retrotransposition event. This approach is adopted due to the tendency of the 3’ end of retrotransposition events, corresponding to the poly(A) tails, to exhibit clusters with lower support. The formula used to estimate VAF of retrotransposition events is as follows:

*nb_alt = maximum(nb_upAlt, nb_downAlt)*

*nb_all = nb_ref + nb_alt*

*vaf = nb_alt / nb_all*

where *nb_upAlt* and *nb_downAlt* are the number alternate read pairs upstream and downstream the insertion, *nb_REF* is the number reference supporting read pairs and *nb_all* corresponds to the total read pairs in region after filtering. Finally, to ensure accuracy, a specific set of criteria must be met to report the VAF of an event:

i.	Alternate supporting read pairs (*nb_alt*) must be three or more.

ii.	The total number of read pairs (*nb_all*) must be equal to or greater than 20.

iii.	The average MAPQ for the region must be equal to or greater than 50.

5.2. Evaluation of MEIGA-SR

To evaluate the performance of our method, we employed MEIsimulator to generate a mock genome featuring 4,480 retrotransposon insertions representing diverse families and insertion types. This comprised 1,000 L1-solo, 500 Alu-solo, 500 SVA-solo, 1,240 L1-partnered and 1,240 L1-orphan insertions. Subsequently, Illumina paired-end reads, 150 bp-long, were simulated to achieve a coverage of 30x and at a VAF of 50%. We then applied MEIGA-SR, with two additional reference pipelines, TraFiC ^15^ and xTea ^4^, for the identification of somatic retrotranspositions. All three tools were evaluated using default parameters. The results confirmed high precision (99.9%) and recall (95.7%) of MEIGA-SR (**Supplementary Fig. 10b**). Notably, MEIGA-SR demonstrated consistent recall rates across different retrotransposon families and insertion types, ranging from 95.0% to 96.6% (**Supplementary Fig. 10c**). In addition, the analysis of identity (i.e., this is the assignment of the right retrotransposition type) showed that MEIGA-SR carried out a precise classification of most insertion types (**Supplementary Fig. 10c**), except for L1-partnered insertions, in which 15.9% (187/1,177) of the events were misclassified as L1-solo insertions. We attributed this deviation to partnered transductions with short transductions (namely microtransductions).

Similarly, to assess the accuracy of our VAF estimation approach, we used a mock cancer genome into which we seeded 6,420 retrotransposon insertions using MEIsimulator and generated short-read sequencing data to a sequencing depth of 30x and a retrotranspositions’ VAF = 0.5. Our algorithm MEIGA-SR obtained a VAF prediction for 89.6% (5,754/6,420) of the events. Analysis of these VAF estimations confirmed a normal distribution centred at the simulated value of 0.5 (**Supplementary Fig. 10d**).

5.3. Real Time WGD Timing

The whole genome duplications (WGDs) were timed using an approach similar to that outlined in ^28^. The proportion of clonal clocklike CpG > TpG SNVs on different numbers of allelic copies, known as their multiplicities, was used to infer the WGD timing. We only considered genomic regions where the highest number of copies of either parental allele, known as the major copy number, was two. This is because all gains in these regions are assumed to occur through the WGD. We assumed that a linear acceleration of 5x occurred in the CpG > TpG mutation rate at a point uniformly distributed 1 and 15 years before diagnosis. Given a sampled acceleration timepoint, we found the WGD timing that corresponded to the maximum likelihood of the proportion of multiplicity one and two clocklike SNVs in genomic region. We marginalized the likelihood over the fraction of multiplicity one mutations that were subclonal and corrected the multiplicity proportions to account for our lack of power to detect SNVs with less than three reads. The uncertainty in our WGD timing estimates was estimated by bootstrapping over the SNVs in the tumour and resampling the acceleration timepoint. See **Supplementary Table 14**.

The sequencing coverage of the samples was generally suboptimal for accurate subclonal SNV structure inference. Therefore, to account for subclonal SNVs in our timing analysis, we assumed the presence of a pseudo-SNV subclone corresponding to 30% of all biopsied tumour cells and containing 10% of detectable SNVs. As we are blind to subclonal mutations below our detection limit, our timing analysis is expected to show a slight bias towards a later WGD.

The WGD in sample PD0312 was not timed as this sample showed a modal major copy number state of greater than two, suggesting the occurrence of more than one WGD and thus unsuited to be timed by this approach.

5.4. Retrotransposition timing

Retrotransposition events were timed using MutationTimeR v 1.00.2 ^28^, using Illumina read counts, copy number profiles derived using Battenberg ^30^ and our pseudo-subclone structures. As with the real time WGD timing analysis using SNVs, we assumed the presence of a pseudo subclone corresponding to 30% of all biopsied tumour cells and accounting for 10% of all detected retrotranspositions. We then used phasing information from Nanopore sequencing to revise maximum likelihood timing classifications produced from MutationTimeR. Clonal retrotransposition events on one allelic copy in copy number regions with major copy number greater than one and minor copy number equal to one, were timed as clonal NA in MutationTimeR, as it is unclear whether they occurred after the gain on the major allele or at any time in the clonal period on the minor allele. We used a binomial model to measure the probability that the allele that the retrotransposition was phased to, was the major allele. If the maximum likelihood was that the retrotransposition occurred on the major allele, its timing was reclassified as clonal late. Only retrotransposition events exclusively phased to one allele and with less than 10% unphased reads were revaluated using this method.

5.5. Evaluation of timing approach

To evaluate our approach, we analysed patient PD0270 – the one with highest number of somatic retrotranspositions in our cohort – in whom we had sequenced six foci from the same primary tumour. We performed VAF estimation and then run our timing pipeline in this dataset to assess the consistency of relative timing estimation across different regions from the same tumour, namely PD0270a, PD0270a_6, PD0270a_7, PD0270a_10, PD0270a_12 and PD0270a_14. Although the results showed 12.4% (142/1,146) of the insertions had discordant timing estimations across any of the six tumour regions, we achieved an error rate <5% when assigning the most probable timing label to a particular region (**Supplementary Fig. 10f**; **Supplementary Table 15**), thus demonstrating the robustness of our approach. Of note, taking advantage of this multi-sample setup, for this tumour, a retrotransposition was classified as subclonal for further analysis if it was missing or classed as subclonal in at least 3/6 samples, otherwise it was classed as clonal early/clonal late if at least 4/6 of the samples had this timing classification for the retrotransposition. The timing was otherwise classed as clonal NA.

5.6. Timing retrotransposon-mediated genomic rearrangements

VAF estimation of retrotransposon insertions associated with rearrangements is more challenging than for canonical insertions. Here, the two breakpoints of a given rearrangement can vary in copy number, which may affect the analysis. Thus, we only considered those retrotransposon-mediated rearrangements whose breakpoints exhibited consistent timing categories. To assess the performance of our approach, we conducted a simulation involving 20 deletions, 20 duplications, 20 inversions and 10 translocations mediated by L1, all with a VAF of 0.5. While we observed a larger dispersion of the VAF estimations compared to canonical insertions, our method consistently estimated VAFs centred at 0.5, with no significant differences observed among the various types of rearrangements studied (**Supplementary Fig. 10e**). Of note, the dispersion of estimations appeared to be notably higher for duplications, a difference that may be attributable to the fact that duplications double the number of reference reads, introducing a larger variability in the inferences.

6. REFERENCES

1.	Li, H. & Durbin, R. Fast and accurate short read alignment with Burrows-Wheeler transform. *Bioinformatics* **25**, 1754-60 (2009).

2.	Li, H. *et al.* The Sequence Alignment/Map format and SAMtools. *Bioinformatics* **25**, 2078-9 (2009).

3.	Tischler, G. & Leonard, S. biobambam: tools for read pair collation based algorithms on BAM files. *Source Code for Biology and Medicine* **9**, 13 (2014).

4.	Chu, C. *et al.* Comprehensive identification of transposable element insertions using multiple sequencing technologies. *Nat Commun* **12**, 3836 (2021).

5.	Ebert, P. *et al.* Haplotype-resolved diverse human genomes and integrated analysis of structural variation. *Science* **372**(2021).

6.	Li, H. Minimap and miniasm: fast mapping and de novo assembly for noisy long sequences. *Bioinformatics* **32**, 2103-10 (2016).

7.	Ruan, J. & Li, H. Fast and accurate long-read assembly with wtdbg2. *Nat Methods* **17**, 155-158 (2020).

8.	Vaser, R., Sovic, I., Nagarajan, N. & Sikic, M. Fast and accurate de novo genome assembly from long uncorrected reads. *Genome Res* **27**, 737-746 (2017).

9.	Huang, W., Li, L., Myers, J.R. & Marth, G.T. ART: a next-generation sequencing read simulator. *Bioinformatics* **28**, 593-4 (2012).

10.	Ono, Y., Asai, K. & Hamada, M. PBSIM: PacBio reads simulator--toward accurate genome assembly. *Bioinformatics* **29**, 119-21 (2013).

11.	Consortium, I.T.P.-C.A.o.W.G. Pan-cancer analysis of whole genomes. *Nature* **578**, 82-93 (2020).

12.	Jiang, T., Liu, B., Li, J. & Wang, Y. rMETL: sensitive mobile element insertion detection with long read realignment. *Bioinformatics* **35**, 3484-3486 (2019).

13.	Zhou, W. *et al.* Identification and characterization of occult human-specific LINE-1 insertions using long-read sequencing technology. *Nucleic Acids Res* **48**, 1146-1163 (2020).

14.	Camacho, C. *et al.* BLAST+: architecture and applications. *BMC Bioinformatics* **10**, 421 (2009).

15.	Tubio, J.M.C. *et al.* Mobile DNA in cancer. Extensive transduction of nonrepetitive DNA mediated by L1 retrotransposition in cancer genomes. *Science* **345**, 1251343 (2014).

16.	Rodriguez-Martin, B. *et al.* Pan-cancer analysis of whole genomes identifies driver rearrangements promoted by LINE-1 retrotransposition. *Nat Genet* **52**, 306-319 (2020).

17.	Ostertag, E.M. & Kazazian, H.H., Jr. Twin priming: a proposed mechanism for the creation of inversions in L1 retrotransposition. *Genome Res* **11**, 2059-65 (2001).

18.	Li, H. Minimap2: pairwise alignment for nucleotide sequences. *Bioinformatics* **34**, 3094-3100 (2018).

19.	Edgar, R.C. MUSCLE: multiple sequence alignment with high accuracy and high throughput. *Nucleic Acids Res* **32**, 1792-7 (2004).

20.	Paradis, E. & Schliep, K. ape 5.0: an environment for modern phylogenetics and evolutionary analyses in R. *Bioinformatics* **35**, 526-528 (2019).

21.	Schliep, K.P. phangorn: phylogenetic analysis in R. *Bioinformatics* **27**, 592-3 (2011).

22.	Posada, D. & Crandall, K.A. MODELTEST: testing the model of DNA substitution. *Bioinformatics* **14**, 817-8 (1998).

23.	Nguyen, L.T., Schmidt, H.A., von Haeseler, A. & Minh, B.Q. IQ-TREE: a fast and effective stochastic algorithm for estimating maximum-likelihood phylogenies. *Mol Biol Evol* **32**, 268-74 (2015).

24.	Boissinot, S., Chevret, P. & Furano, A.V. L1 (LINE-1) retrotransposon evolution and amplification in recent human history. *Mol Biol Evol* **17**, 915-28 (2000).

25.	Liu, Y. *et al.* DNA methylation-calling tools for Oxford Nanopore sequencing: a survey and human epigenome-wide evaluation. *Genome Biol* **22**, 295 (2021).

26.	Bogard, N., Linder, J., Rosenberg, A.B. & Seelig, G. A Deep Neural Network for Predicting and Engineering Alternative Polyadenylation. *Cell* **178**, 91-106 e23 (2019).

27.	Abdennur, N. & Mirny, L.A. Cooler: scalable storage for Hi-C data and other genomically labeled arrays. *Bioinformatics* **36**, 311-316 (2020).

28.	Gerstung, M. *et al.* The evolutionary history of 2,658 cancers. *Nature* **578**, 122-128 (2020).

29.	Jolly, C. & Van Loo, P. Timing somatic events in the evolution of cancer. *Genome Biol* **19**, 95 (2018).

30.	Nik-Zainal, S. *et al.* The life history of 21 breast cancers. *Cell* **149**, 994-1007 (2012).

31.	Cmero, M. *et al.* Inferring structural variant cancer cell fraction. *Nat Commun* **11**, 730 (2020).

32.	Espejo Valle-Inclan, J. & Cortes-Ciriano, I. ReConPlot: an R package for the visualization and interpretation of genomic rearrangements. *Bioinformatics* **39**(2023).

7. SUPPLEMENTARY FIGURES

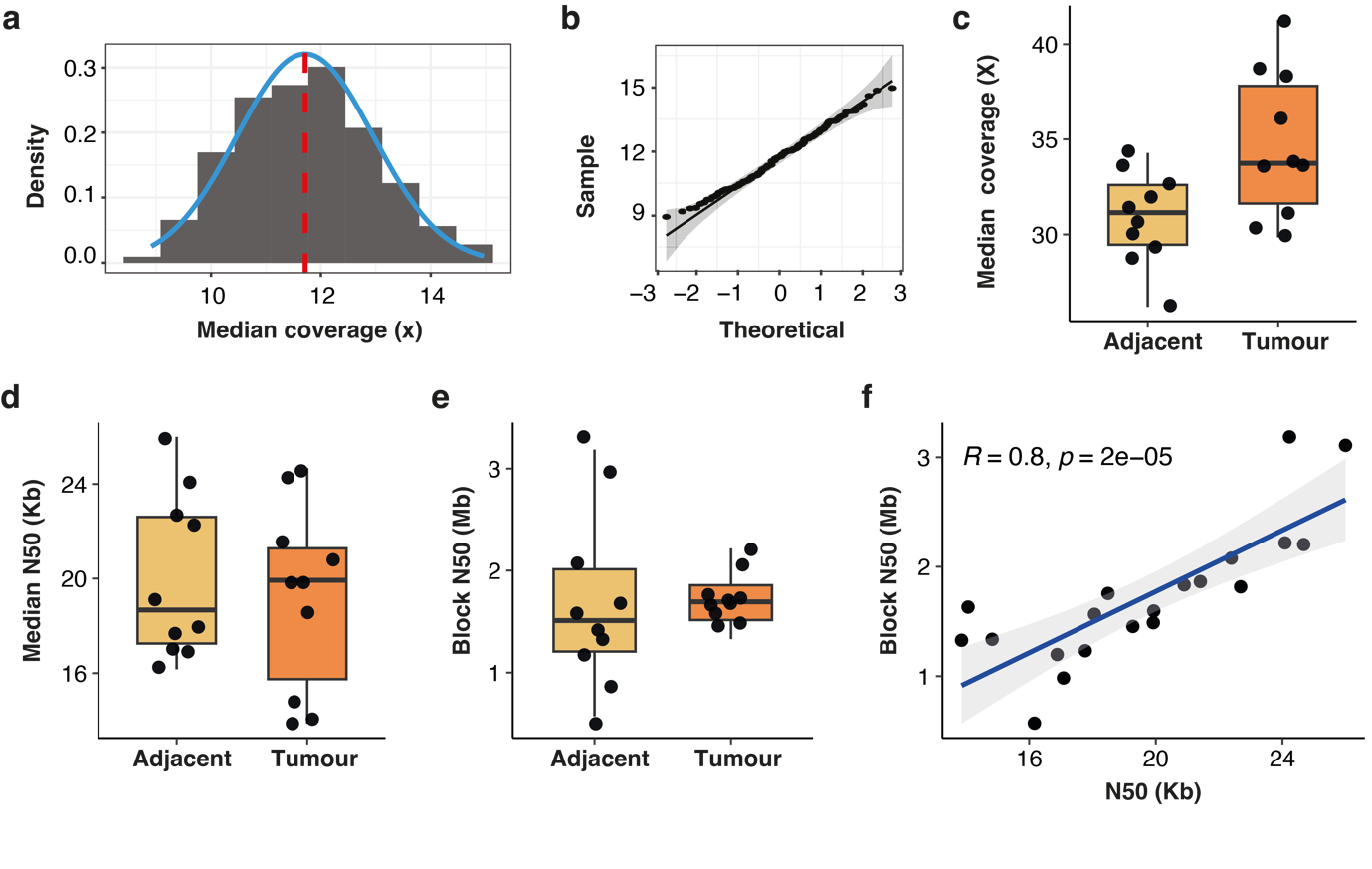

Supplementary Fig. 1. Screening, selection and long-read sequencing of 10 tumours with high rates of somatic retrotransposition.

**(a)** Distribution of the median coverage in a cohort of 137 tumours sequenced with Illumina shallow whole-genome sequencing, as part of a prospective screening. The resulting median value was 11.7x (dashed red line). **(b)** Quantile-quantile plot displaying the quantiles of the residual distribution from panel “a” on the y-axis and the theoretical quantiles from a normal distribution (x-axis). This analysis indicates that the distribution of sequencing coverage conforms to a normal distribution, suggesting an unbiased detection power across the screening cohort. **(c)** Boxplots depicting the median coverage of 10 tumours with high retrotransposition rates and their respective adjacent tissues sequenced using Oxford Nanopore Technologies (ONT). This sequencing yielded a median coverage of 33.7x for tumours and 31.1x for the adjacent tissues. **(d)** For the ONT dataset introduced in panel “c”, boxplots displaying the N50 read length metric. The observed N50 values were 19.9 kb for the tumours and 18.7 kb for the adjacent tissues. **(e)** For the ONT dataset introduced in “c”, boxplots show the N50 values of haplotype blocks derived from WhatsHap phase sets. **(f)** A significant correlation was observed between the N50 read length and the N50 phasing-blocks (Pearson correlation test, R = 0.8, *p* = 2⋅10^−5^).

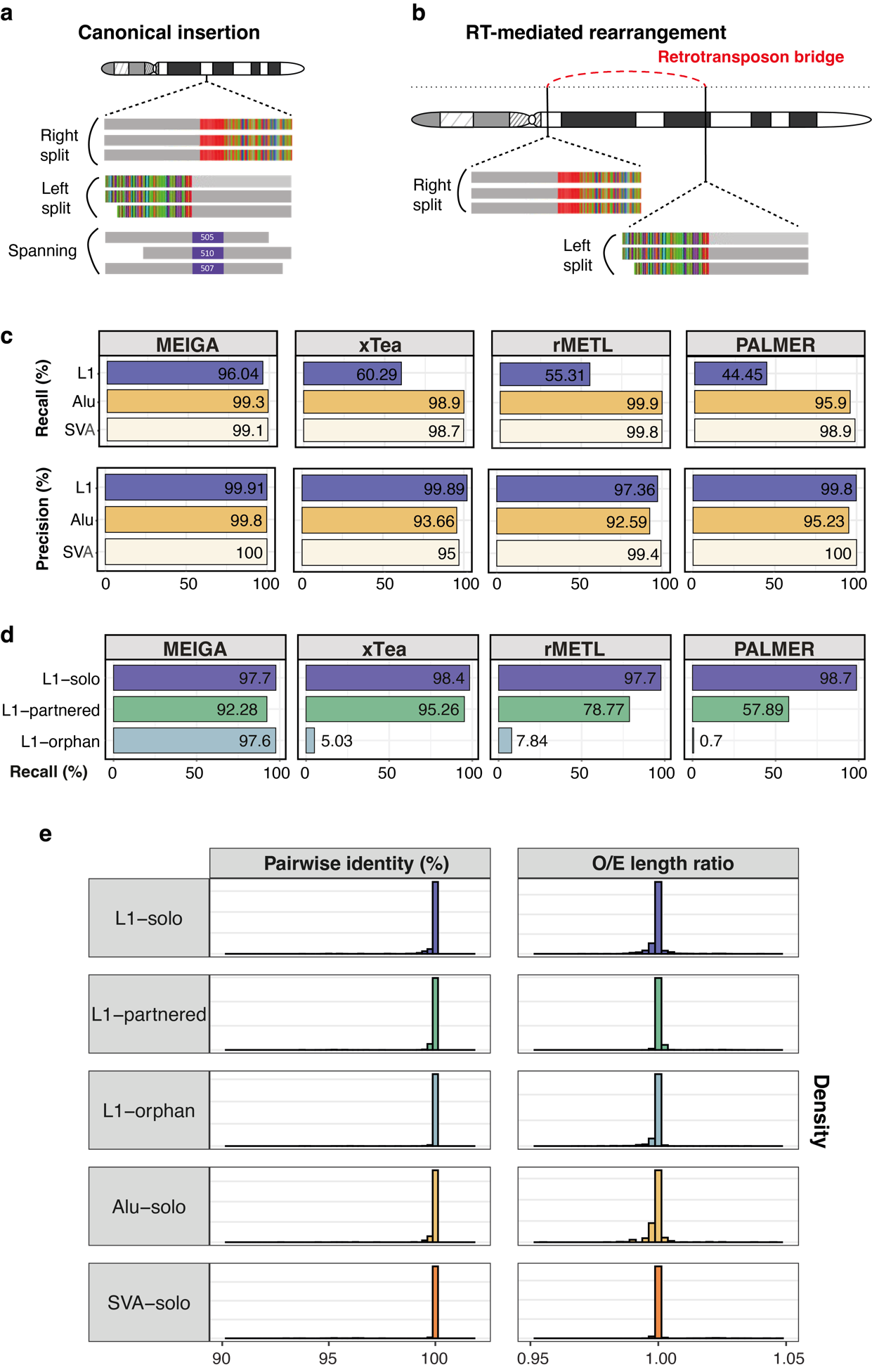

Supplementary Fig. 2. MEIGA strategy and benchmarking using simulated data.

**(a-b)** MEIGA strategy relies on the identification of clusters of split and spanning reads to find canonical retrotransposon insertions and retrotransposon bridges. **(c)** We analysed the performance of MEIGA relative to other three relevant long read pipelines, including xTea, rMETL and PALMER, to detect different types of retrotranspositions. The benchmarking dataset comprised 6,420 retrotransposon insertions representing various families and insertion types. MEIGA demonstrates high precision and recall for all types of somatic retrotranspositions. **(d)** L1 recall segregated in different L1 insertion types, showing that substantial differences arose in the ability of some algorithms to detect L1-transductions. **(e)** Evaluation of MEIGA's capability to reconstruct retrotransposed sequences, showing an average pairwise identity percentage of 99.9% and a mean observed-to-expected length ratio of 0.99.

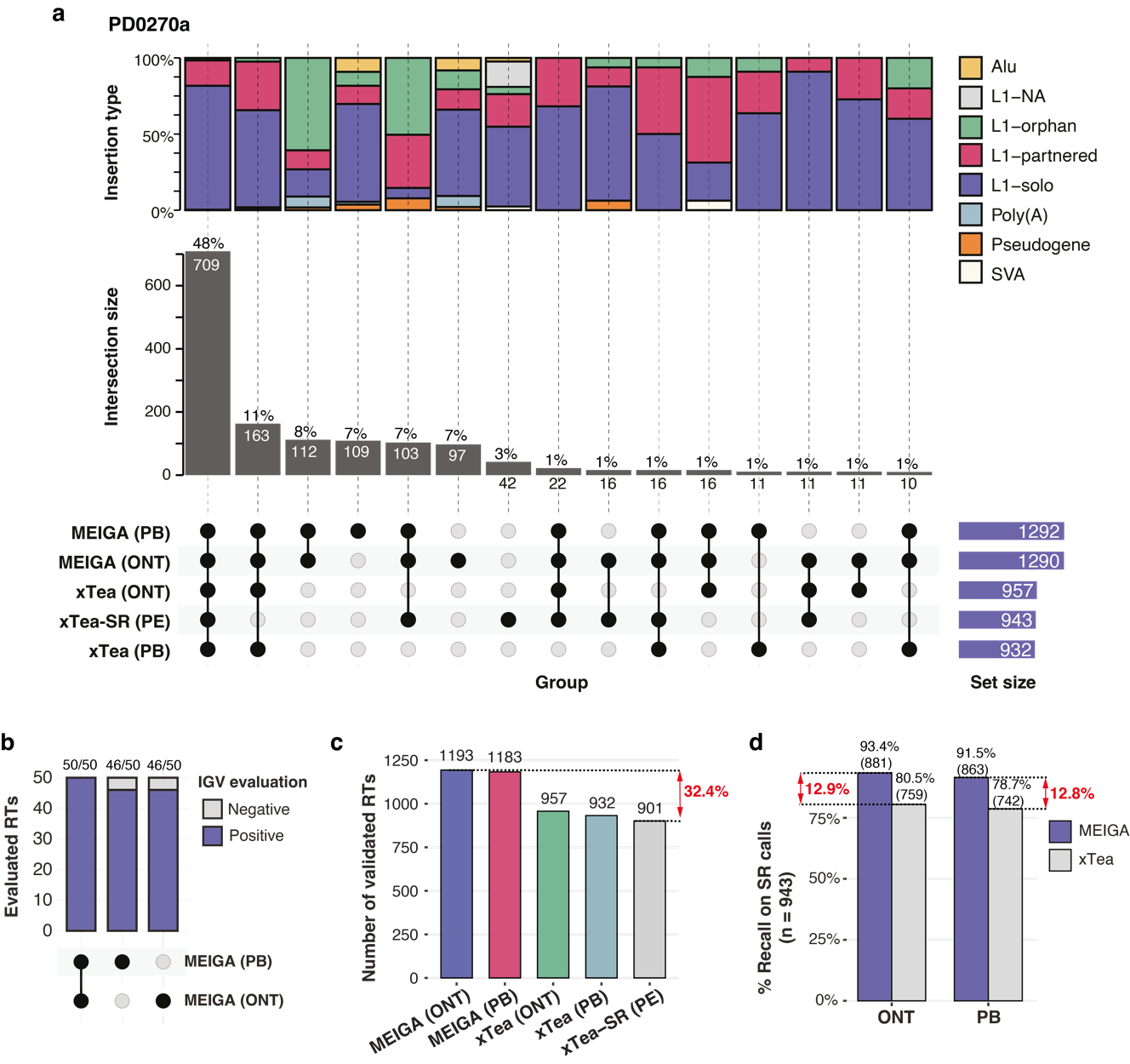

Supplementary Fig. 3. MEIGA benchmarking using real data from donor PD0270a.

In-silico validation of canonical somatic retrotransposition events from tumour PD0270a using different sequencing datasets, including Oxford Nanopore (ONT), PacBio HIFI (PB) and Illumina paired-ends (PE), and different pipelines, including MEIGA, xTea and the xTea module for short reads (xTea-SR). **(a)** Comparative analysis of the detection of somatic retrotransposition insertions by three reference pipelines across various sequencing technologies. The analysis encompasses five categories: MEIGA ONT, MEIGA PB, xTea ONT, xTea PB and xTea-SR PE. The intersection of insertion events identified by each method is depicted. In the upset plot, each intersection group is represented in the panel below, with absolute counts indicated in the central panel and insertion types specified in the upper panel. Of note, since xTea does not support somatic calling, xTea private calls were excluded from the analysis because most corresponded to germline variants. This analysis shows that MEIGA yielded the highest somatic retrotransposition rates, with 1,290 events in the ONT data and 1,292 in the PB data. Notably, approximately 93% of these events, for both the ONT and PB datasets, were detected by at least one alternative method or sequencing technology, thereby pointing to genuine, validated events. **(b)** To evaluate the group of MEIGA-private calls (97 calls on the ONT dataset, 109 calls on the PB dataset and 112 calls detected on both ONT and PB), we carried out visual inspection with Integrative Genomics Viewer (IGV) of a random selection of 150 insertions (50 per group), which revealed a true positive rate of 94.7%. (142/150). **(c)** The analysis with long reads and MEIGA reveals approximately 32.4% more somatic insertion events than with short reads. Of note, here we only represent those somatic events validated by at least one alternative method or sequencing technology as a proxy for genuine events. **(d)** To assess the sensitivity of both long-read methods, MEIGA and xTea, we computed the fraction of somatic events they identified from the dataset of 943 somatic calls detected with short reads in this setup. We observe that MEIGA can call ~12.9% and 12.8% more somatic events than xTea on the ONT and PacBio datasets, respectively.

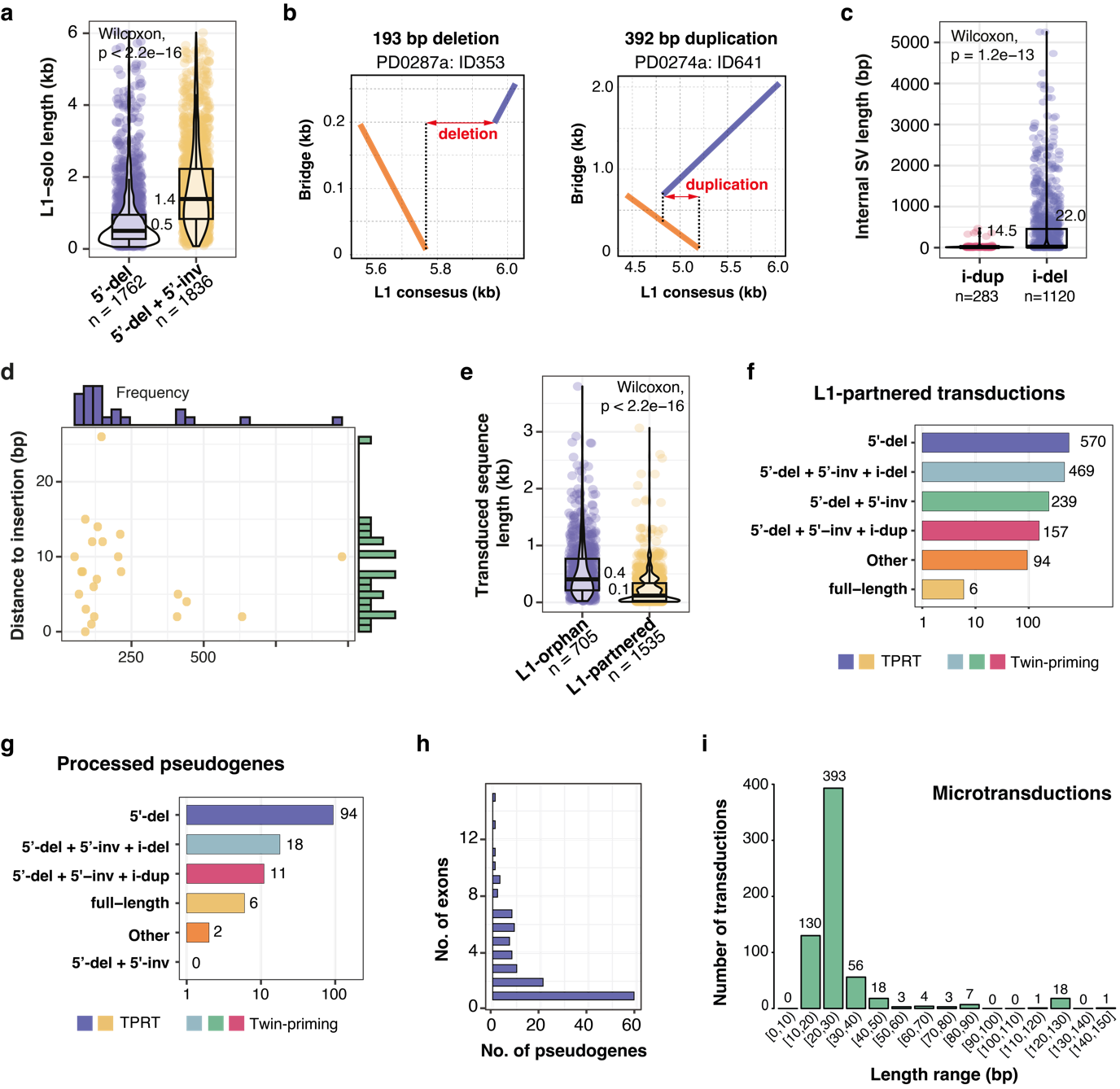

Supplementary Fig. 4. Structural analysis of somatic retrotransposition events.

**(a)** Box plots showing the insertion length distribution of 3,598 non-functional copies of L1 retrotransposition events segregated into two categories: those presenting 5’ deletion only (n = 1,762) and those presenting co-occurrence of 5’ deletion and 5’ inversion (n = 1,836), with median size at 0.5 kb and 1.4 kb, respectively. **(b)** Dot plots showing the patterns of internal deletion (left) and internal duplication (right) in two representative insertions from our cohort. The dot-plots represent the comparison of the sequence from the L1 insertion reconstructed with ONT (Y-axis) and the L1 canonical insertion (X-axis); the orange line represents the inversion generated by twin priming that matches the cDNA growing from the internal primer, while the blue line matches the cDNA extended from the poly(dT) primer. **(c)** Box plots showing the length distribution of internal deletions (n = 1,120) and internal duplications (n = 283) generated by twin priming in our cohort, with median size at 22 bp and 14.5. bp, respectively. **(d)** Distribution of templated insertions according to their length (X-axis) and the distance in the reference genome of the copied template to the insertion site, showing most templates are below 250 bp and less than 20 bp from the insertions site. **(e)** Box plots showing the length distribution of transduced nucleotide sequences found in L1 orphan (n = 705) and L1 partnered (n = 1,535) transductions, with median size at 0.4 kb and 0.1 kb, respectively. **(f)** Barplot showing the number and structural types of somatic L1-partnered transductions observed in the 10 tumours cohort. Events are classified into six types of structures: full-length by canonical TPRT; 5’-deletion only by TPRT; co-occurrence of 5’-deletion and 5’-inversion inversion by twin priming; co-occurrence of 5’-deletion, 5’-inversion and internal deletion (i-del) by twin priming; cooccurrence of 5’-deletion, 5’-inversion and internal duplication (i-dup) by twin priming; co-occurrence of 5’-deletion and other minor structures. **(g)** Barplot showing the number and structural types of somatic processed pseudogenes observed in the 10 tumours cohort. Events are classified into six structural types: full-length by canonical TPRT; 5’-deletion only by TPRT; co-occurrence of 5’-deletion and 5’-inversion inversion by twin priming; co-occurrence of 5’-deletion, 5’-inversion and internal deletion (i-del) by twin priming; cooccurrence of 5’-deletion, 5’-inversion and internal duplication by twin priming; co-occurrence of 5’-deletion and other minor structures. **(h)** Distribution of the number of pseudogenes according to the number of exons that they mobilize. **(i)** Distribution of the lengths of L1-partnered transductions in our cohort. Only transductions below 150 bp are shown, which includes microtransductions (i.e., those below 50 bp)

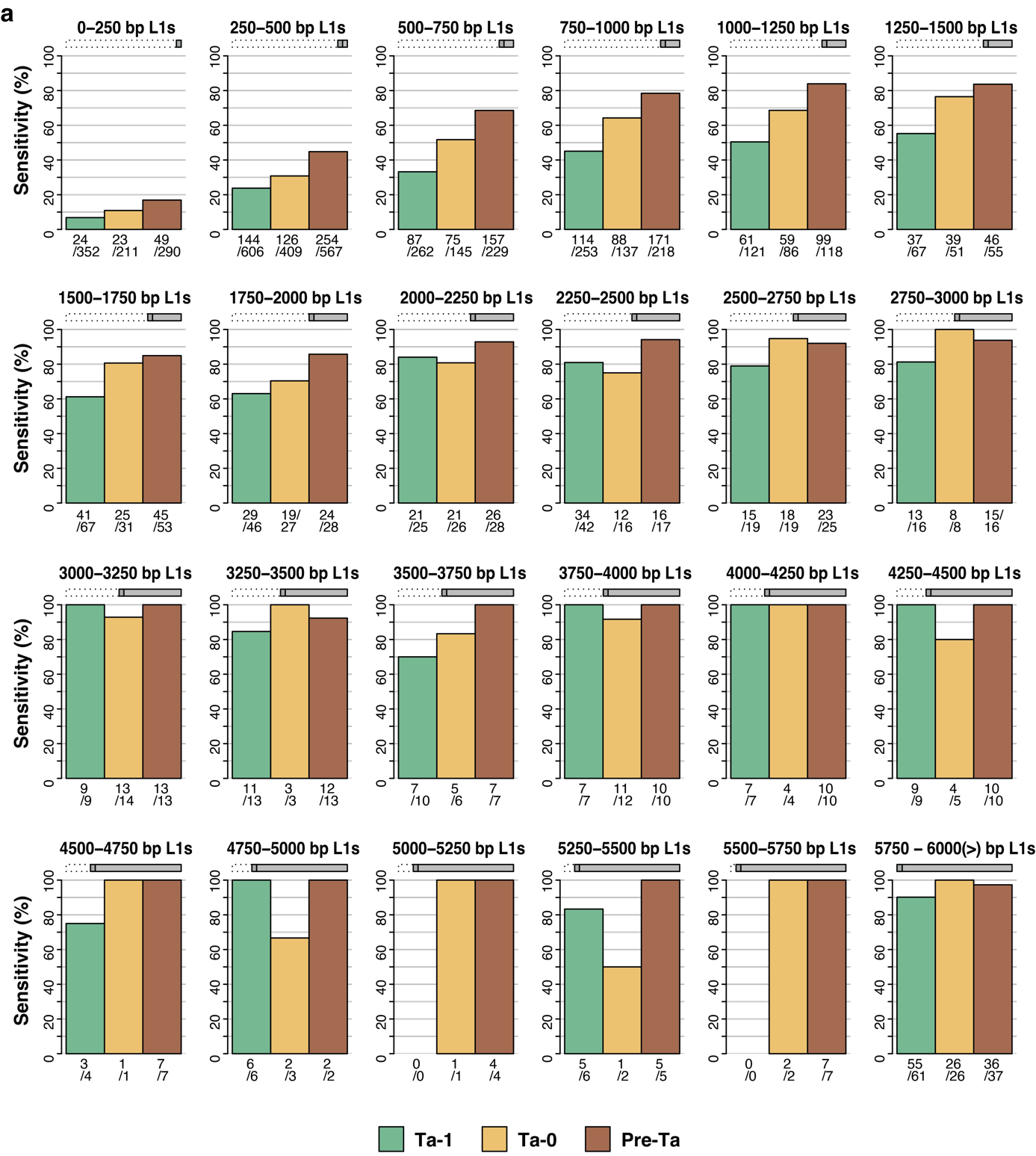

Supplementary Fig. 5. Evaluation of the Source Inference method.

The sensitivity of Source Inference was assessed by testing its ability to trace solo-L1 insertions to its original L1 source element in a mock cancer genome generated with MEIsimulator. We simulated the mobilization of all potential source elements (n = 266) from the patient in the cohort with the highest L1 activity rate (PD0270), assuming a homogeneous activity of 20 retrotranspositions for each one of the 266 elements and with the derived copies showing the same insertion size distribution as in 2954 cancer genomes from the PCAWG project ^16^. Vertical bars indicate the fraction of events traced to its element-of-origin (true positive rate) under each specific insertion size range (5’ truncation interval decreasing in 250 bp on each plot, as represented in the top L1 ideograms). The number of detected and simulated events for each category is indicated under each vertical bar. Bar colours indicate L1Hs subfamily: Ta-1 (green); Ta-0 (orange); pre-Ta (brown).

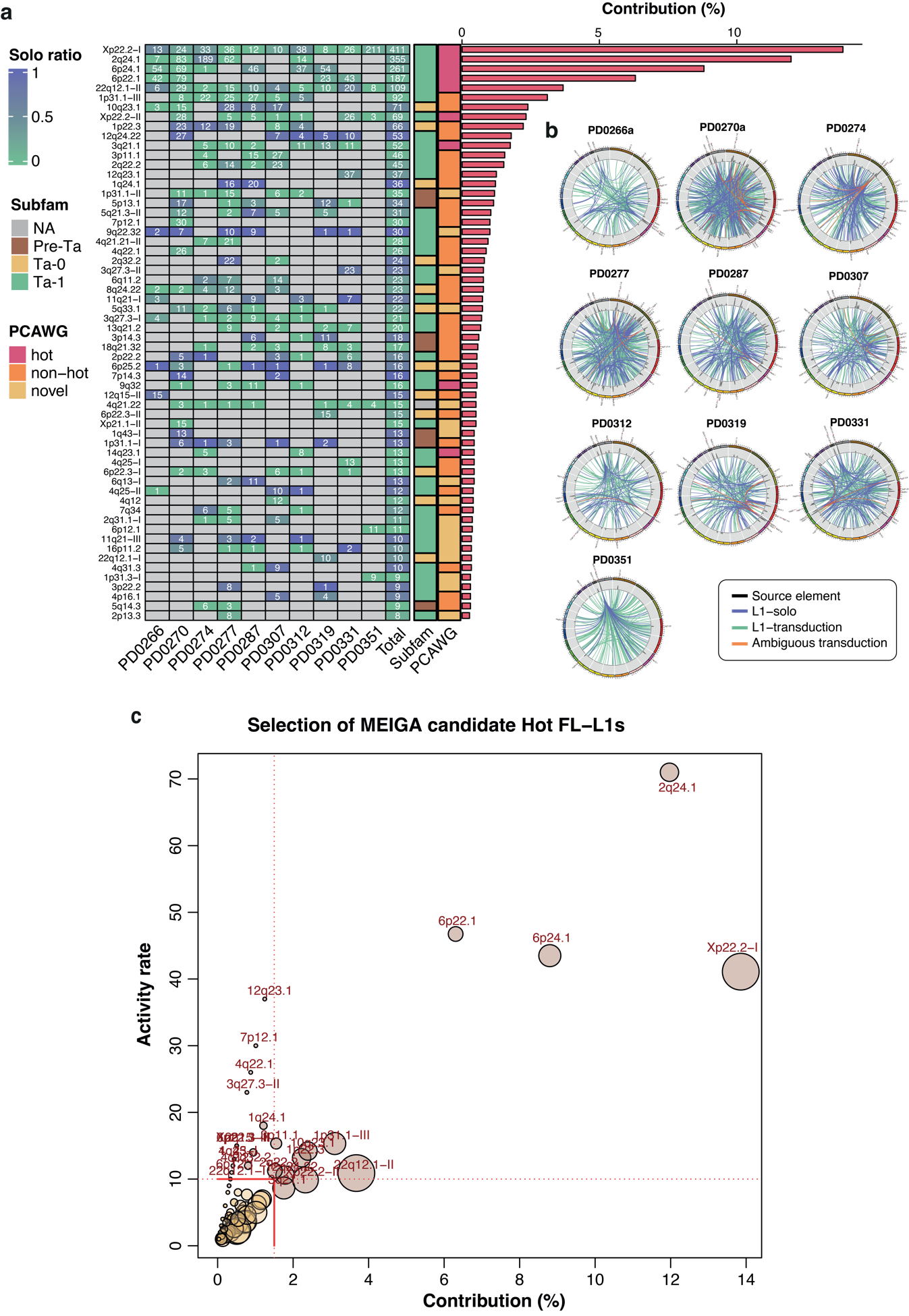

Supplementary Fig. 6. Landscape of source element activity in a cohort of 10 tumours.

**(a)** Solo-ratio of L1 source elements in the 10 tumours cohort. Heatmap showing the total number of somatic mobilizations traced to each active L1 source element (rows) on each sequenced tumour (columns). The heatmap cell colour ranges according to the ratio between activity entirely derived from solo-insertions (purple) and activity entirely derived from transductions (green). The right annotation track indicates: L1Hs subfamily (green: Ta-1, orange: Ta-0, brown: pre-Ta); PCAWG somatic activity (red: PCAWG hot source element, orange: PCAWG non-hot source element, yellow: novel activity); Contribution: L1 source elements are sorted according to their relative contribution (%) to the total number of traced mobilizations in the cohort. Note that only the top 60 source elements showing higher number of derived copies in the cohort are included (the whole dataset can be extracted from **Supplementary Table 4**). **(b)** For the 10 tumours in the cohort, Circos plots show the mobilization of somatic copies from the corresponding L1 source element, including canonical transductions (green links), ambiguous transductions (orange links; these are typically very short L1-partnered transduction whose element-of-origin could only be identified by diagnostic nucleotides present in the companion L1 sequence) and solo-L1 insertions (purple links). The labels indicate the cytogenetic position of each source element. **(c)** Identification of hot L1s. Scatter plot showing the distribution of L1 source elements according to the following two variables: L1 activity rate (y axis) and relative contribution to traced somatic mobilizations (x-axis). Each circle corresponds to an active L1 source element, coloured according to a density-based clustering identity. Circle size is proportional to the number of patients in which the source element is present (1-10). Red lines mark the threshold values applied for hot element selection (L1 activity rate ≥10 and / or contribution ≥1.5%). Hot L1 elements meeting these criteria are denoted with GRCh38 cytoband identifiers.

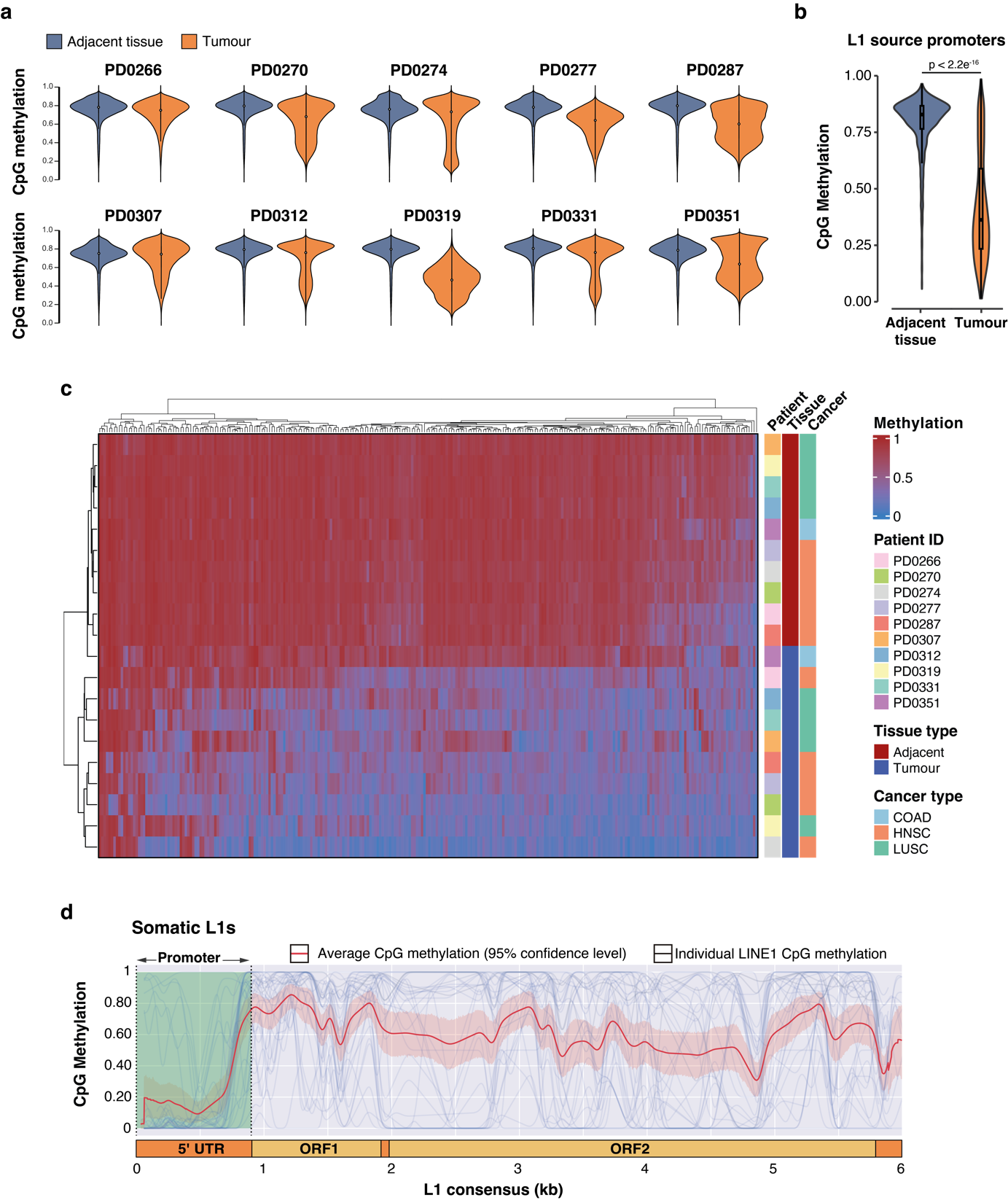

Supplementary Fig. 7. ONT sequencing analysis of DNA methylation.

**(a)** Whole-genome CpG methylation. Violin plots depicting the whole-genome CpG methylation status across the 10 primary tumour samples (blue) and their corresponding matched adjacent tissues (orange) analysed in this study. **(b)** Violin plots showing the distribution of CpG methylation levels across 269 full-length L1 loci shared by all 10 patients in both tumour and matched adjacent normal tissues. **(c)** Building on panel b, heatmap depicting the CpG methylation status across the L1 promoters of the 269 candidate source element loci shared by all patients in the studied cohort with annotations and hierarchical clustering of columns (L1 loci) and rows (Patients) according to their methylation patterns based on Euclidean distances. **(d)** Promoter methylation patterns in somatic L1 insertions. Patterns of CpG methylation along the consensus L1 sequence for 71 somatically acquired L1 insertions that retain all or part of their promoter region.

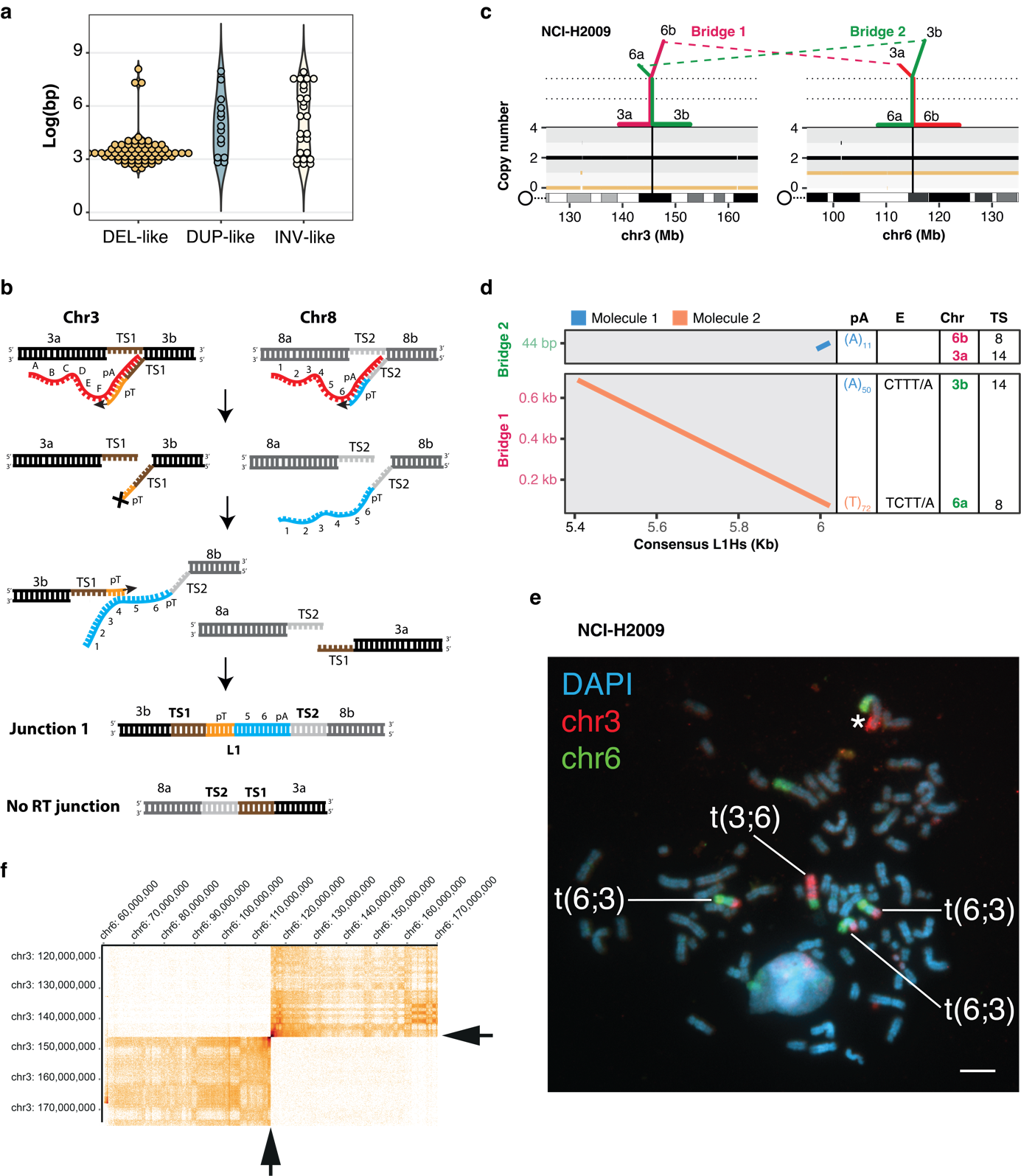

Supplementary Fig. 8. Retrotransposition-mediated rearrangements.

**(a)** Size distribution of retrotransposon-mediated rearrangements according to the categories deletion-like, duplication-like and inversion-like. **(b)** Mechanisms proposed for the reciprocal translocation between chromosomes 3p and 8q found in tumour PD0351a. The cDNAs from two incipient insertion at chromosomes 3 and 8 generated by canonical TPRT, anneal by complementarity to promote crossing over between the poly(T) cDNA of one event and the L1 cDNA of the other. Consequently, this process forms a bridge consisting of an L1 sequence flanked by two poly(A/T) tails. **(c-f)** Validation by FISH of a L1-mediated reciprocal translocation between chromosomes 3q and 6q found in the high retrotransposition rate cell line NCI-H2009: **(c)** ReConPlot ^32^ showing the two bridges and chromosomal configuration of the reciprocal translocation, where bridge 1 joins chr3a and chr6b, while bridge 2 joins chr3b and chr6a; **(d)** For the translocation shown in “d”: Dot plots revealing the internal configuration of two independent L1 retrotranspositions; **(e)** FISH results validating the reciprocal translocation between chromosomes 3 and 6, with the expected configuration for both derivatives; chromosome 3 is labelled in red and chromosome 6 is labelled in green. **(f)** Micro-C contact map demonstrates de novo chromatin interactions between chromosomes 3 and 6 at the expected breakpoints of the reciprocal translocation. Arrows indicate crossover breakpoints.

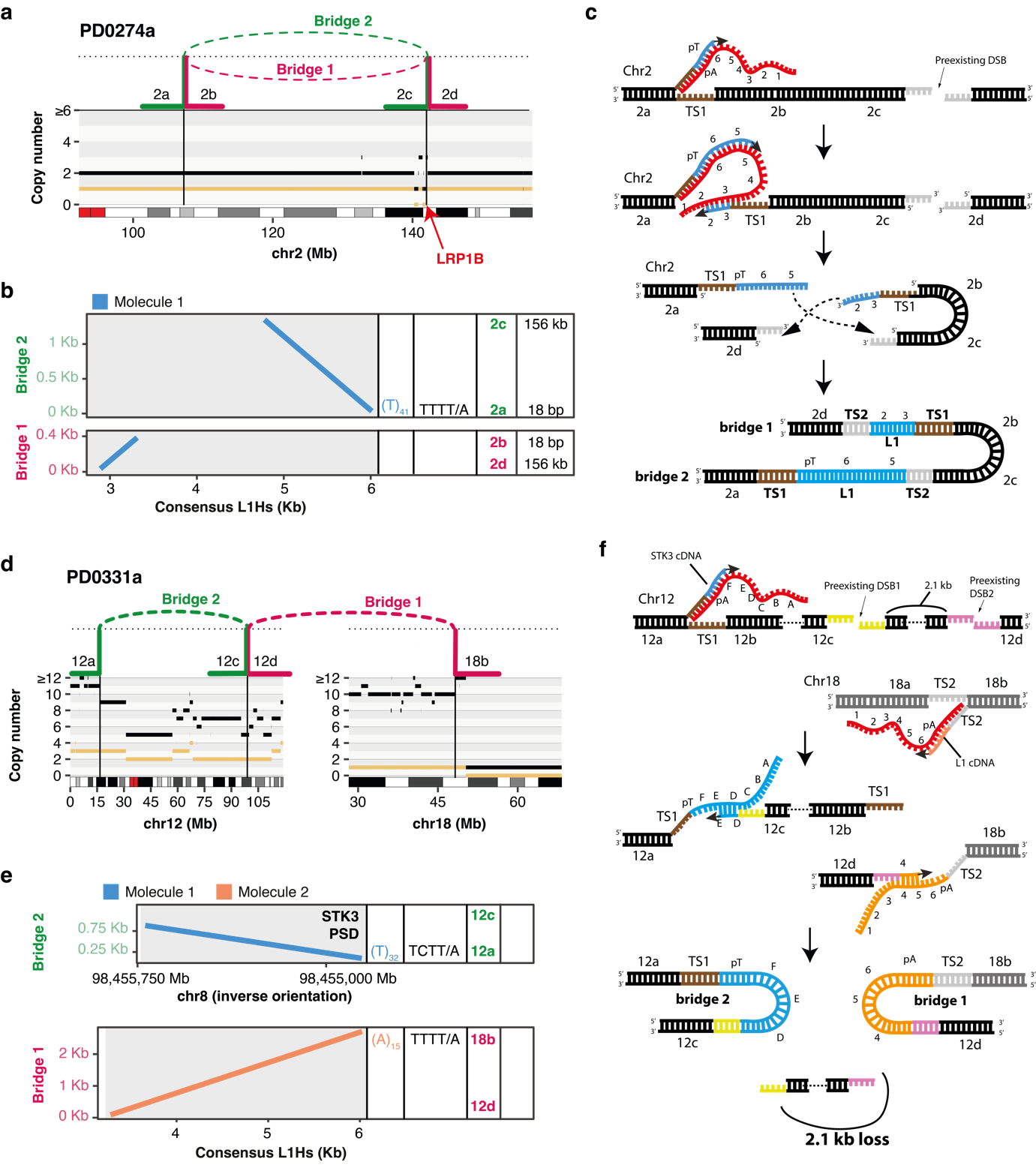

Supplementary Fig. 9. Reciprocal inversions and complex rearrangements mediated by retrotransposition.

**(a)** In tumour PD0274a, ReConPlot showing the bridges and chromosomal configuration of a reciprocal paracentric inversion mediated by two retrotransposition bridges in chromosome 2q affecting the tumour suppressor gene *LRBP1*. **(b)** For the inversion shown in “a”: Dot plots representing the pairwise similarity between the nucleotide sequence of the L1 bridges (Y-axis) and the consensus L1 sequence. Here, a poly(T) tail together with a TTTT/A endonuclease motif are found at the 5’ end of the second L1 bridge and a single L1 internal sequence is found conforming the first bridge. The patterns indicate the involvement of one-single retrotransposition event forming both bridges. **(c)** Mechanism for the reciprocal inversion at chromosome 2q that explains the configuration of the bridges in “b” and the genomic rearrangement in “a”, requiring an L1 insertion following twin priming and the presence of a pre-existing double-strand break (DSB). **(d)** Complex rearrangement with two L1 bridges involving an inversion-like rearrangement at chromosome 12 and an interchromosomal rearrangement between chromosomes 12q and 18q. Here, bridge 1 joins chr12d and chr18b, while bridge 2 joins chr12a and chr12c. Of note, there is a small deletion 2.1 kb-long between breakpoints chr12c and chr12d, which is not shown due to the resolution of the plot. **(e)** For the complex rearrangements in “d”, the dot plot reveals the involvement of two different L1 insertion events. A poly(A) tail together with an endonuclease motif (TTTT/A) are found at the 5’ end of bridge 1, while a poly(T) tails together with the endonuclease motif TCTT/A are found at the 5’ end of the second bridge. **(f)** Mechanism that explains the configuration of the bridges in “e” and the genomic rearrangements in “d”, requiring two independent L1 insertions in distinct chromosomes. Both events are used to repair the extremes from two close double-strand breaks at the 12q separated by 2.1 kb. The final result is the formation of an inversion-like rearrangement between chr12a and chr12c, an interchromosomal rearrangement between chr12d and chr18b, and the loss of 2.1 kb between chr12c and chr12d.

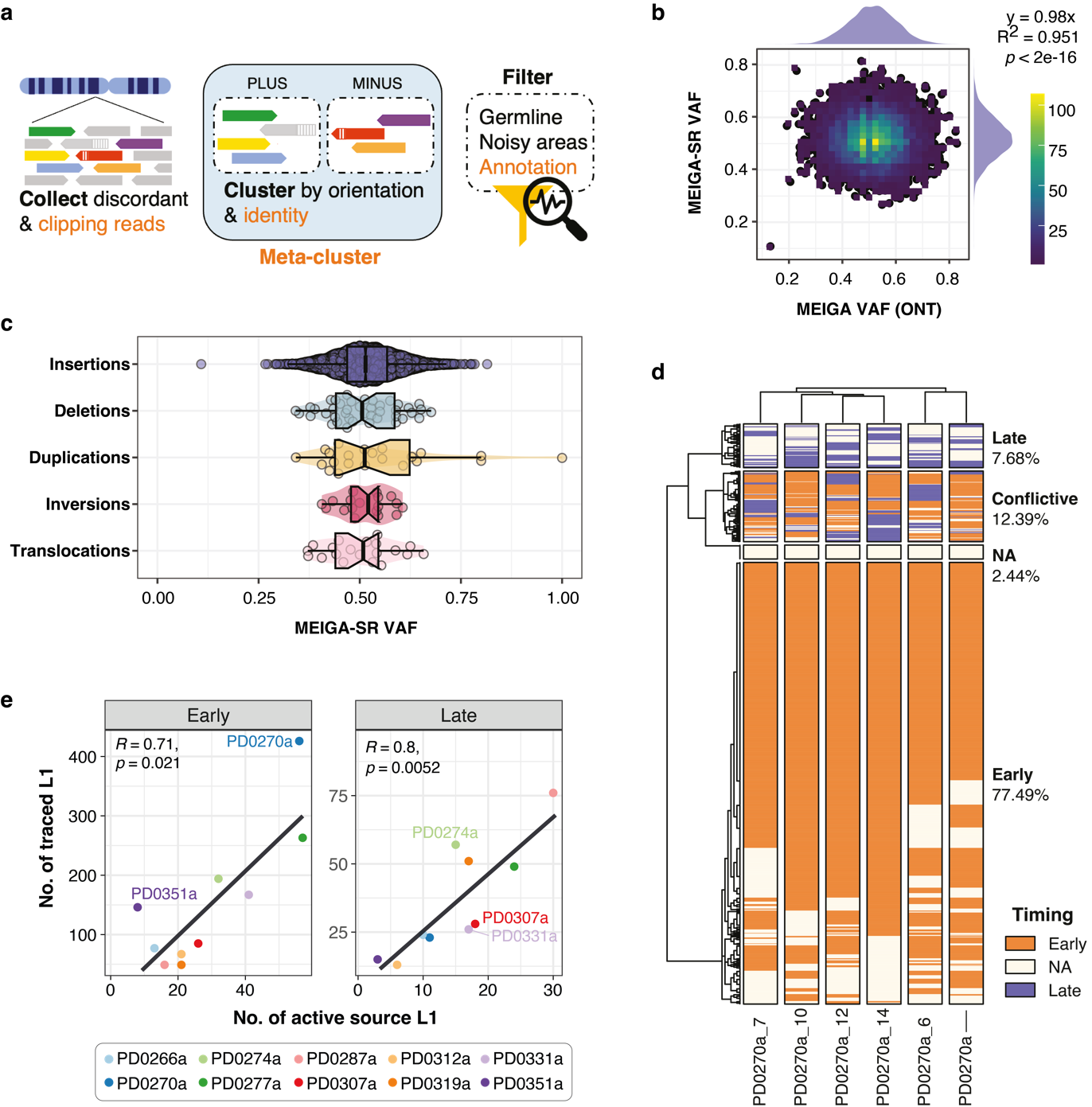

Supplementary Fig. 10. MEIGA-SR and timing of somatic retrotranspositions.

**(a)** MEIGA-SR incorporates four key features, highlighted in orange, to enhance sensitivity over conventional approaches typically used by other retrotransposition callers. Firstly, it identifies RT insertions solely supported by clipped reads, often linked with shorter insertions. Secondly, MEIGA-SR recognises discordant reads that support poly(A/T) repeats, thereby improving overall read support and facilitating detection of RT insertions with extended poly(A/T) tails or those comprised entirely of poly(A/T) tracks. Thirdly, it requires minimal support for primary clusters (i.e., two reads), aiding in identifying events where only one primary cluster in a metacluster receives robust support. Fourthly, relaxing the milli-divergence threshold and incorporating criteria such as poly(A/T) and TSD detection aids in the identification of nested insertions. **(b)** In a benchmarking analysis with simulated data (6,420 retrotransposon insertions), we evaluated the accuracy of MEIGA-SR in determining the VAF of somatic retrotransposition events for timing purposes. Concomitantly, a long-read sequencing dataset with an identical set of simulated events (n = 6,420), each at a 0.5 VAF, was analysed using our long reads method, MEIGA. The results showed that the estimated VAF distributions for both MEIGA-SR and MEIGA (long reads) were closely aligned with the expected value of 0.5. Furthermore, the distributions fitted a normal distribution and exhibited a strong linear correlation (y = 0.98x; R² = 0.951), indicating no significant bias in either method. **(c)** For simulated retrotransposition-mediated rearrangements at a VAF of 0.5, MEIGA-SR consistently estimated VAFs centred around 0.5, with no significant differences across the various rearrangement types analysed (Kruskal-Wallis rank sum test, *p*>0.05). However, the dispersion of VAF estimations was notably higher for duplications, which could be attributed to the fact that duplications double the number of reference reads, introducing greater variability in the inferences (Standard deviations: duplications = 0.157, deletions = 0.088, inversions = 0.061, translocations = 0.086, insertions = 0.074). **(d)** Consistency of relative time timing estimation across multi-region samples in tumour PD0270a. Heatmap showing the relative timing estimates across multiple samples from donor PD0270a. Gower’s distance was used to cluster the samples and the timing labels. We identified a conflict when the same retrotransposon insertion was labelled as clonal early and clonal late, or as clonal early and subclonal in different samples from the same tumour. The results revealed that conflicting timing labels were identified in 12.39% of cases for at least one of the six sequenced regions. Nevertheless, we achieved an accuracy rate of 97.94% when evaluating our ability to correctly assign the most probable timing label to an individual region, thus demonstrating the robustness of our methodology. Timing labels ‘Clonal Late’ and ‘Subclonal’ were grouped as ‘Late,’ while ‘Clonal Early’ events were categorized as ‘Early.’ Any instances where ‘Early’ and ‘Late’ labels were assigned to the same event in different samples were considered as ‘Conflictive’. NA refers to clonal Not Assigned events. **(e)** Correlation between the number of active source elements and the total number of source-traced events for both early (left; Pearson correlation test, R = 0.82, *p* = 0.0036) and late stages (right; Pearson correlation test, R = 0.84, *p* = 0.0022). The number of active source elements was estimated using both transductions and solo L1s.
